# LBD-type transcription factors suppress local and systemic nitrogen responses through distinct regulatory pathways

**DOI:** 10.64898/2026.08.14.744662

**Authors:** Takatoshi Kiba, Hana Takahashi, Kota Monden, Yukino Sada, Keiichi Koshihara, Muneo Sato, Fanny Bellegarde, Takushi Hachiya, Masami Yokota Hirai, Shuichi Yanagisawa, Hitoshi Sakakibara

## Abstract

Nitrogen (N) is a major determinant of plant growth and productivity. Because soil N availability and internal N demand fluctuate, plants have evolved sophisticated mechanisms to coordinate N acquisition and utilization at the whole-plant level. However, how this coordination is achieved remains poorly understood. Here, we show that N-inducible LATERAL ORGAN BOUNDARIES DOMAIN transcription factors LBD37, LBD38, and LBD39 (LBDs) function as repressors of local N uptake and assimilation and systemic N-demand signaling in *Arabidopsis*. Triple mutants lacking these three LBDs displayed enhanced nitrate influx and increased accumulation of nitrate, amino acids, and total N. Transcriptome analysis identified an array of N-starvation- and nitrate-inducible genes derepressed in shoots and roots, including *C-TERMINALLY ENCODED PEPTIDE* (*CEP*) and *CEP DOWNSTREAM* (*CEPD*) genes, as well as genes involved in N uptake and assimilation. Grafting and genetic analyses revealed that LBDs gate the systemic N-demand signaling relay by repressing *CEP* and *CEPD* expression organ-autonomously. We also found that LBDs locally repress genes involved in N uptake and assimilation through a distinct regulatory mechanism. We propose that LBDs are key transcriptional repressors in a regulatory framework for optimizing N acquisition and utilization under fluctuating N conditions at the whole-plant level.

## INTRODUCTION

Nitrogen (N) is a major factor determining plant growth and productivity. Under natural conditions, soil N availability can vary markedly across space and time, owing to factors such as leaching, microbial activity, and local root uptake (Miller et al. 2007; Yeshno et al. 2019). Similarly, internal N demand also changes with growth and development. Plants have therefore evolved mechanisms that coordinate N acquisition and utilization with both fluctuating external N availability and changing internal N status (Forde 2002; Good et al. 2004).

Studies in *Arabidopsis* have revealed two major responses that contribute to this coordination: nitrate responses and N-starvation responses (collectively referred to as N responses hereafter). The nitrate response is activated when nitrate, a major N source in aerobic soils, is supplied to N-depleted plants. This response encompasses a wide range of physiological and developmental processes, including nitrate acquisition and assimilation (Krapp 2015), germination (Alboresi et al. 2005), shoot and root growth and development (Wang et al. 2018; Abualia et al. 2022), transpiration (Guo et al. 2003), and flowering (Sanagi et al. 2021). The nitrate response is underpinned by extensive transcriptome reprogramming, including the induction of genes involved in nitrate uptake, reduction, and assimilation, such as *NITRATE TRANSPORTER2.1* (*NRT2.1*) and *NITRATE REDUCTASE1* (*NIA1*) (Wang et al. 2003; Scheible et al. 2004). NIN-LIKE PROTEINs (NLPs) are master regulators of this nitrate-responsive transcriptional program. In response to nitrate, NLPs are post-translationally activated and directly induce primary nitrate-responsive genes (Konishi and Yanagisawa 2013; Marchive et al. 2013; Liu et al. 2017, 2022; Konishi et al. 2021; Durand et al. 2025).

N-starvation responses are induced when external N supply is limited relative to plant demand. These responses are characterized by enhanced high-affinity nitrate uptake, remodeling of root system architecture, and promotion of N recycling and remobilization (Krapp et al. 2014; Kiba and Krapp 2016). In *Arabidopsis*, *NRT2* family transporter genes, including *NRT2.1*, *NRT2.4*, and *NRT2.5*, are representative N-starvation-inducible genes responsible for enhanced high-affinity nitrate uptake (Kiba et al. 2012; Lezhneva et al. 2014). Both local and systemic signaling pathways have been implicated in the regulation of N-starvation responses (Lejay et al. 1999; Gansel et al. 2001; Ruffel et al. 2011), and several regulatory factors have been identified in *Arabidopsis*. Locally, CALCINEURIN B-LIKE PROTEIN7 (CBL7) acts as a positive regulator of *NRT2.4* and *NRT2.5* expression and root architecture under N starvation (Ma et al. 2015). The microRNA miR169, which is downregulated under N starvation, has been reported to negatively regulate *NRT2.1* expression (Zhao et al. 2011). In systemic regulation, C-TERMINALLY ENCODED PEPTIDE (CEP) and CEP DOWNSTREAM (CEPD) family members function as root-to-shoot and shoot-to-root mobile signals, respectively, and together constitute a systemic N-demand signaling system. *CEP* genes are expressed in N-starved roots, and mature CEPs move to shoots via the xylem. Upon CEP perception in shoots, *CEPD1* and *CEPD2* are induced, and the encoded proteins move from shoots to roots via the phloem to promote nitrate uptake by activating nitrate-uptake-related genes, including *NRT2.1, NRT3.1*/*NAR2.1* (*NAR2.1*), and *CEPD-INDUCED PHOSPHATASE* (*CEPH*) (Tabata et al. 2014; Ohkubo et al. 2017, 2021; Ota et al. 2020). *CEPDL2* is induced by shoot N starvation, and its encoded protein moves and functions similarly to CEPD1 and CEPD2 (Ota et al. 2020). Recent studies have further shown that the action of CEPDs is mediated through TGACG-BINDING FACTOR1 (TGA1) and TGA4 transcription factors (Kobayashi et al. 2024; Thurow et al. 2025). More recently, TGA7 was identified as a shoot-to-root mobile transcription factor that promotes root growth and nitrate uptake under N starvation by directly activating *NRT2.1* and regulating other nitrate-uptake-related genes through a NUCLEAR FACTOR YA10 (NF-YA10)-mediated transcriptional cascade (Ye et al. 2025).

Although nitrate responses and N-starvation responses are triggered by different N-related cues, both are subject to negative regulation when N availability is high (Okamoto et al. 2003; Nazoa et al. 2003; Kiba et al. 2012). Nitrate-responsive gene expression is attenuated over time following its induction by nitrate and is repressed by reduced N sources, such as ammonium and amino acids. N-starvation-inducible genes are also repressed by supplementation with nitrate or reduced N sources. The NITRATE-INDUCIBLE GARP-TYPE TRANSCRIPTIONAL REPRESSOR1/HYPERSENSITIVE TO LOW Pi-ELICITED PRIMARY ROOT SHORTENING1 family transcriptional repressors (NIGT1s), which comprise four members in *Arabidopsis*, have been shown to play a role in this negative regulation. *NIGT1* genes are rapidly induced by nitrate downstream of NLPs and NIGT1 proteins directly repress a subset of nitrate-inducible genes, including *NRT2.1* (Maeda et al. 2018). This NIGT1-mediated repression counteracts NLP-dependent activation, dampening nitrate-induced gene expression over time (Maeda et al. 2018). NIGT1s also directly repress N-starvation-inducible genes, including *NRT2.4* and *NRT2.5*, in response to nitrate (Kiba et al. 2018; Safi et al. 2021). Moreover, *NIGT1.1* and *NIGT1.2* are induced by reduced N sources in a dose-dependent manner (Kiba et al. 2018). Thus, NIGT1-mediated repression helps adjust nitrate responses and N-starvation responses according to external N availability and internal N status. However, loss of NIGT1 activity results in only partial derepression of these N-regulated genes, suggesting that the mechanisms underlying this negative regulation are not yet fully understood (Kiba et al. 2018; Maeda et al. 2018; Safi et al. 2021).

LATERAL ORGAN BOUNDARIES DOMAIN proteins constitute a plant-specific family of transcription factors characterized by a conserved LOB domain (Iwakawa et al. 2002; Shuai et al. 2002). Among them, LBD37, LBD38, and LBD39 (LBDs) are closely related members encoded by genes initially identified through microarray analyses as nitrate- and reduced-N-inducible genes (Scheible et al. 2004; Rubin et al. 2009). Overexpression analyses suggested that these LBDs negatively regulate N-responsive genes involved in nitrate uptake and assimilation, including *NRT2.1*, *NRT2.5*, *NIA1*, and *NIA2*, and affect N metabolism (Rubin et al. 2009; Albinsky et al. 2010). Gene regulatory network analyses placed LBD37 and LBD38 within the early transcriptional cascade downstream of NLP-mediated nitrate signaling and identified them as transcription factors associated with nitrate-dependent repression of gene expression (Brooks et al. 2019; Alvarez et al. 2020). Even though LBDs have long been implicated in the negative regulation of N-responsive gene expression, how LBD37, LBD38, and LBD39 function within endogenous N response regulatory systems has remained unclear.

In this study, we show that LBD37, LBD38, and LBD39 act as major negative regulators that restrain nitrate responses and N-starvation responses under high N availability. This restraint is mediated through two distinct regulatory pathways: (i) promoter-associated repression of genes involved in N uptake and assimilation and (ii) organ-autonomous repression of *CEP* and *CEPD* expression, which gates the systemic N-demand signaling relay. Given that *LBD*s are expressed in response to local N availability, these findings indicate that LBDs couple local N status to both local N uptake and assimilation and systemic N-demand signaling, thereby coordinating N acquisition and utilization at the whole-plant level as N availability and demand changes.

## RESULTS

### *LBD37*, *LBD38*, and *LBD39* expression responds to local nitrogen availability

First we measured the expression levels of *LBD37*, *LBD38*, and *LBD39* by quantitative reverse transcription PCR (RT-qPCR) in Col-0 (wild type) seedlings incubated under high-N-availability (+N) or N-starved (–N) conditions, corresponding to agar plates containing 0.5× MS-based medium supplemented with 10.3 mM NH_4_NO_3_ and 9.4 mM KNO_3_ or lacking N, respectively (Figure 1A). The transcript levels of all *LBD*s were higher in +N-grown seedlings than in –N-grown seedlings in both shoots and roots, and the results concurred with those reported by Rubin et al. (2009).

**Figure 1.**
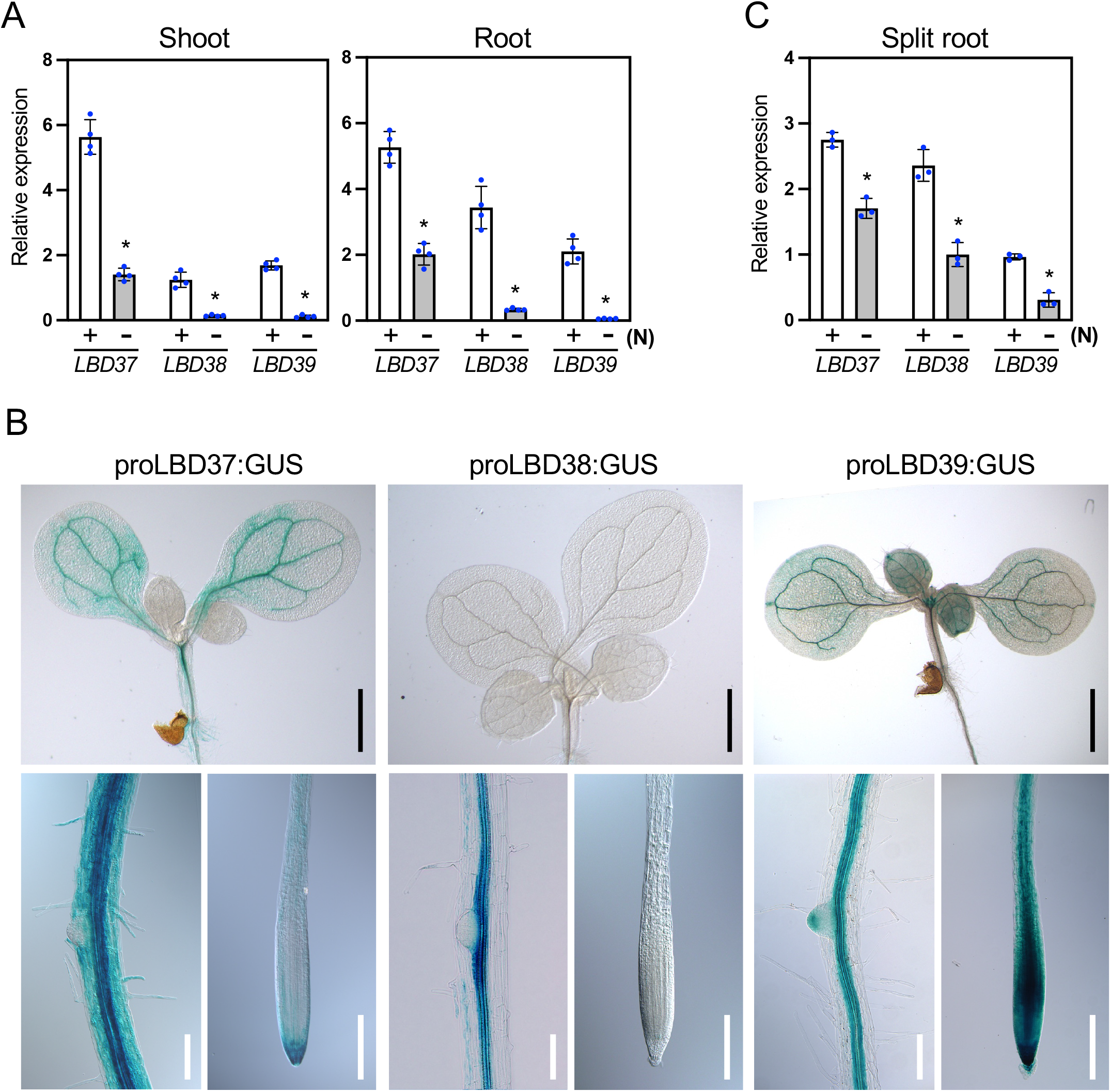
Expression patterns of LBD37, LBD38, and LBD39 under high nitrogen availability and nitrogen starvation. **(A)** Expression of *LBD37*, *LBD38*, and *LBD39* (*LBD*s) in shoots and roots of Col-0 (wild type) seedlings under +N and −N conditions. Seven-day-old Col-0 seedlings were transferred to 0.5× MS agar plates containing either 10.3 mM NH_4_NO_3_ and 9.4 mM KNO_3_ (+N) or no nitrogen (−N) and incubated for 3 days. **(B)** Spatial expression patterns of *LBD*s revealed by GUS staining in 7-day-old transgenic seedlings harboring *LBD* promoter:*GUS* reporter constructs grown on +N plates. Black scale bars, 1 mm; white scale bars, 100 µm. **(C)** Expression of *LBD*s in the +N- and −N-exposed root branches of Col-0 seedlings grown in a split-root culture system. Seedlings were grown in two-compartment Petri dishes such that one lateral root branch of each seedling was exposed to +N and the other to −N conditions for 3 days. (A) and (C) Expression levels were analyzed by RT-qPCR and normalized to *TIP41* as an internal control. Error bars represent SD (A, *n* = 4 independent pools of the indicated tissue from 10–20 seedlings; C, *n* = 3 independent pools of the indicated tissue from 2 seedlings). Asterisks indicate statistically significant differences between +N and −N (*P* < 0.05, Student’s *t* test).

Next, we analyzed the spatial expression patterns of *LBD*s in transgenic seedlings harboring a fusion between the *LBD* promoter and the *GUS* reporter gene (Figures 1B and S1). Consistent with the RT-qPCR results shown in Figure 1A, GUS staining was detected in both shoots and roots (Figure 1B) and was generally stronger under +N than under −N conditions (Figure S1), except that no staining was observed in shoots of *proLBD38:GUS* lines. In seedlings grown under +N conditions, GUS staining was particularly strong in the vascular tissues of shoots and roots (when present), although it was also observed in other tissues. At the root tip, the expression patterns differed among *LBD*s: *LBD37* expression was strong around the root cap, whereas *LBD39* expression was prominent in the developing vascular region and around the root cap. By contrast, no *LBD38* promoter activity was detected in the root tip.

We then tested whether *LBD* expression is regulated by local N availability. *LBD* expression was analyzed in Col-0 seedlings treated in a split-root culture system, in which one lateral root branch of each seedling was exposed to +N conditions and the other to −N conditions (Tabata et al. 2014; Figure 1C). The expression levels of all *LBD*s were higher in the root branch exposed to +N compared with that exposed to –N, indicating that *LBD* expression responds to local N availability. Similar results were obtained in a previous split-root transcriptome analysis (Ruffel et al. 2011), and these findings are also consistent with previous reports showing that *LBD*s are directly regulated by NLPs (Alvarez et al. 2020; Liu et al. 2022; Durand et al. 2025).

### LBD37, LBD38, and LBD39 act redundantly to control nitrogen status under high nitrogen availability

Although the effects of LBD overexpression on N responses have been reported previously (Rubin et al. 2009; Albinsky et al. 2010), the endogenous roles of LBDs have not been well examined using mutants. To investigate this, we generated *lbd37 lbd38 lbd39* triple mutants either using the CRISPR-Cas9 system or by combining T-DNA insertion alleles. A transfer RNA-based multiplex CRISPR-Cas9 vector carrying six guide RNAs, two for each of the three *LBD*s, was used to simultaneously edit the three genes (Figures S2 and S3). Two triple mutant lines, *lbd37-4 lbd38-3 lbd39-3* (*lbdT-c1*) and *lbd37-5 lbd38-4 lbd39-4* (*lbdT-c2*), were obtained from independent T1 plants (Figure S2). The *lbd37-4*, *lbd37-5, lbd39-3,* and *lbd39-4* alleles each contained a one-base insertion that causes a frameshift and were predicted to encode proteins consisting only of a truncated DNA-binding domain (Figure S3). The *lbd38-3* allele carried a 466-bp deletion that removed the C-terminal half of the DNA-binding domain, whereas the *lbd38-4* allele carried a 433-bp deletion and was predicted to encode a protein retaining only a truncated DNA-binding domain (Figure S3). Because these mutant proteins were unlikely to function as transcription factors, we considered them as null alleles. Two triple T-DNA insertion mutant lines, *lbd37-3 lbd38-2 lbd39-2* (*lbdT-t1*) and *lbd37-3 lbd38-1 lbd39-2* (*lbdT-t2*) were produced by crossing the corresponding single mutants (Figures S2A and S2F). The T-DNA insertions in *lbd37-3*, *lbd38-1*, and *lbd38-2* were located within the region encoding the DNA-binding domain and were therefore likely to disrupt it, whereas the insertion in *lbd39-2* was located downstream of this region.

The *lbdT-c1* and *lbdT-c2* seedlings grown under +N conditions showed reduced shoot and root growth compared with Col-0, as indicated by lower fresh weight, shorter primary roots, and reduced lateral root density (Figures 2A, 2B, S4A, and S4B). When grown on nutrient-rich soil, reduced shoot growth was also evident at the vegetative stage (Figures S5A and S5B). Although *lbdT-c1* and *lbdT-c2* took longer to flower than Col-0, the number of rosette leaves at flowering was on par (Figures S5C and S5D), and by flowering they had reached a rosette diameter comparable to that of Col-0 (Figures S5E and S5F), indicating that their reduced shoot growth at the vegetative stage reflected slower growth progression rather than a reduced final rosette size. The *lbdT-t1* and *lbdT-t2* mutants exhibited similar but weaker growth phenotypes than *lbdT-c1* and *lbdT-c2* (Figures 2, S4, and S5), possibly because *lbd39-2* is not a null allele. Consistently, the double mutants, *lbd37-4 lbd38-3*, *lbd37-4 lbd39-3*, and *lbd38-3 lbd39-3*, generated by backcrossing *lbdT-c1* with Col-0, showed no apparent growth defect at the seedling stage (Figure S6A).

**Figure 2.**
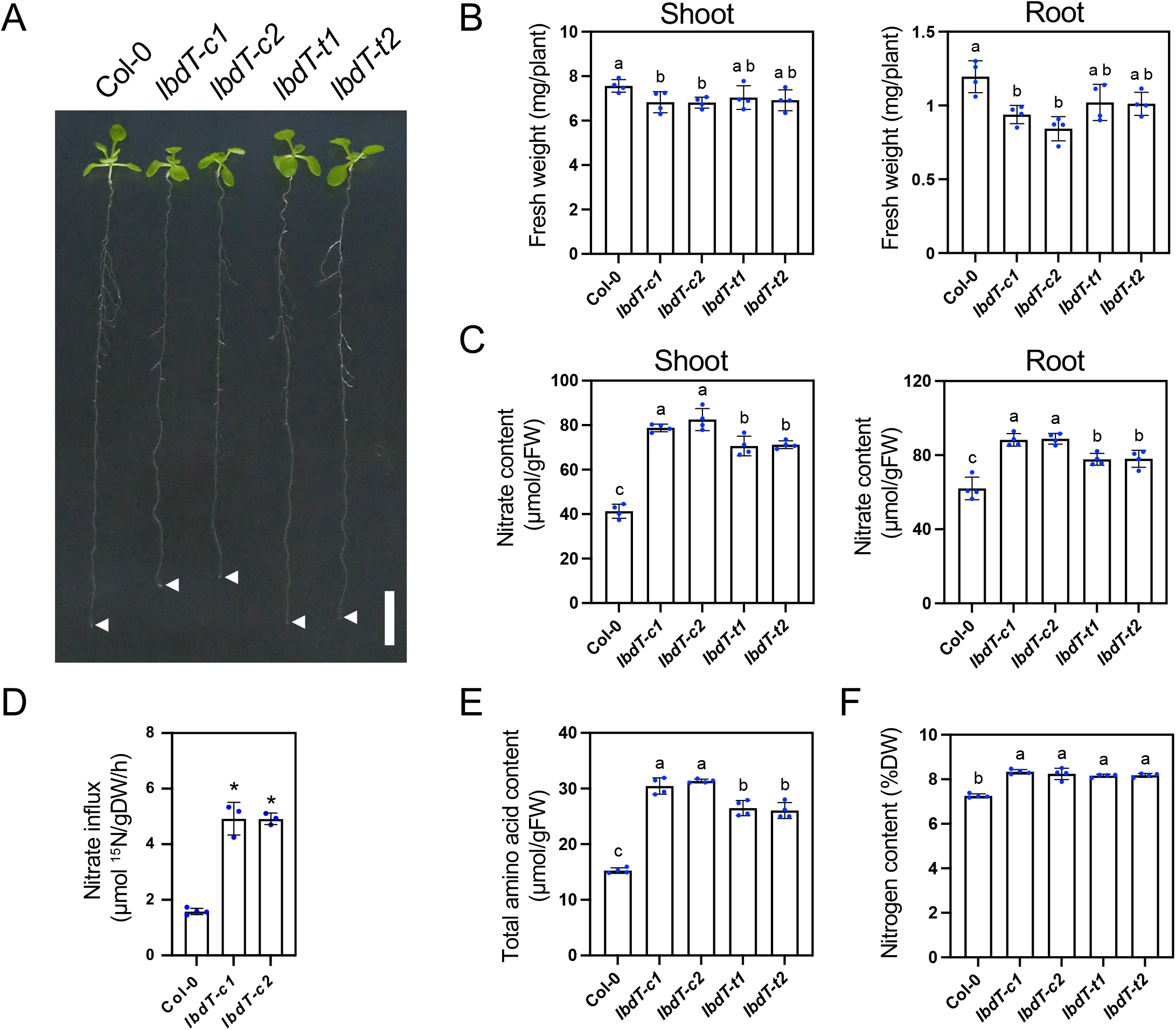
Phenotypes of *lbd37 lbd38 lbd39* triple mutant seedlings grown under high nitrogen availability. **(A)** and **(B)** Shoot and root growth of Col-0 (wild type) and *lbd37 lbd38 lbd39* triple mutant lines (*lbdT*) grown for 11 days on +N agar plates. A representative image (A) and shoot and root fresh weights (B) of the mutants are shown. *lbdT-c1* and *lbdT-c2* were generated using the CRISPR-Cas9 system; *lbdT-t1* and *lbdT-t2* were generated by crossing T-DNA insertion lines. White arrowhead indicates the position of primary root tip. Scale bar, 1 cm. **(C)** to **(F)** Nitrate content (C), high-affinity nitrate influx measured at 0.2 mM nitrate (D), total free amino acid content (E), and total nitrogen content (F) in Col-0 and *lbdT* lines grown for 11 days on +N agar plates. (B) to (F) Error bars represent SD (B, *n* = 4 independent pools of the indicated organ from 10 seedlings; C to F, *n* = 3– 4 independent pools of 10–20 seedlings). Asterisks indicate significant differences compared with Col-0, as evaluated by Student’s *t* test (*P* < 0.05). Different lowercase letters indicate significant differences, as determined by Tukey’s multiple comparisons test at *P* < 0.05.

Under +N conditions, *lbdT-c1* and *lbdT-c2* accumulated more nitrate than Col-0 in both shoots and roots, with shoot nitrate levels nearly doubling (Figure 2C). High-affinity nitrate influx also increased more than twofold (Figure 2D). Similarly, total free amino acid content more than doubled in the mutants (Figure 2E), largely because of increases in glutamine, glutamate, arginine, and asparagine content (Figure S4C). Total N content also increased in the mutant (Figure 2F). Similar increases in nitrate and amino acid accumulation were observed when 15 mM nitrate was supplied as the sole N source (Figure S7). The *lbdT-t1* and *lbdT-t2* mutants showed similar but less pronounced N accumulation phenotypes (Figures 2 and S4). The double mutants also exhibited increased nitrate accumulation relative to Col-0, although the increase was much smaller than that observed in the triple mutants (Figure S6B). Together, these results indicate that the three *LBD* genes, which act redundantly, play important roles in the control of growth and N status under high N availability.

### LBDs are transcriptional repressors that suppress nitrogen-starvation- and nitrate-inducible genes under high nitrogen availability

Although overexpression of *LBD*s has been reported to reduce the expression of N-responsive genes (Rubin et al. 2009), whether LBD proteins possess intrinsic transcriptional repressor activity has not been directly examined. To test this, we conducted Arabidopsis mesophyll protoplast co-transfection assays (co-transfection assays) using a reporter construct harboring a *GAL4 promoter:Firefly luciferase* fusion and effector constructs expressing LBD proteins fused to the GAL4 DNA-binding domain (GBD) (Figure S8A). As expected, GBD-NIGT1.1, a known transcriptional repressor (Kiba et al. 2018), reduced *GAL4* promoter activity relative to the control (GBD-GFP). Expression of GBD-LBD fusion proteins reduced reporter activity in a similar manner, indicating that LBD proteins act as transcriptional repressors (Figure S8B).

To identify genes negatively regulated by LBDs, we performed transcriptome analysis using total RNA from shoots and roots of Col-0 and *lbdT-c1* seedlings grown under +N conditions. Compared to Col-0, a total of 401 genes were upregulated in the shoots and 529 genes in the roots of *lbdT-c1* mutants (Tables S1 and S2). KEGG pathway enrichment analyses of these genes identified “N metabolism” as the most significantly enriched pathway in both organs (Tables S3 and S4). To determine the nature of LBD regulation, we compared the upregulated genes in *lbdT-c1* (*lbdT* UP) with genes induced by N starvation (N-stv UP; Tables S5 and S6) and previously reported primary nitrate-inducible genes (Nitrate UP; Table S7; Liu et al. 2022). This analysis revealed significant overlap of *lbdT* UP genes with both N-stv UP and Nitrate UP genes in shoots and roots (Figure 3A; Tables S8 and S9). The overlapping category included genes encoding N-starvation signaling and transcription components, such as *CEP* genes (*CEP5*, *CEP6*, and *CEP7*; Tabata et al. 2014), *CEPD* genes (*CEPD1*, *CEPD2*, and *CEPDL2*; Ohkubo et al. 2017; Ota et al. 2020), and *NF-YA10* (Ye et al. 2025), as well as canonical nitrate-inducible genes involved in nitrate uptake, assimilation, and signaling and transcription, including *NRT2.1*, *NRT2.2, NRT3.1*/*NAR2.1* (*NAR2.1*), *NIA1*, *NIR1*, *CEPH*, *BTB-TAZ DOMAIN PROTEIN1* (*BT1*), *BT2, NIGT1.3,* and *NLP3* (Wang et al. 2018). Notably, a subset of these genes, such as *NRT2.1* and *NAR2.1*, showed dual responsiveness to N starvation and nitrate, aligning with both the N-stv UP and Nitrate UP categories. RT-qPCR analysis confirmed that representative *lbdT* UP genes overlapping with the N-stv UP and/or Nitrate UP gene sets were upregulated in two *lbdT* mutant lines relative to Col-0 under +N conditions (Figures 3B and 3C). In the double mutants, the extent of upregulation of representative genes was smaller than that in *lbdT-c1*, although the magnitude of difference varied among double-mutant combinations and representative genes (Figures S6C and S6D). Together, these results indicate that LBDs redundantly suppress both N-starvation-inducible and primary nitrate-inducible genes under high N availability.

**Figure 3.**
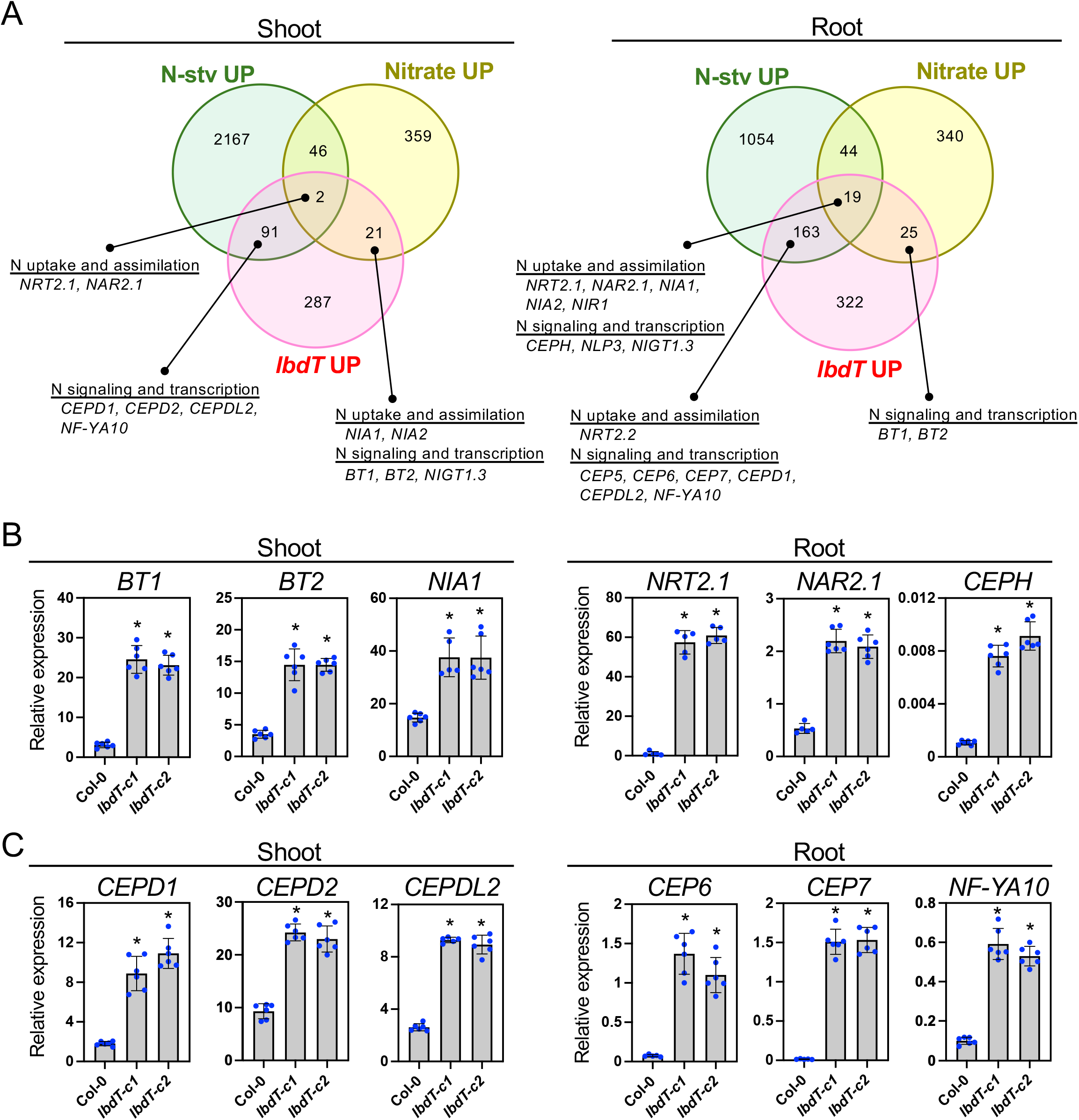
Transcriptome analysis of the *lbd37 lbd38 lbd39* triple mutant grown under high nitrogen availability. **(A)** Venn diagrams showing the overlap between N-starvation-inducible genes (Nstv UP), primary nitrate-inducible genes (Nitrate UP), and upregulated genes in *lbdT-c1* (*lbdT* UP) in shoots (left) and roots (right). Genes upregulated in Col-0 seedlings incubated for 3 days under N-starved conditions (Nstv UP; FC > 2, *P* < 0.05, FDR < 0.1), primary nitrate-inducible genes (Nitrate UP; Liu et al. 2022), and genes upregulated in the *lbdT-c1* mutant (*lbdT* UP; FC > 2, *P* < 0.05, FDR < 0.1) were compared (Tables S1 to S7). Representative N-starvation-inducible and/or nitrate-inducible genes upregulated in *lbdT-c1* are presented. **(B)** and **(C)** Expression levels of representative *lbdT*-upregulated genes. Genes shown in (B) are nitrate-inducible genes, either with or without N-starvation inducibility. Genes shown in (C) are induced only by N starvation. Col-0, *lbdT-c1*, and *lbdT-c2* seedlings were grown for 11 days on +N agar plates. Error bars represent SD (*n* = 5–6 independent pools of the indicated tissue from 10–20 seedlings). Asterisks indicate significant differences compared with Col-0, as evaluated by Student’s *t* test (*P* < 0.05). Expression levels were analyzed by RT-qPCR and normalized to *TIP41* as an internal control.

### LBDs and NIGT1s suppress distinct and partially overlapping subsets of nitrogen-responsive genes

NIGT1s have been shown to act as repressors of both N-starvation-inducible and primary nitrate-inducible genes (Kiba et al. 2018; Maeda et al. 2018; Safi et al. 2021). Our transcriptome analysis of the *lbdT* mutant showed that LBDs also contribute to the repression of N-responsive genes in the same category (Figure 3), raising the possibility that these LBDs and NIGT1s share roles in the regulation of N-responsive gene expression. Consistent with this possibility, a significant overlap was observed between genes upregulated in *lbdT-c1* and genes downregulated in NIGT1.2-overexpressing plants (Kiba et al. 2018; Figure S9, Tables S10 and S11). In addition, expression levels of *LBD37*, *LBD38*, and *LBD39* were not significantly altered in the *nigt1.1 nigt1.2 nigt1.3 nigt1.4* quadruple mutant (*nigtQ-4*; Figure S10A), whereas we found an increase in expression levels of all *NIGT1* genes in shoots, and of *NIGT1.3* and *NIGT1.4* in roots in *lbdT* mutants (Figure S10B). Together, these observations prompted us to generate the *nigt1.1 nigt1.2 nigt1.3 nigt1.4 lbd37 lbd38 lbd39* septuple mutant (*nigtQlbdT*), with the aim of assessing the individual and overlapping contributions of LBDs and NIGT1s to N-responsive gene repression.

The septuple mutants, *nigtQlbdT-1* and *nigtQlbdT-2,* were generated by crossing *nigtQ-4* with *lbdT-c1* and *lbdT-c2*, respectively. Transcriptome analysis was performed using total RNA extracted from shoots and roots of *nigtQ-4*, *lbdT-c1*, and *nigtQlbdT-1* seedlings grown under +N conditions. We compared the transcriptomes of *nigtQlbdT-1* with those of *nigtQ-4* and *lbdT-c1* to assess the effects of loss of LBD function in the *nigtQ* background and loss of NIGT1 function in the *lbdT* background, respectively. This design was intended to reveal the contributions of LBDs and NIGT1s, even in cases where these transcription factors regulate overlapping sets of genes. In this framework, we focused on genes upregulated in *nigtQlbdT-1* in each comparison (Figure 4; Tables S12 to S15). Comparison of *nigtQlbdT-1* with *nigtQ-4* identified 1,060 genes to be upregulated in shoots and 533 in roots, while comparison with *lbdT-c1* revealed 836 genes to be upregulated in shoots and 240 in roots. The genes identified in these comparisons were designated LBD-repressed and NIGT1-repressed genes, respectively (Tables S12 to S15). Venn diagram analyses of these gene sets with the N-stv UP (Tables S5 and S6) and Nitrate UP (Table S7) gene sets revealed that substantial fractions of both LBD-repressed and NIGT1-repressed genes overlapped with N-starvation-inducible and/or primary nitrate-inducible genes (Figure 4A; Table S16). Specifically, 54% of shoot and 40% of root LBD-repressed genes, and 64% of shoot and 33% of root NIGT1-repressed genes, overlapped with these N-responsive gene sets, with more N-starvation-inducible genes than primary nitrate-inducible genes. These analyses further revealed non-overlapping and overlapping subsets between the LBD-repressed and NIGT1-repressed gene sets. The non-overlapping subset likely represents genes for which either LBDs or NIGT1s preferentially contribute to repression, while the overlapping subsets suggest genes for which both the transcription factor groups contribute substantially.

**Figure 4.**
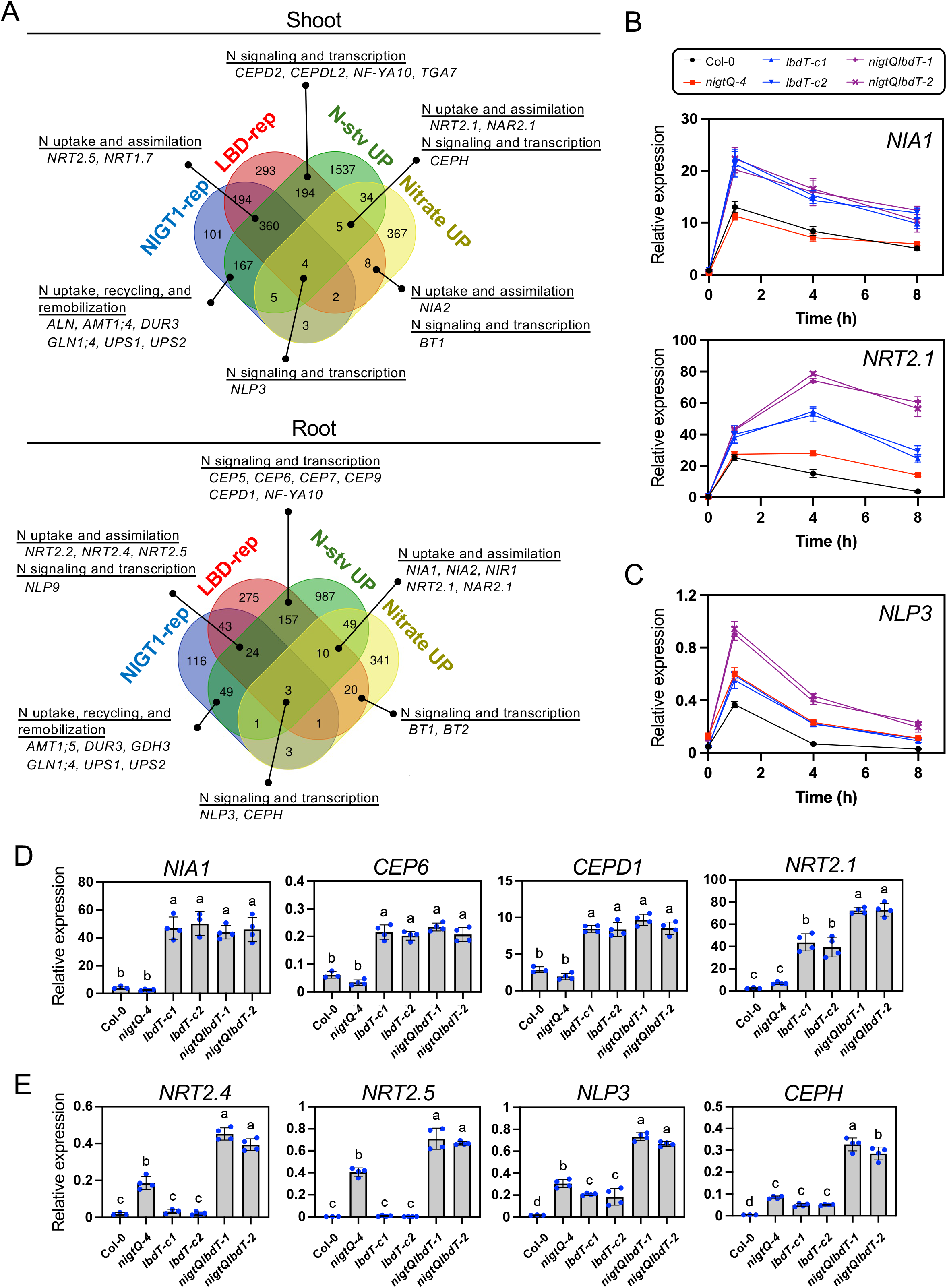
Transcriptome analysis of the *nigt1.1 nigt1.2 nigt1.3 nigt1.4 lbd37 lbd38 lbd39* septuple mutant grown under high nitrogen availability. **(A)** Venn diagrams showing overlaps among NIGT1-repressed genes (NIGT1-rep), LBD-repressed genes (LBD-rep), N-starvation-inducible genes (Nstv UP), and primary nitrate-inducible genes (Nitrate UP) in shoots and roots. NIGT1-rep genes were defined as genes upregulated in the *nigt1.1 nigt1.2 nigt1.3 nigt1.4 lbd37 lbd38 lbd39* septuple mutant (*nigtQlbdT-1*) compared with *lbdT-c1*, and LBD-rep genes were defined as genes upregulated in *nigtQlbdT-1* compared with *nigtQ-4* (FC > 2, *P* < 0.05, FDR < 0.1). Nstv and Nitrate UP genes are the same as those in Figure 3A. Representative genes in the overlapping regions are indicated. See also Tables S12 to S16. **(B)** and **(C)** Time-course expression patterns of representative nitrate-inducible genes classified as non-overlapping LBD-repressed genes (B) or overlapping LBD/NIGT1-repressed genes (C) shown in (A). Col-0, *nigtQ-4*, *lbdT-c1*, *lbdT-c2*, *nigtQlbdT-1*, and *nigtQlbdT-2* were grown for 8 days in 0.5× MS liquid medium containing 0.5 mM ammonium succinate and then supplemented with 3 mM KNO_3_. **(D)** and **(E)** Expression levels of representative N-starvation-inducible genes classified as non-overlapping LBD-repressed genes (D) or overlapping LBD/NIGT1-repressed genes (E) shown in (A), in roots of Col-0, *nigtQ-4*, *lbdT-c1*, *lbdT-c2*, *nigtQlbdT-1*, and *nigtQlbdT-2*. Seedlings were grown for 11 days on +N agar plates. Error bars represent SD (*n* = 3-4 independent pools of the indicated tissue from 10–20 seedlings). Different lowercase letters indicate significant differences, as determined by Tukey’s multiple comparisons test at *P* < 0.05. Expression levels were analyzed by RT-qPCR and normalized to *TIP41* as an internal control.

To verify this RNA-seq-based classification, we performed RT-qPCR analysis of representative N-responsive genes from the non-overlapping and overlapping subsets of LBD-repressed genes in Col-0, *nigtQ-4*, *lbdT-c1*, *lbdT-c2*, *nigtQlbdT-1*, and *nigtQlbdT-2* (Figures 4B to 4E, S11, and S12). We performed time-course experiments and analyzed the expression of nitrate-inducible genes classified as non-overlapping LBD-repressed genes (*NIA1*, *NRT2.1*, *BT1*, *NIA2*, and *NAR2.1*) and an overlapping LBD/NIGT1-repressed gene (*NLP3*) (Figures 4B, 4C, S11A, and S12). After nitrate addition, all six genes were induced in all genotypes, with peak expression between 1 and 4 hours. In the non-overlapping genes, expression levels were comparable between Col-0 and *nigtQ-4* or only slightly elevated in *nigtQ-4*, whereas they were significantly higher in *lbdT*s and *nigtQlbdT*s than in *nigtQ-4*, except for *NAR2.1* at 8 hours (Figures 4B, S11A, and S12A to S12E), confirming a preferential contribution of LBDs to their repression. By contrast, *NLP3* expression was elevated to a similar extent in *nigtQ-4* and *lbdT*s and was further upregulated in *nigtQlbdT*s (Figures 4C and S12F), consistent with substantial contributions from both LBDs and NIGT1s.

We next analyzed N-starvation-inducible genes from the non-overlapping and overlapping LBD-repressed subsets in +N-grown seedlings (Figures 4D, 4E, S11B, and S11C). The non-overlapping LBD-repressed genes in roots (*NIA1*, *CEP6*, *CEPD1*, and *NRT2.1*) and shoots (*NF-YA10* and *CEPH)* showed no upregulation in *nigtQ-4* but showed higher expression in *lbdT*s and *nigtQlbdT*s than in Col-0 (Figures 4D and S11B), again confirming a preferential LBD contribution. In contrast, the overlapping genes, (*NRT2.4*, *NRT2.5*, *NLP3*, and *CEPH* in roots and *NLP3* and *NRT2.5* in shoots) showed elevated expression in *nigtQ-4* compared with Col-0. Although their expression levels were comparable between Col-0 and *lbdT*s or only slightly higher in *lbdT*s, they were significantly higher in *nigtQlbdT*s than in *nigtQ-4*, supporting substantial contributions from both LBDs and NIGT1s to their repression (Figures 4E and S11C).

Having confirmed that the expression patterns of representative genes generally followed their RNA-seq-based classifications, we designated the non-overlapping LBD-repressed subset as LBD-preferentially repressed; the non-overlapping NIGT1-repressed subset as NIGT1-preferentially repressed; and the overlapping LBD/NIGT1-repressed subset as LBD/NIGT1 co-repressed gene sets. We then examined the biological processes associated with these gene sets by GO term enrichment analysis (Table S17). In roots, the NIGT1-preferentially repressed gene set was enriched for nutrient-related terms, including “ammonium transmembrane transport”, “response to starvation”, and “response to nutrient levels”. By contrast, no nutrient-related terms were enriched in the NIGT1-preferentially repressed gene set in shoots. Nevertheless, examination of the shoot gene lists identified genes involved in N uptake, recycling, and remobilization (Figure 4A; Table S16), such as *ALN, AMT1;4*, *DUR3*, *GLN1;4*, *UPS1*, and *UPS2* (Kojima et al. 2007; Yuan et al. 2007, 2009; Masclaux-Daubresse et al. 2010; Werner and Witte 2011). These features are consistent with the previously described role of NIGT1s in N-starvation responses (Kiba et al. 2018).

The LBD/NIGT1 co-repressed gene set in roots was enriched for nitrate-related terms, including “nitrate transmembrane transport”, “response to nitrate”, “nitrate metabolic process”, and “nitrate assimilation” (Table S17). This gene set contained nitrate transporter genes involved in N-starvation responses, such as *NRT2.2*, *NRT2.4*, and *NRT2.5* (Kiba et al. 2012; Lezhneva et al. 2014). It also included *CEPH*, *NLP3*, and *NLP9*, genes related to signaling and transcriptional regulation in both N-starvation and nitrate responses (Ohkubo et al. 2021; Lee et al. 2022). Although no nutrient-related terms were enriched in the corresponding shoot gene set, it included *NRT2.5*, *NLP3*, and *NRT1.7*, which are involved in nitrate remobilization upon N starvation (Fan et al. 2009). These observations indicate that the LBD/NIGT1 co-repressed gene set, particularly in roots, includes genes related to N-starvation and nitrate responses.

In the LBD-preferentially repressed gene set, nitrate-related terms were enriched in both shoots and roots, including “nitrate metabolic process”, “nitrate assimilation”, and “nitrate import” (Table S17). Notably, the root gene set was also enriched for “cellular response to N starvation”. These enriched processes were represented by canonical nitrate-inducible genes involved in nitrate uptake, assimilation, and signaling, including *NRT2.1*, *NAR2.1*, *NIA1*, *NIA2*, *NIR1*, *BT1*, and *BT2* (Wang et al. 2018). This gene set also included genes involved in N-starvation signaling, including *CEP* genes (Tabata et al. 2014) and *CEPD* genes (Ohkubo et al. 2017; Ota et al. 2020), as well as transcriptional regulators involved in N-starvation responses, such as *NF-YA10* and *TGA7* (Ye et al. 2025; Figure 4A; Table S16).

Thus, compared with the NIGT1-preferentially repressed gene set, LBD-repressed genes (encompassing both the LBD-preferentially repressed and LBD/NIGT1 co-repressed gene sets) showed greater representation of genes involved in N-starvation and nitrate responses.

### The *lbdT* mutants exhibit more severe nitrogen-related physiological phenotypes than *nigtQ*

We next investigated whether the differences revealed by the transcriptome analysis (Figure 4) were reflected in physiological phenotypes by comparing *nigtQ-4*, *lbdT*, and *nigtQlbdT* mutants (Figures 5, S13, and S14). Under +N agar-plate and nutrient-rich soil conditions, growth of *nigtQ-4* was comparable to that of Col-0, as reported previously (Figures S13A to S13D and S14A to S14D; Kiba et al. 2018). By contrast, *lbdT-c1* showed reduced growth on agar plates and at the vegetative stage on soil, as shown in Figures 2, S4, and S5. Growth of *nigtQlbdT* mutants was further reduced compared with *lbdT-c1*, as indicated by lower fresh weight and shorter primary roots on agar plates and reduced shoot growth on soil at the vegetative stage, whereas lateral root density was similar between *lbdT-c1* and *nigtQlbdT*s. *nigtQlbdT*s took longer to flower than any other genotype examined, but they reached a rosette diameter similar to that of Col-0 at flowering (Figures S14C to S14F), indicating that the reduced shoot size of *nigtQlbdT* mutants at the vegetative stage mainly reflected slower growth.

**Figure 5.**
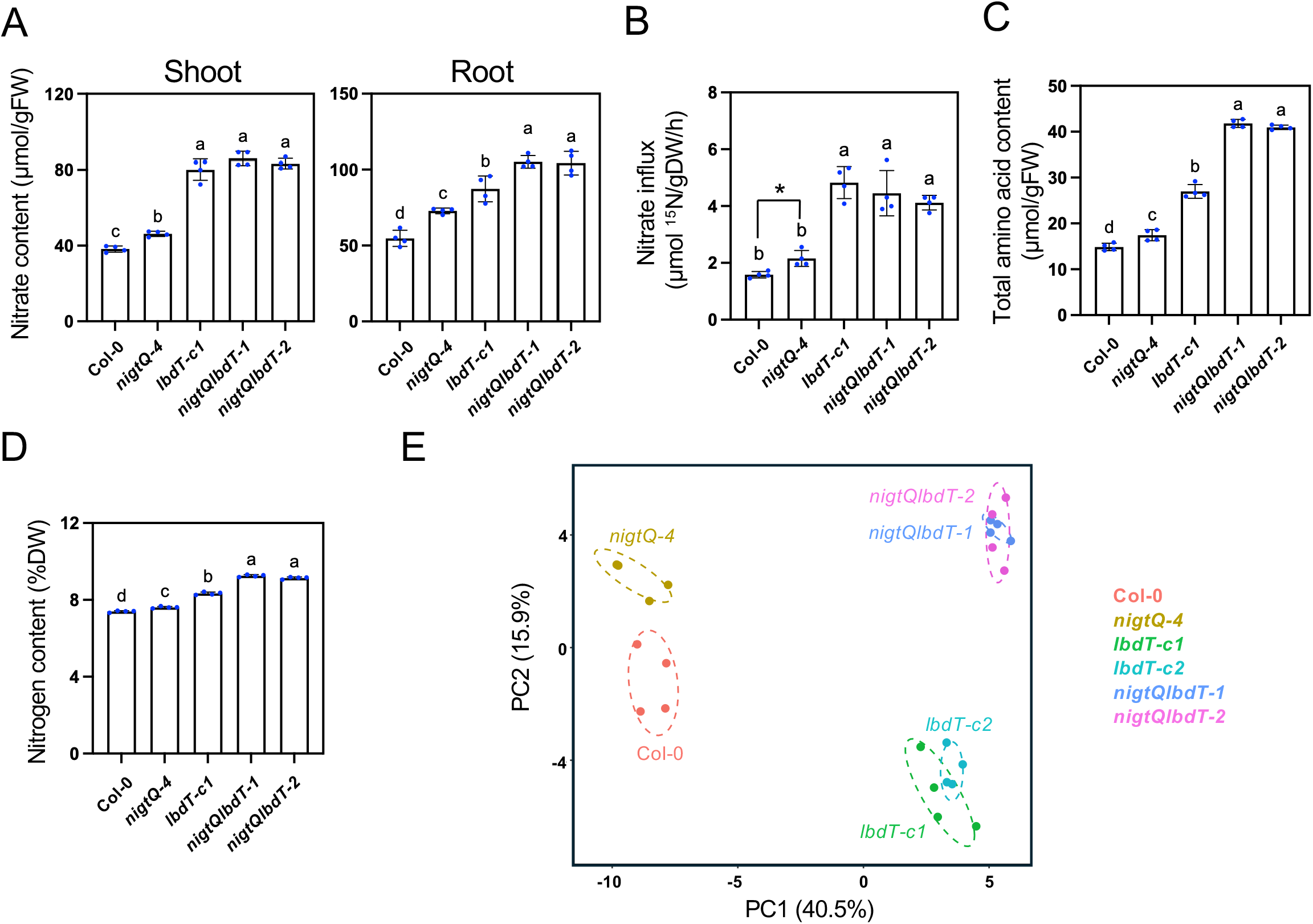
Phenotypes of *nigt1.1 nigt1.2 nigt1.3 nigt1.4 lbd37 lbd38 lbd39* septuple mutants grown under high nitrogen availability. **(A)** to **(D)** Nitrate content (A), high-affinity nitrate influx measured at 0.2 mM nitrate (B), total free amino acid content (C), and total nitrogen content (D) in Col-0, *nigtQ-4*, *lbdT-c1*, and *nigt1.1 nigt1.2 nigt1.3 nigt1.4 lbd37 lbd38 lbd39* septuple mutants (*nigtQlbdT-1* and *nigtQlbdT-2*). **(E)** Principal component analysis (PCA) score plot of metabolome data obtained from Col-0, *nigtQ-4*, *lbdT-c1*, *lbdT-c2*, *nigtQlbdT-1*, and *nigtQlbdT-2*. Symbols represent four biological replicates from each genotype. Seedlings were grown for 11 days on +N agar plates. Error bars represent SD (*n* = 3–4 independent pools of 10–20 seedlings). Different lowercase letters indicate significant differences, as determined by Tukey’s multiple comparisons test at *P* < 0.05. Asterisks indicate significant differences compared with Col-0, as evaluated by Student’s *t* test (*P* < 0.05).

We then measured N-related physiological parameters in seedlings grown under +N conditions (Figures 5 and S13E). Consistent with previous findings, *nigtQ-4* showed elevated nitrate contents in shoots and roots, high-affinity nitrate influx, and total amino acid and N contents compared with Col-0 (Kiba et al. 2018; Safi et al. 2021). These parameters were more strongly affected in *lbdT-c1* than in *nigtQ-4*. Although shoot nitrate content and high-affinity nitrate influx were comparable between *lbdT-c1* and *nigtQlbdT*s, root nitrate content and total amino acid and N contents were higher in *nigtQlbdT*s than in *lbdT-c1* (Figures 5A to 5D). Similar increases in nitrate and amino acid levels in *lbdT-c1* and *nigtQlbdT*s were observed when 15 mM nitrate was supplied as the sole N source (Figure S15).

To assess the broader metabolic consequences of these mutations, we performed widely targeted metabolome analysis in +N-grown Col-0, *nigtQ-4*, *lbdT-c1*, *lbdT-c2*, *nigtQlbdT-1*, and *nigtQlbdT-2* (Figure 5E; Table S18). After quality control and metabolite curation, the relative peak areas of 100 metabolites were subjected to principal component analysis (PCA). PC1 and PC2 explained 40.5% and 15.9% of the total variance, respectively, and the PCA score plot showed genotype-dependent separation along these two axes. Along PC1, *lbdTs* and *nigtQlbdTs* occupied similar positions and were clearly separated from Col-0 and *nigtQ-4,* which were positioned close to each other, indicating that the major metabolic variation captured by PC1 was primarily associated with loss of LBD function. Along PC2, *nigtQ-4* was separated from Col-0 and *nigtQlbdT*s were separated from *lbdT*s in the same direction, suggesting that the loss of NIGT1 function was associated with a smaller but distinct component of metabolic variation.

Collectively, these results show that both LBDs and NIGT1s act to suppress N-related responses, including nitrate uptake and assimilation as well as amino acid metabolism, under high N availability. Furthermore, the more pronounced N-related phenotypes of *lbdT*s and *nigtQlbdT*s compared with *nigtQ-4* indicate that LBDs play a more prominent role than NIGT1s in this regulation.

### LBDs suppress systemic nitrogen-demand signaling by repressing N-starvation-inducible *CEP* and *CEPD* genes mainly in an organ-autonomous manner

To explore the mechanism by which LBDs suppress N responses under high N availability, we focused on genes involved in systemic N-demand signaling. Transcriptome analysis showed that multiple *CEP* and *CEPD* genes were upregulated in *lbdT*s and were included among LBD-repressed genes (Figures 3 and 4). Because several *CEP*s are not represented in the TAIR10 gene annotation and therefore could not be properly evaluated in our RNA-seq analysis, we examined the expression of *CEP1* to *CEP9*, together with *CEPD1*, *CEPD2*, *CEPDL1*, and *CEPDL2*, by RT-qPCR in Col-0 seedlings incubated under −N conditions and in Col-0, *nigtQ-4*, *lbdT-c1*, *lbdT-c2*, *nigtQlbdT-1*, and *nigtQlbdT-2* seedlings grown under +N conditions (Figure 6A). None of the *CEP* genes examined were detected in shoots, and *CEP1* and *CEP2* were not detected in roots under our experimental conditions. Among the remaining *CEP* and *CEPD* genes, N-starvation induced the expression of all *CEP* genes except *CEP4* in roots and all *CEPD* genes except *CEPDL1* in shoots, consistent with previous reports (Tabata et al. 2014; Ohkubo et al. 2017; Ota et al. 2020). *CEPD1, CEPD2,* and *CEPDL2* were induced by N starvation in roots under our experimental conditions. In *lbdT*s and *nigtQlbdT*s grown under +N conditions, all N-starvation-inducible *CEP* and *CEPD* genes were upregulated. Notably, the expression levels of these genes in +N-grown *lbdT*s and *nigtQlbdT*s were comparable to or higher than those in N-starved Col-0, with only one exception, *CEP9* (Table S19). By contrast, none of the examined *CEP* and *CEPD* genes was significantly upregulated in *nigtQ-4*. Thus, our data suggests that the majority of *CEP* and *CEPD* genes require LBDs for repression under high N availability and may require relief from LBD-mediated repression for their induction upon N starvation.

**Figure 6.**
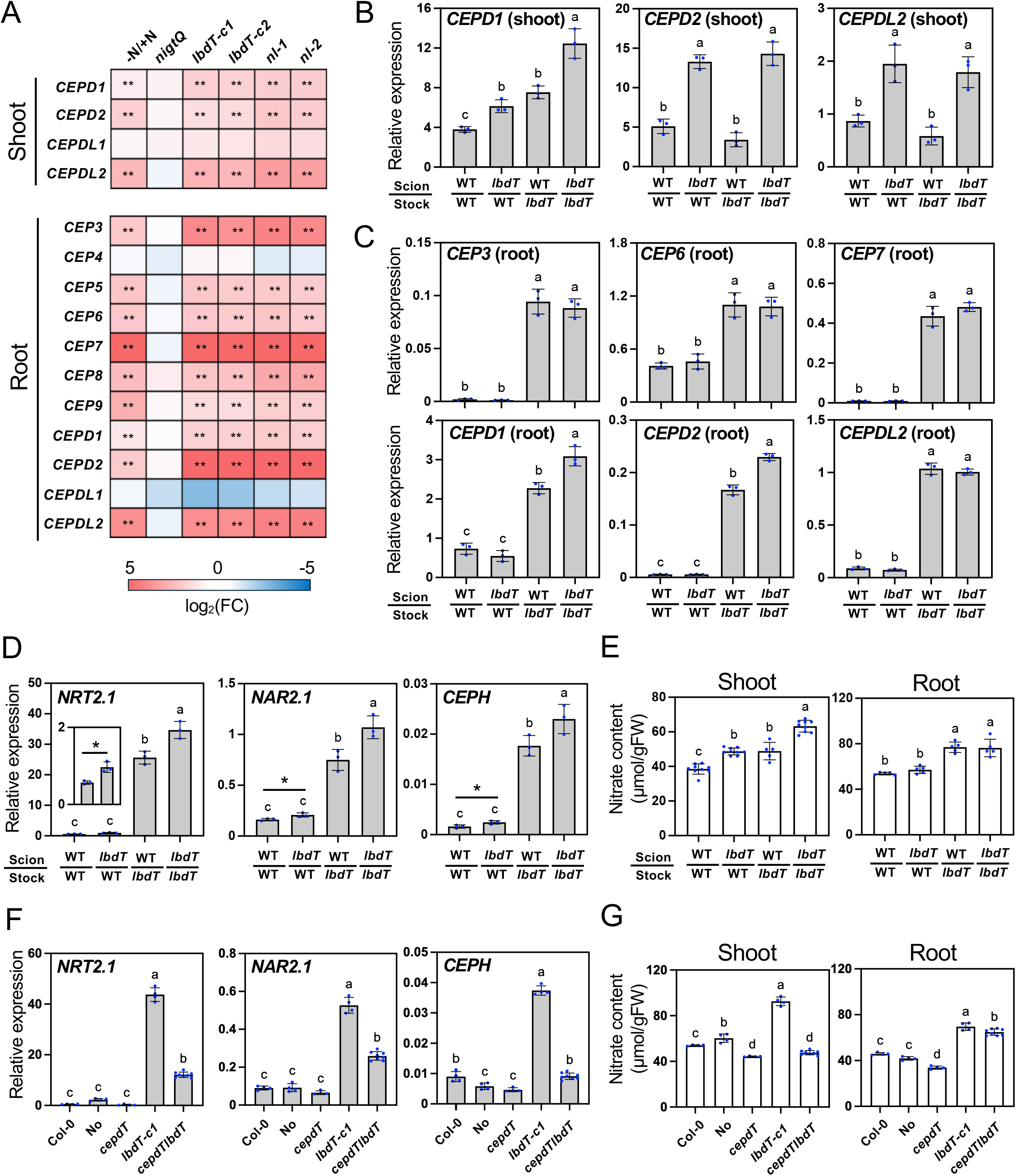
Involvement of C-TERMINALLY ENCODED PEPTIDE (CEP) and CEP DOWNSTREAM (CEPD) members in LBD-mediated repression of nitrogen responses. **(A)** Heat maps showing the effects of N starvation and mutations in *NIGT1*s, *LBD*s, or both *NIGT1*s and *LBD*s on the expression levels of *CEP* and *CEPD* genes. Changes in transcript levels are shown relative to Col-0 (wild type) grown on +N agar plates. Eight-day-old Col-0 seedlings incubated for 3 days on −N agar plates and 11-day-old Col-0, *nigtQ-4* (*nigtQ*), *lbdT-c1*, *lbdT-c2*, *nigtQlbdT-1*, and *nigtQlbdT-2* seedlings grown on +N agar plates were used. Double asterisks indicate significant differences compared with +N-grown Col-0, as evaluated by Student’s *t* test (\*\**P* < 0.01). See also Table S19. **(B)** to **(E)** Reciprocal grafting between Col-0 and *lbdT-c1*. Expression levels of *CEPD* genes in shoots (B), *CEP* and *CEPD* genes in roots (C), *NRT2.1*, *NAR2.1*, and *CEPH* in roots (D), and nitrate content in shoots and roots (E) were analyzed in grafted seedlings incubated on +N agar plates. Asterisks indicate significant differences compared with the WT/WT graft, as evaluated by Student’s *t* test (*P* < 0.05). **(F)** and **(G)** Expression levels of *NRT2.1*, *NAR2.1*, and *CEPH* in roots (F) and nitrate content in shoots and roots (G) in Col-0, Nössen (No), *cepd1 cepd2 cepdl2* (*cepdT*), *lbdT-c1*, and the *cepdT lbdT-c1* sextuple mutant (*cepdTlbdT*) grown for 11 days on +N agar plates. Because *cepdT* is in the No background, No was included as an additional wild-type control. Expression levels were analyzed by RT-qPCR and normalized to *TIP41* as an internal control. Error bars represent SD (A, *n* = 4 independent pools of the indicated organ from 10–20 seedlings; B to D, *n* = 3 independent pools of the indicated organ from 4 seedlings; E, *n* = 5–9 independent pools of the indicated organ from 4 seedlings; F and G, *n* = 4–8 independent pools of the indicated organ from 10–20 seedlings). Different lowercase letters indicate significant differences, as determined by Tukey’s multiple comparisons test at *P* < 0.05.

In systemic N-demand signaling, CEPs are produced in roots in response to N starvation and transported to shoots and upon perception by CEPR receptors induce *CEPD* expression (Tabata et al. 2014; Ohkubo et al. 2017; Ota et al. 2020). Therefore, increased expression of *CEPD* genes in *lbdT*s could reflect either organ-autonomous regulation or systemic regulation through root-derived CEP signaling. To distinguish these possibilities, we performed reciprocal grafting between Col-0 (WT) and *lbdT-c1* (*lbdT*) (Figures 6B to 6E). Compared to WT self-grafts, the expression of *CEPD2* and *CEPDL2* in shoots of grafted seedlings grown on +N agar plates was equally elevated in *lbdT*/WT (scion/rootstock) grafts and *lbdT* self-grafts but remained unchanged in WT/*lbdT* (Figure 6B). By contrast, *CEPD1* expression was increased in all graft combinations containing *lbdT*, with the highest expression observed in *lbdT* self-grafts. These results indicate that *CEPD2* and *CEPDL2* upregulation in shoots is dependent on the shoot genotype, whereas *CEPD1* upregulation in shoots is influenced by both shoot and root genotypes in an additive manner. Because elevation of *CEP3*, *CEP6*, and *CEP7* expression in roots was dependent on the root genotype (Figure 6C), the root-genotype effect on shoot *CEPD1* expression may reflect root-derived CEP signaling. In roots, expression of *CEPD1*, *CEPD2*, and *CEPDL2* was upregulated in WT/*lbdT* grafts and *lbdT* self-grafts but not in *lbdT*/WT grafts. Although *CEPD1* and *CEPD2* expression was higher in *lbdT* self-grafts than in WT/*lbdT* grafts, these patterns indicate that upregulation of *CEPD1*, *CEPD2*, and *CEPDL2* in roots was largely determined by the root genotype. Overall, *CEPD* genes in shoots and *CEP* and *CEPD* genes in roots appear to be repressed by LBDs mainly in an organ-autonomous manner, suggesting that LBDs have a dual role by acting in both shoots and roots in regulating the root–shoot–root signaling relay underlying systemic N-demand signaling. Consistent with this dual role, root expression levels of systemic N-demand signaling target genes, *NRT2.1*, *NAR2.1*, and *CEPH*, were elevated when *lbdT* was present in either the scion or rootstock and were the highest in *lbdT* self-grafts (Figure 6D). Similarly, whereas root nitrate content followed the root genotype, shoot nitrate accumulation was enhanced by *lbdT* in either the scion or rootstock and was most pronounced in *lbdT* self-grafts (Figure 6E).

To determine the genetic relationship between LBDs and systemic N-demand signaling, we generated the *cepd1 cepd2 cepdl2 lbdT-c1* sextuple mutant (*cepdTlbdT*) by crossing the *cepd1 cepd2 cepdl2* triple mutant (*cepdT*; Ota et al. 2020) with *lbdT-c1*. When the expression of *NRT2.1*, *NAR2.1*, and *CEPH* was analyzed in roots of wild-type (Col-0 and Nössen), *cepdT*, *lbdT-c1*, and *cepdTlbdT* seedlings grown under +N conditions, the elevated expression of these genes in *lbdT-c1* was significantly suppressed by the *cepdT* mutations (Figure 6F). Although the increased nitrate accumulation in roots of *lbdT-c1* was not affected by the *cepdT* mutations, accumulation in shoots was reduced in *cepdTlbdT* to a level comparable to that in *cepdT* (Figure 6G). Furthermore, elevated systemic N-demand signaling target-gene expression and shoot nitrate accumulation in *nigtQlbdT-1* were similarly attenuated in the *cepdT nigtQlbdT-1* decuple mutant, generated by crossing *cepdT* with *nigtQlbdT-1* (Figure S16). Together, these genetic analyses reveal an epistatic relationship between LBDs and CEPDs in the regulation of systemic N-demand signaling.

Considering that LBD expression responds to local N availability (Figure 1C), these results suggest that LBDs repress *CEP* and *CEPD* genes mainly in an organ-autonomous manner in shoots and roots in accordance with local N status, thereby gating the systemic N-demand signaling relay. Thus, systemic N-demand signaling constitutes a downstream pathway through which LBDs restrain N responses at the whole-plant level.

### LBDs use distinct mechanisms to repress nitrate uptake and assimilation pathway genes and *CEP*/*CEPD* genes

To investigate how LBDs repress N-responsive genes, we generated transgenic lines expressing LBD37 or LBD38 fused to two VP16 transcriptional activation domains in tandem (LBD37-VP or LBD38-VP) under the control of a β-estradiol-inducible promoter (37VP and 38VP lines). If LBDs associate with the regulatory regions of genes they repress, fusion to the VP16 activation domains would be expected to activate the expression of those genes. β-Estradiol-inducible expression of *LBD37*-*VP* and *LBD38*-*VP* was confirmed in two independent lines for each construct (Figure S17A). Because the preceding analyses placed *CEP* and *CEPD* genes downstream of LBDs (Figure 6), we first examined representative genes from these groups. Unexpectedly, expression of *CEP6*, *CEP9*, and *CEPDL2* was not significantly affected by the induction of *LBD37*-*VP* or *LBD38*-*VP* expression for 12 hours (Figure 7A). By contrast, *NIA1*, *NRT2.1*, and *NRT2.5*, representative nitrate uptake and assimilation pathway genes among the LBD-repressed genes (Figure 4), were upregulated (Figure 7B). We then performed co-transfection assays to examine whether LBDs affect the promoter activities of these genes. The activity of the Cauliflower mosaic virus 35S promoter (pro35S), which was used as a control promoter, was unaffected by LBD co-expression. In this assay, co-expression of LBD37, LBD38, or LBD39 repressed the activities of the *NIA1*, *NRT2.1*, and *NRT2.5* promoters (Figure 7C), whereas it did not significantly affect the activities of the *CEP6*, *CEP9*, and *CEPDL2* promoters (Figure S17B).

**Figure 7.**
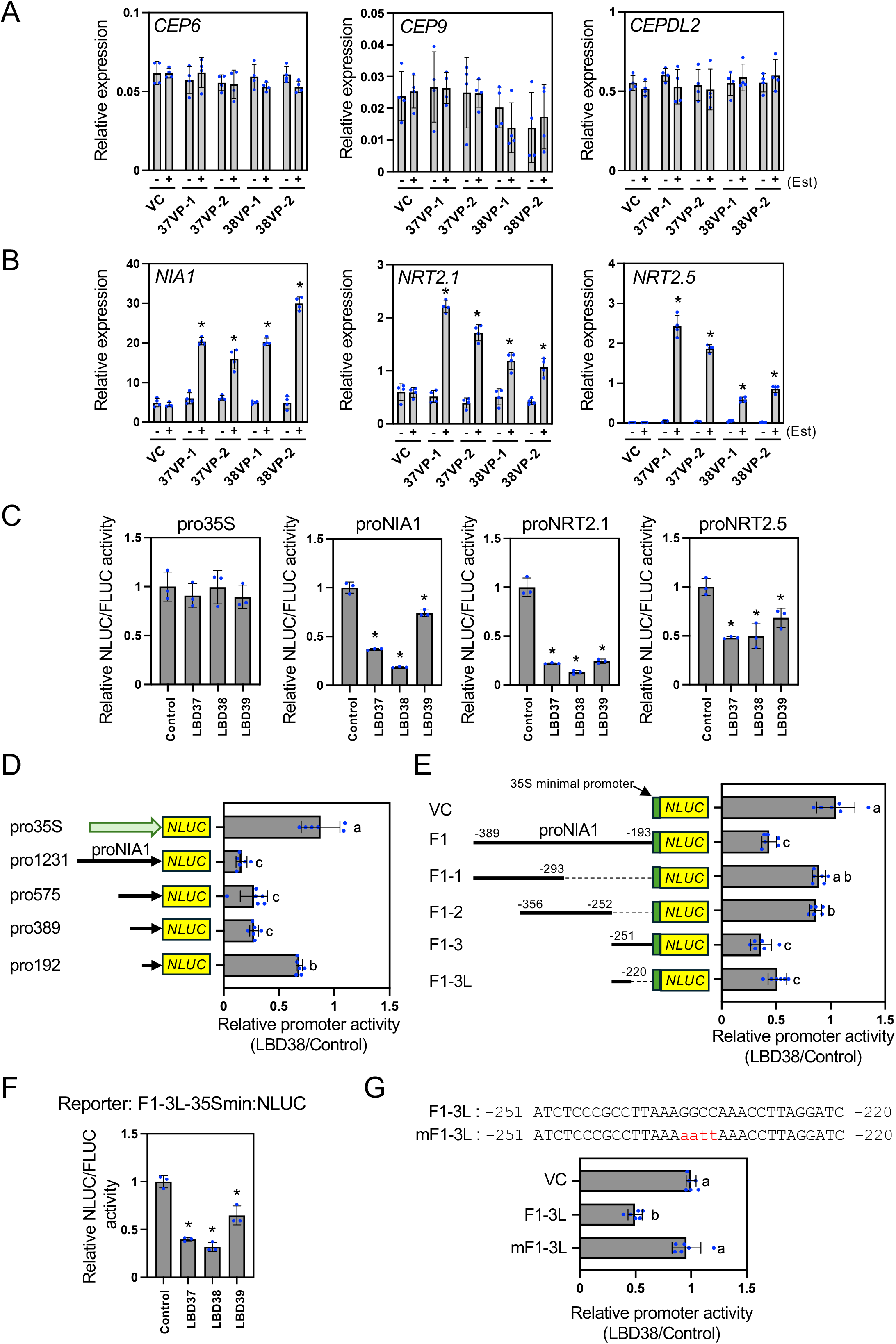
Characterization of LBD-mediated repression of *CEP* and *CEPD* genes and nitrate uptake and assimilation pathway genes. **(A)** and **(B)** Expression analysis of LBD-repressed genes in two independent transgenic lines expressing LBD37 or LBD38 fused to two VP16 transcriptional activation domains in tandem under the control of a β-estradiol (Est)-inducible promoter (37VP and 38VP). Among LBD-repressed genes identified in the transcriptome analysis, *CEP* and *CEPD* genes *CEP6*, *CEP9*, and *CEPDL2* (A) and nitrate uptake and assimilation pathway genes *NIA1, NRT2.1*, and *NRT2.5* (B) were analyzed. VC indicates the empty-vector control. Eleven-day-old seedlings grown on +N agar plates were incubated with (+) or without (−) 10 μM Est for 12 hours. Expression levels were analyzed by RT-qPCR and normalized to *TIP41* as an internal control. Asterisks indicate significant differences between the −Est and +Est treatments (*P* < 0.05, Student’s *t* test). **(C)** Effects of LBD37, LBD38, and LBD39 on the activities of the *NIA1*, *NRT2.1*, and *NRT2.5* promoters in co-transfection assays. Plasmids carrying the indicated promoters fused to *NLUC* were used as reporters. Plasmids expressing GUS (control), LBD37, LBD38, or LBD39 were used as effectors. Asterisks indicate significant differences compared with the GUS control (*P* < 0.05, Student’s *t* test). **(D)** and **(E)** Delimitation of the *NIA1* promoter region involved in LBD-mediated repression using co-transfection assays. (D) Reporter plasmids contained 1,231-, 575-, 389-, or 192-bp regions upstream of the *NIA1* start codon (pro1231, pro575, pro389, and pro192, respectively) fused to *NLUC*. (E) Reporter plasmids contained the indicated *NIA1* promoter fragments, with numbers denoting positions relative to the *NIA1* start codon, inserted upstream of the 35S minimal promoter (35Smin) and *NLUC*. The fragments were designated as F1 (−389 to −193), F1-1 (−389 to −293), F1-2 (−356 to −252), F1-3 (−251 to −193), and F1-3L (−251 to −220. The construct containing only 35Smin and *NLUC* was used as the empty reporter (VC). Plasmids expressing GUS (control) or LBD38 were used as effectors. **(F)** Effects of LBD37, LBD38, and LBD39 on the activities of the F1-3L fragment in co-transfection assays. The reporter plasmid contained the F1-3L fragment inserted upstream of 35Smin and *NLUC* (F1-3L-35Smin:NLUC). Plasmids expressing GUS (control), LBD37, LBD38, or LBD39 were used as effectors. Asterisks indicate significant differences compared with the corresponding GUS control (*P* < 0.05, Student’s *t* test). **(G)** Effect of mutations in the F1-3L fragment on LBD38-mediated repression in co-transfection assays. Reporter plasmids contained the F1-3L fragment or its mutated derivative (mF1-3L) inserted upstream of 35Smin and *NLUC*. Plasmids expressing GUS (control) or LBD38 were used as effectors. The nucleotide sequences of the native and mutated fragments are presented. (C) to (G) The plasmid containing *Firefly luciferase* (*FLUC*) driven by p35S was used as a reference. In (C) and (F), promoter activities are presented as NLUC/FLUC activity ratios. In (D), (E), and (G), relative promoter activities are presented as the ratio of NLUC/FLUC activity upon co-expression with LBD38 to that upon co-expression with GUS. Error bars represent SD (A and B, *n* = 4 independent pools of 10–20 seedlings; C and F, *n* = 3; D, E, and G, *n* = 6). Different lowercase letters indicate significant differences, as determined by Tukey’s multiple comparisons test at *P* < 0.05.

Observations on the gene expression patterns in 37VP and 38VP lines revealed that *LBD37*, *LBD38*, and *LBD39* themselves were upregulated by the induction of the fusion proteins, raising the possibility that LBD37, LBD38, and LBD39 regulate each other’s expression. Induction of LBD37-VP and LBD38-VP increased the expression of the other two *LBD* genes (Figures S18A and S18B). In co-transfection assays, co-expression of LBD37, LBD38, or LBD39 repressed the activities of the *LBD37*, *LBD38*, and *LBD39* promoters, indicating that LBDs negatively regulate *LBD* genes through their promoters (Figure S18C). Consistent with these results, expression of *LBD37*, *LBD38*, and *LBD39* was increased in *lbd38 lbd39*, *lbd37 lbd39*, and *lbd37 lbd38* double mutants, respectively (Figure S18D).

The LBD-VP inducible-expression and LBD co-transfection assays consistently distinguished *NIA1*, *NRT2.1*, *NRT2.5*, and *LBD* genes from *CEP6*, *CEP9*, and *CEPDL2*. Because neither the expression of *CEP* and *CEPD* genes in the LBD-VP inducible-expression lines nor their promoter activity in the LBD co-transfection assays was affected (Figures 7A and S17B), the observed effects of LBDs on *NIA1*, *NRT2.1*, *NRT2.5*, and the *LBD* genes themselves are unlikely to be mediated through CEPs and CEPDs. Instead, these results indicate that LBDs repress the nitrate uptake and assimilation pathway genes *NIA1*, *NRT2.1*, and *NRT2.5*, as well as the *LBD* genes themselves, through a promoter-associated mechanism, whereas repression of *CEP* and *CEPD* genes occurs through a distinct mechanism.

### LBD37, LBD38, and LBD39 repress *NIA1* promoter activity through a 32-bp region

To investigate the promoter sequences underlying LBD-mediated repression, we first examined the involvement of a previously reported LATERAL ORGAN BOUNDARIES DOMAIN protein-binding motif, referred to as the LOB motif. This motif has a 6-bp consensus sequence, GCGGCG (Husbands et al. 2007). We therefore searched for this motif within the promoter regions used in the co-transfection assays and repressed by LBDs. The *LBD37* and *NRT2.1* promoters each contained one copy of this motif, whereas the *NIA1* and *NRT2.5* promoters contained multiple LOB motif-like sequences with a single nucleotide mismatch instead (Figure S19). To determine whether these motifs or motif-like sequences were involved in LBD-mediated repression, we introduced mutations into each of them and analyzed the resulting promoters in co-transfection assays. Co-expression of LBD38 repressed the activities of all mutated promoters to an extent comparable to that observed for the corresponding native promoters, indicating that these motifs and motif-like sequences were not required for LBD-mediated repression.

We therefore sought to identify the *NIA1* promoter region involved in LBD38-mediated repression by analyzing a 5′ deletion series of the promoter (containing 1,231-, 575-, and 389-bp regions upstream of the *NIA1* start codon) in co-transfection assays. Compared with pro35S (control), the activities of pro1231, pro575, and pro389 were repressed by LBD38 co-expression (Figure 7D). Repression was significantly attenuated for pro192, indicating that the region between −389 and −193 contains a sequence involved in LBD38-mediated repression. To further delimit this region, we generated reporters harboring the −389 to −193 fragment or its sub-fragments inserted upstream of the 35S minimal promoter (Figure 7E). These fragments were designated as F1 (−389 to −193), F1-1 (−389 to −293), F1-2 (−356 to −252), F1-3 (−251 to −193), and F1-3L (−251 to −220). Co-expression of LBD38 repressed the activities of F1, F1-3, and F1-3L, but had little or no effect on F1-1 or F1-2, narrowing the region to the F1-3L sub-fragment. Similarly, co-expression of LBD37 or LBD39 also repressed the activity of F1-3L (Figure 7F). Furthermore, substitution of the central GGCC sequence in F1-3L with AATT (mF1-3L) abolished repression by LBD38 co-expression (Figure 7G). Together, these analyses identified a 32-bp region between −251 and −220 that is sufficient for repression by LBD37, LBD38, and LBD39.

## DISCUSSION

### LBD37, LBD38, and LBD39 are nitrogen-induced repressors that suppress nitrogen responses

Previous studies have implicated LBD37, LBD38, and LBD39 in the negative regulation of N-responsive gene expression (Rubin et al. 2009; Albinsky et al. 2010; Brooks et al. 2019; Alvarez et al. 2020). Our analysis of *lbdT* mutants extends this view by demonstrating that these three LBDs act redundantly to suppress N responses under N-sufficient conditions. *lbdT* mutants showed derepression of nitrate- and N-starvation-responsive gene expression and enhanced N accumulation (Figures 2 to 5), effects opposite to those reported for LBD-overexpressing plants. Consistent with this, genes derepressed in *lbdT* significantly overlapped with genes downregulated in LBD-overexpressing plants (Figure S20; Table S20). Comparison with NIGT1s, established repressors of N responses (Kiba et al. 2018; Maeda et al. 2018), further highlighted the importance of LBDs within the negative regulatory network. Comparative transcriptome and phenotypic analyses using *lbdT*, *nigtQ*, and *nigtQlbdT* mutants revealed that, although LBDs and NIGT1s share both overlapping and distinct roles in the negative regulation of N responses, LBDs play a dominant role in this regulation (Figures 4 and 5).

### LBDs act as major repressors of CEP**–**CEPD-mediated systemic nitrogen-demand signaling

*CEP* and *CEPD* genes encode key inter-organ signaling components of systemic N-demand signaling (Tabata et al. 2014; Ohkubo et al. 2017; Ota et al. 2020). Our grafting and genetic analyses identify LBDs as major repressors of N-starvation-inducible *CEP* and *CEPD* genes and suggest that this repression gates the systemic N-demand signaling relay in both shoots and roots, allowing integration of N-status information from individual organs for determining a coordinated signaling output.

However, the molecular basis of this repression remains elusive. A simple direct promoter-repression mechanism is unlikely, because representative *CEP* and *CEPD* genes were not activated by LBD-VP fusion proteins and their promoters were not repressed by LBDs in co-transfection assays (Figures 7A and S17B). Since VP16 generally activates transcription most effectively when positioned near the transcription start site in plants (Waki et al. 2012; Gong et al. 2020), LBDs may instead act at distal regions, such as within gene bodies or downstream of these genes. Alternatively, LBDs may repress these genes indirectly by acting through other factors. Previous studies have reported HOMOLOG OF BRASSINOSTEROID ENHANCED EXPRESSION2 INTERACTING WITH IBH1 (HBI1) and TEOSINTE BRANCHED1/CYCLOIDEA/PROLIFERATING CELL FACTOR120 (TCP20) to promote *CEP* expression by binding to the *CEP* loci (Chu et al. 2020). Cytokinins have been implicated in the induction of *CEPD1, CEPD2*, and *CEPDL2* (Ota et al. 2020; Taleski et al. 2023). In our transcriptome analysis, however, *HBI1*, *TCP20*, and immediate-early cytokinin-inducible type-A response regulator genes were not upregulated in the *lbdT* mutant. Thus, the upregulation of *CEP* and *CEPD* genes in *lbdT* cannot be readily explained by these known regulators, suggesting that LBDs may repress their expression through as-yet-unidentified regulatory mechanisms.

### LBDs repress nitrogen-responsive genes through two distinct pathways

In contrast to our results on *CEP* and *CEPD* genes, we found that several genes involved in nitrate uptake and assimilation, such as *NRT2.1*, *NRT2.5*, and *NIA1*, as well as the *LBD* genes themselves, were activated by LBD-VP fusion proteins, and their promoter activities were repressed by LBDs in co-transfection assays (Figures 7 and S18). These results suggest that LBDs repress N-responsive gene expression through at least two distinct regulatory pathways: (i) a promoter-associated pathway that likely acts at or near the promoters of nitrate uptake and assimilation genes and the *LBD* genes themselves to locally restrain nitrate uptake and assimilation and mediate autoregulation of *LBD* gene expression and (ii) a pathway that represses *CEP* and *CEPD* genes and gates the systemic N-demand signaling relay. Consistent with this two-pathway model, elevated *NRT2.1* expression and nitrate accumulation in *lbdT* were not fully suppressed by the *cepdT* mutations (Figure 6).

Regarding the promoter-associated pathway, neither the canonical LOB motif (Husbands et al. 2007) nor LOB motif-like sequences were required for LBD-mediated repression in the *NRT2.1*, *NRT2.5*, *NIA1*, and *LBD37* promoters (Figure S19). Instead, deletion and fragment analyses identified a 32-bp region of the *NIA1* promoter that was sufficient to confer LBD-mediated repression on a minimal promoter, and mutation of the central GGCC sequence abolished the repression conferred by this region (Figure 7). This region is similar to a previously identified binding motifs of class I LATERAL ORGAN BOUNDARIES DOMAIN proteins (characterized by two GC-rich blocks separated by an A/T-rich spacer), with the central GGCC forming one of the two GC-rich blocks (O’Malley et al., 2016). This region is also highly conserved among *Brassicaceae NIA1* promoters, supporting its potential functional significance (Figure S21). However, direct binding of LBDs to this sequence *in vitro* and association of LBDs with the endogenous *NIA1* locus *in vivo* remain to be demonstrated, and the consensus sequence recognized by LBDs is undetermined.

### Slow growth of *lbdT* mutants is associated with nitrogen-related metabolic imbalance and excessive CEP-CEPD-mediated signaling

Although N nutrition generally promotes plant growth, the *lbdT* mutants showed slow growth despite accumulating abundant N (Figure 2 and Figure S5). Slow growth was also observed under low-N conditions (Figure S22), indicating that this phenotype is not simply due to N excess.

One possible cause of this slow growth is disruption of N-related metabolic homeostasis as the severity of N-related metabolic perturbation paralleled the severity of the growth phenotype. N-related metabolite levels were more strongly altered in *nigtQlbdT* than in *lbdT,* whereas *nigtQ* showed only slight alterations (Figure 5). Consistent with this, *nigtQlbdT* showed slower growth than *lbdT,* while *nigtQ* alone did not display an obvious growth defect (Figures S13 and S14). Further analyses will be required to determine whether particular N-related metabolic changes underlie this slow-growth phenotype.

Alternatively (or additionally), the slow growth of *lbdT* may result from derepression of *CEP* and *CEPD* genes. CEPs and CEPDs have been reported to inhibit primary root growth, reduce lateral root density, and suppress shoot growth (Roberts et al. 2016; Jung et al. 2018; Huang et al. 2023; Taleski et al. 2023). CEPs have also been implicated in promoting plant immunity, raising the possibility that excessive CEP signaling may divert resources from growth to defense-related processes (Rzemieniewski et al. 2024). However, whether the growth-inhibitory effects of CEP and CEPD members are mediated through N-demand signaling and/or involve additional mechanisms beyond N regulation remains unclear. Together, our results suggest that LBDs are required for normal growth, potentially through maintaining N metabolic homeostasis and/or repressing *CEP* and *CEPD* genes.

In conclusion, we propose that LBD37, LBD38, and LBD39 act as transcriptional repressors that integrate local N status into the control of local and systemic N responses, thereby optimizing N acquisition and utilization at the whole-plant level. Under N-sufficient conditions in both shoots and roots, *LBD* expression is upregulated in both organs. LBDs then restrain N responses in an organ-autonomous manner through two regulatory pathways: (i) promoter-associated repression of N uptake and assimilation genes and (ii) repression of *CEP* and *CEPD* genes (Figure 8A). Through these pathways, LBDs limit excessive N acquisition and utilization. Conversely, under N-deficient conditions, *LBD* expression is downregulated in both shoots and roots, allowing activation of local nitrate uptake and assimilation pathways and systemic N-demand signaling (Figure 8B).

**Figure 8.**
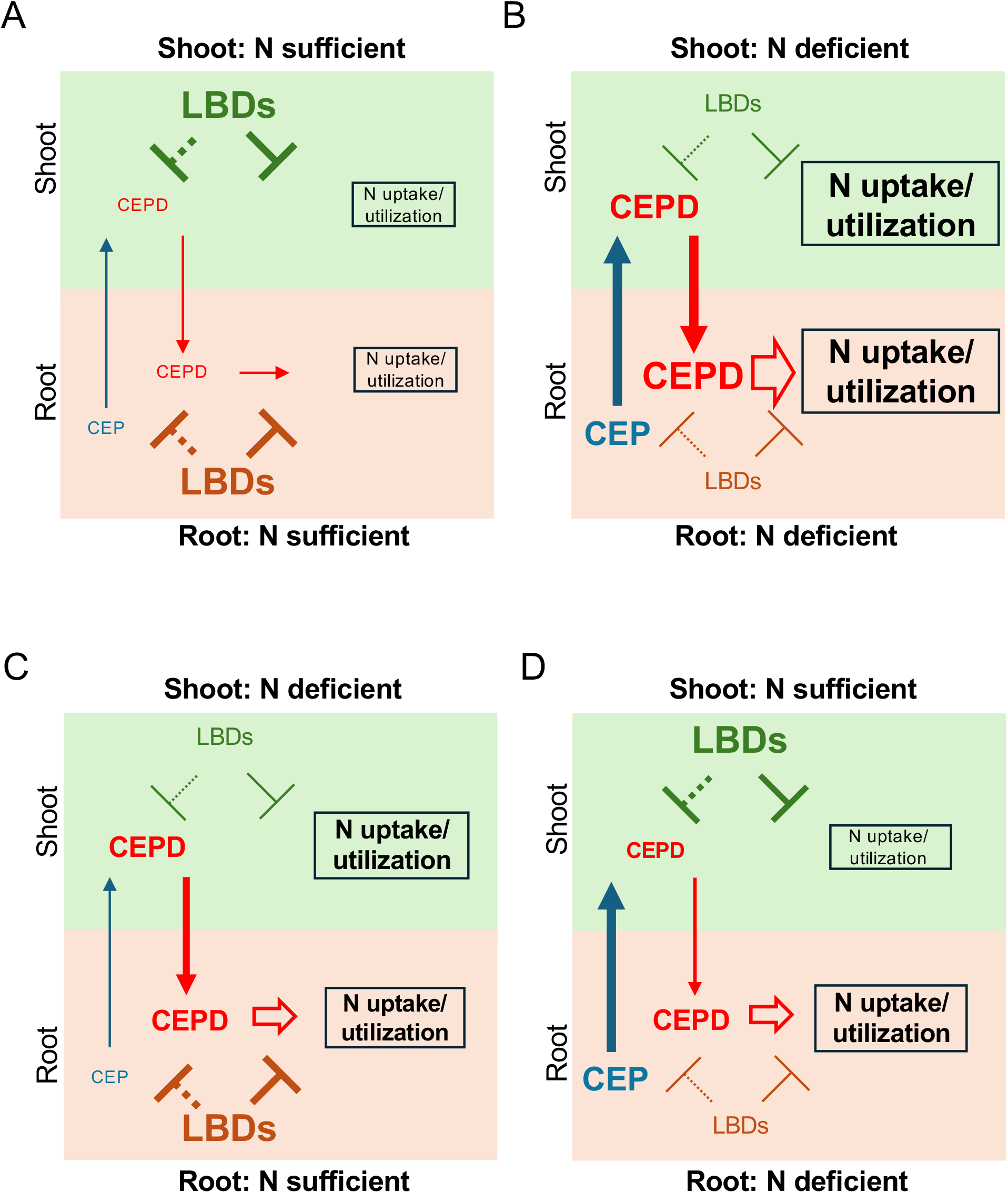
Model for LBD37-, LBD38-, and LBD39-mediated coordination of nitrogen acquisition and utilization at the whole-plant level in response to fluctuations in nitrogen availability and demand. LBD37, LBD38, and LBD39 (LBDs) act as N-induced repressors that restrain N responses through two regulatory pathways: promoter-associated repression of nitrate uptake and assimilation pathway genes and organ-autonomous repression of *CEP* and *CEPD* genes (dotted line). Four different situations are presented. (A) Plants are N-sufficient at the whole-plant level. This situation corresponds to the WT self-graft with respect to *LBD* expression. (B) Plants are N-deficient at the whole-plant level. This situation corresponds to the *lbdT* self-graft with respect to *LBD* expression. (C) N is available in the soil, but root N uptake is insufficient to satisfy shoot demand. This situation corresponds to the *lbdT*/WT graft with respect to *LBD* expression. (D) N availability decreases in roots while the shoot remains N-sufficient. This situation corresponds to the WT/*lbdT* graft with respect to *LBD* expression.

This model provides a framework for understanding the role of LBDs under spatially or developmentally changing N conditions. For example, when N is available in the soil but root N uptake is insufficient to satisfy shoot demand, shoot *LBD* expression would decline, relieving repression of *CEPD* expression and activating systemic N-demand signaling to enhance N uptake in roots (Figure 8C). In this situation, LBDs in roots would continue to repress nitrate uptake and assimilation genes to restrain the root response. When N availability decreases in roots while the shoot remains N-sufficient, *LBD* expression would decline in roots, relieving repression of *CEP* and *CEPD* genes as well as nitrate uptake and assimilation genes (Figure 8D). However, LBDs in the shoot would gate shoot-derived systemic N-demand signaling, limiting full activation of the systemic output. The reciprocal grafting experiments support these predictions. Root expression of *NRT2.1* and shoot nitrate accumulation increased in both *lbdT*/WT and WT/*lbdT* grafts compared with WT self-grafts but were highest in *lbdT* self-grafts (Figures 6D and 6E). *LBD37* and *LBD38* transcripts start accumulating rapidly (within 5 minutes) after nitrate addition (Brooks et al. 2019), whereas *LBD37*, *LBD38*, and *LBD39* transcript levels fall to less than half within 15 minutes after nitrate deprivation (Menz et al. 2016). These observations further suggest that rapid changes in expression would enable LBDs to dynamically adjust local and systemic N responses as N availability and demand fluctuate.

## METHODS

### Plant Materials and Growth Conditions

*Arabidopsis thaliana* accession Columbia-0 (Col-0) was used as the wild type. The Nössen accession was used as the wild type for experiments involving *cepd1-1 cepd2-1 cepdl2-1*, which are from Nössen background. T-DNA insertion lines GK-862G11 (*lbd37-3*), GK-049C12 (*lbd38-1)*, SALK_201956C (*lbd38-2)*, and SALK_044184 (*lbd39-2*) were obtained from the Arabidopsis Biological Resource Center. The *lbd37-3 lbd38-2 lbd39-2* (*lbdT-t1*) and *lbd37-3 lbd38-1 lbd39-2* (*lbdT-t2*) were generated by crossing these lines. The *nigt1.1-1* nigt1*.2-2 nigt1.3-1 nigt1.4-1* (*nigtQ-4*) and *cepd1-1 cepd2-1 cepdl2-1* have been described previously (Kiba et al. 2018; Ota et al. 2020). The *cepd1-1 cepd2-1 cepdl2-1* was a kind gift from Y. Matsubayashi (Nagoya University). The genotypes of T-DNA insertion lines were determined by genomic PCR using the primers shown in Table S21.

For studies on seedlings, surface-sterilized seeds were sown on half-strength Murashige and Skoog (0.5× MS)-based medium solidified with 1.1% agar and containing 1% sucrose and 2.6 mM MES (pH 5.8) in 140 × 100 mm rectangular plates (20 seeds/plate). Seedlings were grown vertically at 22°C under fluorescent light (50 µmol m^−2^ s^−1^, 16 h light/8 h dark). High-N-availability (+N) medium contained 10.3 mM NH_4_NO_3_ and 9.4 mM KNO_3_, whereas N-starved (−N) medium lacked both NH_4_NO_3_ and KNO_3_. To maintain ionic balance in −N medium, NH_4_NO_3_ and KNO_3_ were replaced with NaCl and KCl, respectively. A 0.5× MS-based medium containing 15 mM KNO_3_ as the sole N source was used where indicated. Adult plants were grown on nutrient-rich soil (Supermix A; Sakata) at 22°C under fluorescent light (100 µmol m^−2^ s^−1^, 16 h light/8 h dark).

### Quantitative Reverse Transcription PCR

Total RNA was isolated from plant tissues using NucleoSpin RNA (Macherey-Nagel). First-strand cDNA was synthesized from total RNA using the ReverTra Ace qPCR RT Master Mix (Toyobo). Quantitative PCR (qPCR) was performed on a QuantStudio 3 Real-Time PCR system (Thermo Fisher) using the KAPA SYBR Fast qPCR kit (KAPA Biosystems) and gene-specific primer sets (Table S21). Expression levels were normalized to *TIP41* (*At4g34270*) as an internal control (Kiba et al. 2018).

### Split-Root Experiment

Surface-sterilized seeds (20 seeds/plate) were sown on 140 × 100 mm rectangular plates containing +N medium solidified with 1.5% agar and grown vertically at 22°C under fluorescent light (50 µmol m^−2^ s^−1^, 16 h light/8 h dark). The primary root tip of 7-day-old seedlings was excised to enhance lateral root growth. Four days later, the primary root was severed just below the second lateral root, and the seedlings were cultured for an additional 5 days. Then, the seedlings were transferred to 9-cm-diameter, two-compartment Petri dishes containing a +N medium solidified with 1.5% agar on one side and a −N medium solidified with 1.5% agar on the other side, such that the two lateral root branches were separately placed in the two compartments. The seedlings were grown for an additional 3 days.

### GUS Staining

The *LBD37*, *LBD38*, and *LBD39* promoter fragments, comprising 2384, 2332, and 2357 bp upstream of the inferred initiation codons, respectively, were amplified by PCR with specific primers (Table S21) and cloned into the pENTR/D-TOPO vector (Invitrogen) to generate entry clones. The entry clones were recombined into the Gateway destination binary vector pBA002a-GUS (Kiba et al. 2018) to generate proLBD37:GUS, proLBD38:GUS, and proLBD39:GUS reporter constructs. The constructs were introduced into *Agrobacterium tumefaciens* strain EHA105, and Col-0 plants were transformed by the floral dip method (Clough and Bent 1998).

Histochemical GUS staining was performed as described previously (Kiba et al. 2018). Seedlings were vacuum-infiltrated for 5 minutes in staining buffer (50 mM potassium phosphate buffer, pH 7.0, 0.05% Triton X-100, 2 mM potassium ferrocyanide, 2 mM potassium ferricyanide, and 2 mM 5-bromo-4-chloro-3-indolyl β-D-glucuronide). Samples were incubated overnight at 37°C in the dark. Stained seedlings were cleared by incubation in a 70% to 100% ethanol series and mounted in a clearing solution consisting of chloral hydrate:glycerol:water (8:1:2, w/v/v) for observation by light microscopy (Axioplan 2; Zeiss).

### CRISPR-Cas9 Mutagenesis of *LBD37*, *LBD38*, and *LBD39*

Triple mutants of *LBD37*, *LBD38*, and *LBD39* were generated using a transfer RNA-based multiplex CRISPR-Cas9 vector, pMgPec12-137-2A-GFP (Hashimoto et al. 2018). Guide sequences were designed using CHOPCHOP (Labun et al., 2019), and the multiplex CRISPR-Cas9 vector pMgPec12-lbd37lbd38lbd39, harboring six guides, two for each *LBD* gene, was constructed as described previously (Kiba et al. 2023). The construct contained guide sequences g37-1, g38-1, g39-2, g37-2, g38-2, and g39-1 in this order. The construct was introduced into *Agrobacterium tumefaciens* strain EHA105, and Col-0 plants were transformed by the floral dip method (Clough and Bent 1998) to generate triple mutants, *lbd37-4 lbd38-3 lbd39-3* (*lbdT-c1*) and *lbd37-5 lbd38-4 lbd39-4* (*lbdT-c2*). Each line was obtained from an independent T1 plant. Mutations were identified by DNA sequencing of PCR products amplified from genomic DNA prepared from the corresponding lines using specific primer sets (Table S21). These lines were backcrossed to Col-0 to remove the CRISPR-Cas9 transgene before characterization. The septuple mutants, *nigtQlbdT-1* and *nigtQlbdT-2* were generated by crossing *nigtQ-4* with *lbdT-c1* or *lbdT-c2*. The double mutants, *lbd37-4 lbd38-3* (*lbd37 lbd38*), *lbd37-4 lbd39-3* (*lbd37 lbd39*), and *lbd38-3 lbd39-3* (*lbd38 lbd39*), were generated by backcrossing *lbdT-c1* with Col-0. The *cepd1-1 cepd2-1 cepdl2-1 lbdT-c1* (*cepdTlbdT*) sextuple mutant and *cepd1-1 cepd2-1 cepdl2-1 nigtQlbdT-1* (*cepdTnigtQlbdT*) decuple mutant were generated by crossing *cepd1-1 cepd2-1 cepdl2-1* with *lbdT-c1* or *nigtQlbdT-1*, respectively. The *cepdTlbdT* and *cepdTnigtQlbdT* lines were backcrossed twice to the corresponding parental lines, *lbdT-c1* or *nigtQlbdT-1*, before characterization.

### Growth Analysis

Rosette diameters, primary root lengths, and lateral root numbers were measured from digital images using ImageJ software (NIH, Bethesda, MD, USA). Flowering time was recorded as the number of days from sowing until the primary inflorescence stem reached 2 cm in length and the number of rosette leaves was counted at that time.

### Nitrate and Total N Measurements

For nitrate measurement, weighed shoots or roots of eleven-day-old seedlings were pulverized in liquid N and nitrate was extracted with 1 mL of ultrapure water. After centrifugation, the supernatant was collected and subjected to quantification by a colorimetric method at 540 nm as previously described (Miranda et al. 2001).

To determine total N, whole seedlings were dried to constant weight at 80°C and ground into fine powder. N content was determined using 1 mg of the powder with a CHN Corder MT-6 (Yanaco).

### Amino Acid Measurement

Eleven-day-old +N-grown seedlings were ground in liquid N and mixed with 10 volumes of 100 mM HCl containing 0.02 mM methionine sulfone as an internal standard. After centrifugation at 18,000g at 4°C for 5 minutes, the supernatant was derivatized using the derivatization reagent from the AccQ-Fluor Reagent Kit (Waters) following the manufacturer’s protocol. Derivatized amino acid levels were determined using a high-performance liquid chromatography system (Alliance 2695; Waters) linked to a photodiode array detector (2995; Waters) with AccQ-Tag eluents (Waters) and an AccQ-Tag Amino Acids C18 Column (3.9 mm × 150 mm; Waters) according to the manufacturer’s instructions.

### 15N Influx Measurement

Influx of ^15^NO_3_^−^ was assayed as previously described (Lezhneva et al. 2014) with some modifications. Seedlings were grown on +N agar plates for 11 days. Seedling roots were submerged in 0.1 mM CaSO_4_ for 1 minute and then transferred to 0.5× MS-based medium containing ^15^NO_3_^−^ (atom% ^15^N: 99%) at 0.2 mM as the sole N source for 10 minutes. Roots were transferred to 0.1 mM CaSO_4_ for 1 minute before harvest. Whole seedlings were dried to constant weight at 80°C and analyzed using a FlashEA1112 elemental analyzer (Thermo Fisher) and a DELTA V Advantage isotope-ratio mass spectrometer (Thermo Fisher) coupled via a ConFlo IV interface (Thermo Fisher).

### Co-transfection Assays in Arabidopsis Mesophyll Protoplasts

For mesophyll protoplast preparation, Col-0 plants were grown on nutrient-rich soil (Supermix A; Sakata) at 22°C under fluorescent light (100 µmol m^-2^ s^-1^, 12 h light/12 h dark) for 25 days. Mesophyll protoplast preparation and co-transfection were performed as described previously (Wu et al. 2009).

For transcriptional repressor activity assays (Figure S8), protoplasts were co-transfected with a reporter plasmid harboring pGAL4:FLUC (GAL4-responsive promoter:*firefly luciferase*), an effector plasmid expressing LBD37, LBD38, or LBD39 fused with the GBD at its N terminus under the CaMV 35S promoter (35Spro), and a reference plasmid expressing *Renilla luciferase* (*RLUC*) gene under the 35Spro (Fujimoto et al. 2000; Nakamichi et al. 2010). To construct effector plasmids, the coding region of LBD37, LBD38, or LBD39 including the stop codon was amplified by RT-PCR with specific primers (Table S21) and cloned into the pENTR/D-TOPO vector (Invitrogen) to obtain entry clones, pENTR-LBD37, pENTR-LBD38, and pENTR-LBD39. The entry clones were recombined into the Gateway destination vector pBS-GAL4DB-GW (Nakamichi et al. 2010) using Gateway LR Clonase II (Invitrogen), generating GBD-LBD37, GBD-LBD38, and GBD-LBD39. FLUC and RLUC activities were measured using the Dual-Luciferase Reporter Assay System (Promega) and TriStar2 LB942 (Berthold Technologies). Promoter activity was calculated as the FLUC/RLUC ratio.

For analysis of the effects of LBDs on promoter activity (Figures 7C to 7G, S17B, S18C, and S19), protoplasts were co-transfected with a reporter plasmid in which the promoter of interest drives *NanoLuc* (*NLUC*), an effector plasmid expressing LBD37, LBD38, or LBD39 under the 35Spro, and a reference plasmid expressing *FLUC* under 35Spro. To construct effector plasmids, pENTR-LBD37, pENTR-LBD38, and pENTR-LBD39 were recombined into the Gateway destination vector pBS-35S-GW using Gateway LR Clonase II (Invitrogen). pBS-35S-GW was generated from pBS-GAL4DB-GW by removal of the GAL4 DNA-binding domain (GAL4DB) cassette. Reporter plasmids were generated by cloning PCR-amplified promoter fragments (Table S21) into p35S-Nluc (Taoka et al. 2021) between the *Hin*dIII and *Nco*I sites, replacing the 35S promoter, or into p35Smin-NLUC between the *Hin*dIII and *Eco*RV sites, using the NEBuilder HiFi DNA Assembly Master Mix (New England Biolabs). p35Smin-NLUC contains a minimal 35S promoter (35Smin; −67 to +1) and was derived from p35S-Nluc by truncation of the 35S promoter, with the *Hin*dIII and *Eco*RV sites located upstream of 35Smin. Mutant promoter constructs were generated by PCR amplification of promoter fragments using primers carrying the desired mutations (Table S21), followed by simultaneous assembly of the resulting fragments and cloning into p35S-Nluc or p35Smin-NLUC using the NEBuilder HiFi DNA Assembly Master Mix (New England Biolabs). NLUC and FLUC activities were measured using the Nano-Glo Luciferase Assay System (Promega) and TriStar2 LB942 (Berthold Technologies). Promoter activity was calculated as the NLUC/FLUC ratio.

### RNA-seq Analysis

Total RNA was extracted from shoots and roots of 11-day-old Col-0, *nigtQ-4*, *lbdT-c1*, and *nigtQlbdT-1* seedlings grown on +N agar plates, as well as from shoots and roots of Col-0 seedlings grown on +N agar plates for 8 days and transferred to −N agar plates for 3 days, using NucleoSpin RNA (Macherey-Nagel). RNA-seq libraries were prepared using the NEBNext Poly(A) mRNA Magnetic Isolation Module (New England Biolabs) and NEBNext Ultra II RNA Library Prep Kit for Illumina (New England Biolabs) according to the manufacturer’s protocol. The libraries were sequenced on a NextSeq 550 (Illumina) using a NextSeq 500/550 High Output Kit v2 (Illumina) with single-end 81-bp reads.

RNA-seq read quality was assessed with FastQC. Reads were then mapped to the Arabidopsis reference genome (TAIR10) using HISAT2. Reads assigned to genes were counted using HTSeq, and normalization, low-count filtering, and differential expression analysis were performed using the edgeR Bioconductor package (Robinson et al. 2010). Each mutant genotype was independently compared with +N-grown Col-0. To identify N-starvation-inducible genes, −N-grown Col-0 was compared with +N-grown Col-0. DEGs were defined as genes with an absolute log2 fold change > 1, FDR < 0.1, and P < 0.05. KEGG pathway enrichment and GO term enrichment analyses were performed using ShinyGO v0.85.1 (Ge et al. 2020), with KEGG and Gene Ontology Biological Process selected as the pathway databases, respectively, and default settings for all other parameters.

### Widely Targeted Metabolome Analysis

Seedlings grown on +N agar plates for 11 days were lyophilized, and 4 mg of powdered tissue was extracted with 1 mL of extraction solvent consisting of 80% methanol (v/v) and 0.1% formic acid (v/v). Two internal standards were used: 8.4 nM lidocaine (positive ion mode) and 210 nM 10-camphorsulfonic acid (negative ion mode). The extracts were diluted to 1 mg mL^−1^ using the extraction solvent. Then, 25 µL of the extract was transferred to a 96-well plate, dried, reconstituted in 250 µL of ultrapure water, and 60 µL of the reconstituted extract was filtered using a MultiScreen HTS 384-well filter plate (MZHVN0W50; Merck Millipore). One microliter of the filtrate was analyzed using an ultra-high-performance liquid chromatography (UHPLC)– triple quadrupole mass spectrometry system (Nexera X2/LCMS-8050; Shimadzu) with an octadecylsilyl column (ACQUITY UPLC HSS T3, 1.8 µm, 1 mm × 50 mm; Waters). A metabolomic data matrix containing 474 metabolite intensities was obtained using optimized selected reaction monitoring (SRM) conditions and retention times (Sawada et al. 2009, 2017).

Peaks were manually inspected to confirm correct peak assignment and integration, and peaks with a height <100 or with retention-time deviations exceeding ±0.03 min were removed. Metabolites were excluded if the ratio of the peak intensity to that of the extraction solvent control was ≤ 5 in two or more of four biological replicates in any of the six genotype groups. Metabolite annotations were further manually curated, and annotations considered biologically implausible for *Arabidopsis* based on known metabolite occurrence were excluded. After curation, 100 metabolites were retained. Peak intensities of the retained metabolites were normalized to internal standards, and the resulting data matrix was used for PCA. PCA was performed on the sample-by-metabolite matrix using the prcomp function in R with mean-centering and unit-variance scaling, and score plots were generated for PC1 and PC2 using ggplot2. Differential metabolite levels between genotypes were tested on log2-transformed data using limma, with P values adjusted by the Benjamini–Hochberg FDR method.

### Grafting Experiments

Surface-sterilized seeds (approximately 120 seeds/dish) were sown on 140 × 100 mm rectangular plates containing +N medium solidified with 0.5% gellan gum (1% sucrose, 2.6 mM MES, pH 5.7) and grown at 22°C under fluorescent light (25 µmol m^−2^ s^−1^, 16 h light/8 h dark). Five-day-old seedlings were cut transversely through the hypocotyl using the tip of an injection needle, and the scion and rootstock were aligned within 0.4-mm-diameter silicone microtubing (Monden et al. 2025, 2026). Grafted plants were incubated for 5 days at 27°C under continuous light (50 μmol m^−2^ s^−1^). Plants were transferred to +N agar plates and grown vertically for an additional 5 days at 22°C under fluorescent light (70 μmol m^−2^ s^−1^, 16 h light/8 h dark) before analysis.

### LBD-VP Inducible Expression Lines

The coding regions of *LBD37* and *LBD38* without stop codons were amplified by RT-PCR with specific primers (Table S21) and cloned into the pENTR/D-TOPO vector (Invitrogen) to obtain entry clones, pENTR-LBD37w/o and pENTR-LBD38w/o, respectively. The entry clones were recombined into the Gateway destination binary vector pER8-VP (Kiba et al. 2018) using Gateway LR Clonase II (Invitrogen), generating constructs that express LBD37 or LBD38 fused to two tandem VP16 transcriptional activation domains (VP) at their C termini under the control of the β-estradiol-inducible promoter. The constructs were transferred into *Agrobacterium tumefaciens* strain EHA105, and Col-0 plants were transformed by the floral dip method (Clough and Bent, 1998).

Seedlings from two independent lines each expressing LBD37-VP or LBD38-VP (37VP and 38VP, respectively) were grown on +N agar plates for 11 days, sprayed with solutions containing 10 μM β-estradiol or 0.1% ethanol (mock), and incubated for 12 hours.

### Statistical Analysis

Data are presented as means ± standard deviation (SD). Statistical analyses were performed using Prism (version 11.0.0; GraphPad), except for transcriptome and metabolome analyses, which were conducted using the methods described for each assay.

### Accession Numbers

Arabidopsis Genome Initiative locus identifiers for genes mentioned in this article are shown in Table S22. The RNA-seq data generated in this study have been deposited in the Gene Expression Omnibus under accession number GSE337997. Germplasm used in this study included: GK-862G11 (*lbd37-3*), GK-049C12 (*lbd38-1*), SALK_201956C (*lbd38-2*), SALK_044184 (*lbd39-2*), GK-267G03 (*nigt1.1-1*), SALK_070096 (*nigt1.2-2/hho2-2*), SAIL_28_D03 (*nigt1.3-1/hho1-1*), SALK_067074 (*nigt1.4-1/hrs1-1*), psi00303 (*cepd1-1 cepd2-1 cepdl2-1*).

## ACKNOWLEDGEMENTS

We thank the Arabidopsis Biological Resource Center for supplying T-DNA insertion lines. This work was supported by the Japan Society for the Promotion of Science (JSPS) KAKENHI Grant Numbers 20H02888 and 25K22344, the JSPS Program for Forming Japan’s Peak Research Universities (J-PEAKS) Grant Number JPJS00420230010, the Mishima Kaiun Memorial Foundation, the Sapporo Bioscience Foundation, and the Yakumo Foundation for Environmental Science to T.K.

## AUTHOR CONTRIBUTIONS

T.K., M.Y.H., S.Y., and H.S. designed the study, T.K., H.T., K.M., Y.S., K.K., M.S., F.B., and T.H. performed the experiments, T.K., H.T., K.M., Y.S., K.K., M.S., F.B., T.H., M.Y.H., S.Y., and H.S. analyzed the data, and T.K wrote the paper.

## Supplemental Data

**Figure S1.** GUS staining of proLBD37:GUS, proLBD38:GUS, and proLBD39:GUS seedlings exposed to high nitrogen availability or nitrogen starvation

**Figure S2.** *lbd37 lbd38 lbd39* triple mutants generated using the CRISPR-Cas9 system or by combining T-DNA insertion alleles

**Figure S3.** Deduced amino acid sequences of LBD37, LBD38, and LBD39 and their CRISPR-Cas9-generated mutant proteins

**Figure S4.** Root phenotypes and levels of individual amino acids of *lbd37 lbd38 lbd39* triple mutant seedlings grown under high nitrogen availability

**Figure S5.** Phenotypes of *lbd37 lbd38 lbd39* triple mutant plants grown in nutrient-rich soil

**Figure S6.** Phenotypes of *lbd37 lbd38*, *lbd37 lbd39*, and *lbd38 lbd39* double mutant seedlings grown under high nitrogen availability

**Figure S7.** Nitrate and amino acid contents of *lbd37 lbd38 lbd39* triple mutant seedlings grown with nitrate as the sole nitrogen source

**Figure S8.** Transcriptional repression activity assay of LBD37, LBD38, and LBD39 in Arabidopsis mesophyll protoplast co-transfection assays

**Figure S9.** Venn diagrams showing overlap between genes upregulated in *lbdT-c1* and downregulated in NIGT1.2ox

**Figure S10.** Expression levels of *LBD37*, *LBD38*, *and LBD39* in *nigtQ-4* and *NIGT1* family genes in *lbdT* mutants under high nitrogen availability

**Figure S11.** RT-qPCR analysis of additional representative LBD-repressed genes in Col-0, *nigtQ-4*, *lbdT* mutants, and *nigtQlbdT* mutants

**Figure S12.** Statistical evaluation of the expression levels of representative nitrate-inducible genes at individual time points in Figures 4B, 4C, and S11A

**Figure S13.** Additional phenotypes of *nigtQlbdT* mutants grown under high nitrogen availability

**Figure S14.** Phenotypes of *nigt1.1 nigt1.2 nigt1.3 nigt1.4 lbd37 lbd38 lbd39* septuple mutant plants grown in nutrient-rich soil

**Figure S15.** Nitrate and amino acid contents of *nigt1.1 nigt1.2 nigt1.3 nigt1.4 lbd37 lbd38 lbd39* septuple mutant seedlings grown with nitrate as the sole nitrogen source

**Figure S16.** Expression of *NRT2.1*, *NAR2.1*, and *CEPH* and nitrate content in the *cepd1 cepd2 cepdl2 nigtQ lbdT* decuple mutant grown under high nitrogen availability CR and normalized to *TIP41* as an internal control.

**Figure S17.** Additional data for characterization of LBD-mediated repression of *CEP* and *CEPD* genes and nitrate uptake and assimilation pathway genes

**Figure S18.** Analysis of transcriptional autoregulation among *LBD37*, *LBD38*, and *LBD39*

**Figure S19.** Effects of mutations in the LOB motif and LOB motif-like sequences on LBD38-mediated repression

**Figure S20.** Venn diagrams showing overlaps among genes downregulated in LBD37OX in LBD38OX, and LBD-repressed genes

**Figure S21.** Conservation of the 32-bp region in the *NIA1* promoter among *Brassicaceae* species

**Figure S22.** Growth of Col-0, *nigtQ-4*, *lbdT-c1*, and *nigtQlbdT* mutants under low nitrogen availability

**Table S1.** Upregulated genes in *lbdT-c1* shoots compared with Col-0 shoots

**Table S2.** Upregulated genes in *lbdT-c1* roots compared with Col-0 roots

**Table S3.** Significantly enriched KEGG pathways for genes upregulated in *lbdT-c1* shoots compared with Col-0 shoots

**Table S4.** Significantly enriched KEGG pathways for genes upregulated in *lbdT-c1* roots compared with Col-0 roots

**Table S5.** Upregulated genes in shoots of nitrogen-starved Col-0 seedlings compared with nitrogen-sufficient Col-0 seedlings

**Table S6.** Upregulated genes in roots of nitrogen-starved Col-0 seedlings compared with nitrogen-sufficient Col-0 seedlings

**Table S7.** List of primary nitrate-inducible genes from Liu et al. (2022).

**Table S8.** Statistical evaluation of overlaps between genes upregulated in *lbdT-c1* and nitrogen-starvation-inducible or primary nitrate-inducible genes. The significance of the overlap was assessed by a one-sided Fisher’s exact test using the genes testable in both analyses as background.

**Table S9.** List of genes in the overlapping regions of the Venn diagrams shown in Figure 3A

**Table S10.** List of genes in the overlapping regions of the Venn diagrams shown in Figure S9

**Table S11.** Statistical evaluation of overlaps between genes upregulated in *lbdT-c1* and genes downregulated in NIGT1.2ox

**Table S12.** Upregulated genes in *nigtQlbdT-1* shoots compared with *nigtQ-4* shoots (LBD-repressed genes in shoots)

**Table S13.** Upregulated genes in *nigtQlbdT-1* roots compared with *nigtQ-4* roots (LBD-repressed genes in roots)

**Table S14.** Upregulated genes in *nigtQlbdT-1* shoots compared with *lbdT-c1* shoots (NIGT1-repressed genes in shoots)

**Table S15.** Upregulated genes in *nigtQlbdT-1* roots compared with *lbdT-c1* roots (NIGT1-repressed genes in roots)

**Table S16.** List of genes in the overlapping regions of the Venn diagrams shown in Figure 4A.

**Table S17.** GO term enrichment analysis of LBD-preferentially repressed, NIGT1-preferentially repressed, and LBD/NIGT1 co-repressed gene sets in shoots and roots

**Table S18.** Relative peak area intensities and PCA parameters for compounds detected by widely targeted metabolome analysis

**Table S19.** Relative expression levels of *CEP* and *CEPD* genes used to generate the heat maps in Figures 6A, and statistical analysis of these values

**Table S20.** Statistical evaluation of overlaps between LBD-repressed genes and genes downregulated in LBD37OX and LBD38OX

**Table S21.** Primers used in this study

**Table S22.** Arabidopsis Genome Initiative locus identifiers of genes analyzed in this article

## Notes

### Competing Interest Statement

The authors have declared no competing interest.

