## SupplementaryFigures for "LBD-type transcription factors suppress local and systemic nitrogen responses through distinct regulatory pathways"

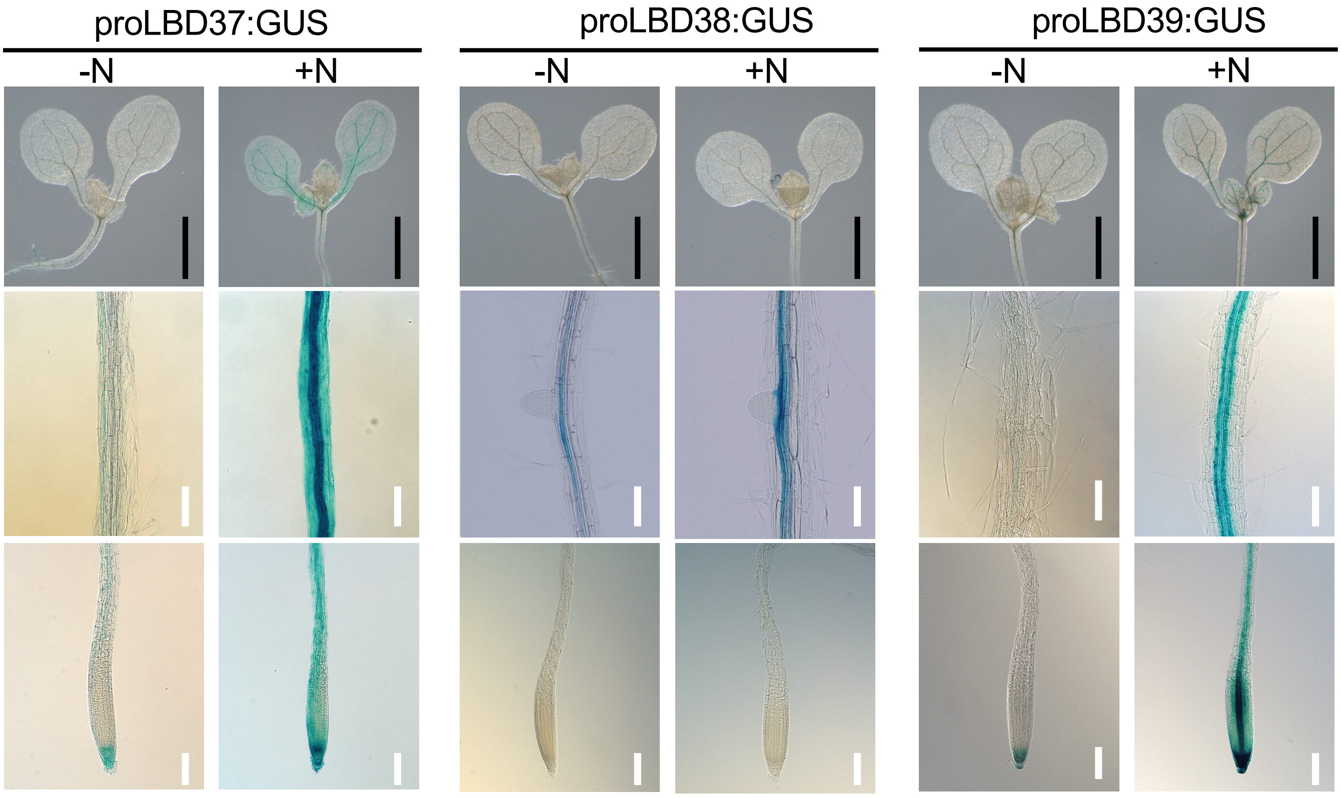


**Figure S1. GUS staining of proLBD37:GUS, proLBD38:GUS, and proLBD39:GUS seedlings exposed to high nitrogen availability or nitrogen starvation**

Eight-day-old N-starved seedlings were transferred to and incubated on 0.5× MS agar plates containing 10.3 mM $NH_4NO_3$ and 9.4 mM $KNO_3$ (+N) or no nitrogen (–N) for 24 hours before GUS staining. Black scale bars, 1 mm; white scale bars, 100 μm.

**A**

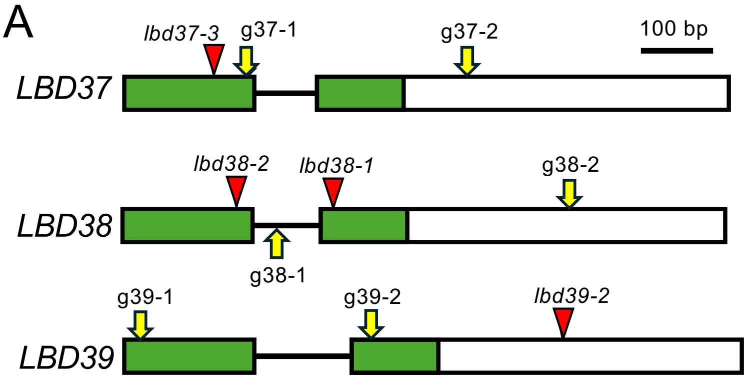


**B**

| Mutation | Description |
|---|---|
| *lbd37-4* | 1 bp insertion at g37-1 target site |
| *lbd37-5* | 1 bp insertion at g37-1 target site |
| *lbd38-3* | 466 bp deletion around g38-1 and g38-2 target sites |
| *lbd38-4* | 433bp deletion around g38-1 and g38-2 target sites |
| *lbd39-3* | 1 bp insertion at g39-1 target site |
| *lbd39-4* | 1 bp insertions at g39-1 and g39-2 target sites |

**C**

```
                            g37-1
LBD37    : 151 TTTATCTCCGCTG-TTCCGGATTCTCAACGTCCTGGTACTT 190
lbd37-4  :     TTTATCTCCGCTGATTCCGGATTCTCAACGTCCTGGTACTT
lbd37-5  :     TTTATCTCCGCTGTTTCCGGATTCTCAACGTCCTGGTACTT

                            g37-2
LBD37    : 450 TGGCTTTCCTTCGCCTACATCTGAAGAAGCATCTGAGATC 489
lbd37-4  :     TGGCTTTCCTTCGCCTACATCTGAAGAAGCATCTGAGATC
lbd37-5  :     TGGCTTTCCTTCGCCTACATCTGAAGAAGCATCTGAGATC
```

**D**

```
LBD38    : 126 CTTCGGCCGTGCCGGTTTAATGTCTTTCATCTCCGCCGTACCGGAATCTCAATGCCCTGGTACTT 195
lbd38-3  :     CTTCGGCCGTGCCGGTTTAATGTCTTTCATCTCCGCCGTACCGGAATCTCAATGCCCTGGTACTT
lbd38-4  :     CTTCG------------------------------------------------------------

                            g38-1
LBD38    : 196 AAAAACAAAAACCTCTCGGTCCTTTAGATCTTCGTCTA......CGATTCTCAAGCTCTAGATCT 580
lbd38-3  :     AAAAACAAAAA---------------------------......---------------------
lbd38-4  :     --------------------------------------......----TCTCAAGCTCTAGATCT

                                          g38-2
LBD38    : 581 AGATCAAGATCAACAGCTTCTCCGCCGAAACGGAAACGATTATCGTCGGAACAACAACCTTCGTC 645
lbd38-3  :     -----------------------------------------------------------------
lbd38-4  :     AGATCAAGATCAACAGCTTCTCCGCCGAAACGGAAACGATTATCGTCGGAACAACAACCTTCGTC

LBD38    : 646 GGAGCTTGATCTCTCTCTTATTCCTATTTATCCGATTAAAACCTTGCCGTTTAAG 700
lbd38-3  :     ----------------------CCTATTTATCCGATTAAAACCTTGCCGTTTAAG
lbd38-4  :     GGAGCTTGATCTCTCTCTTATTCCTATTTATCCGATTAAAACCTTGCCGTTTAAG
```

**E**

```
                               g39-1
LBD39    :   1 ATGAGTTGCAATGGATGTAGAGTTCTTCG-AAAAGGTTGCAGT  42
lbd39-3  :     ATGAGTTGCAATGGATGTAGAGTTCTTCGTAAAAGGTTGCAGT
lbd39-4  :     ATGAGTTGCAATGGATGTAGAGTTCTTCG-AAAAGGTTGCAGT

                            g39-2
LBD39    : 353 GTGGGAGAACGGTGAATCCGGTTAACG-GAGCGGTTGGTATGT 394
lbd39-3  :     GTGGGAGAACGGTGAATCCGGTTAACG-GAGCGGTTGGTATGT
lbd39-4  :     GTGGGAGAACGGTGAATCCGGTTAACGGGAGCGGTTGGTATGT
```

**F**

| Name | Description |
|---|---|
| *lbdT-t1* | *lbd37-3 lbd38-2 lbd39-2* |
| *lbdT-t2* | *lbd37-3 lbd38-1 lbd39-2* |
| *lbdT-c1* | *lbd37-4 lbd38-3 lbd39-3* |
| *lbdT-c2* | *lbd37-5 lbd38-4 lbd39-4* |

**Figure S2. *lbd37 lbd38 lbd39* triple mutants generated using the CRISPR-Cas9 system or by combining T-DNA insertion alleles**

**(A)** Schematic representation of CRISPR target sites and T-DNA insertion sites used to generate *lbd37 lbd38 lbd39* triple mutants. Boxes represent exons; horizontal bars, introns; red triangles, T-DNA insertion sites; yellow arrows, CRISPR target sites. Green boxes represent the DNA-binding domains of the LBD proteins.

**(B)** List of mutations in LBD genes generated using the CRISPR-Cas9 system.

**(C)** to **(E)** Partial sequences of wild-type and mutant alleles generated using the CRISPR-Cas9 system for *LBD37* (C), *LBD38* (D), and *LBD39* (E). The sequence corresponding to each guide RNA is highlighted in yellow. Numbers represent positions in the genomic sequence, with the first nucleotide of the putative start codon counted as 1. Red letters and red dashes indicate inserted and deleted nucleotides, respectively.

**(F)** Summary of *lbd37 lbd38 lbd39* triple mutants (*lbdT*) used in this study.

A

```
LBD37        1   MSCNGCRVLRKGCSENCILRPCIQWIETADAQGHATVFVAKFFGRAGLMSFISAVPDSQRPALFQ 65
lbd37-4          MSCNGCRVLRKGCSENCILRPCIQWIETADAQGHATVFVAKFFGRAGLMSFISAVSGFSTSCFVS
lbd37-5          MSCNGCRVLRKGCSENCILRPCIQWIETADAQGHATVFVAKFFGRAGLMSFISADSGFSTSCFVS
                 ****************************************************** . .   .:..

LBD37        66  SLLYEACGRTVNPVNGAIGMLWTGNWNICQAAVETVLRGGSLRPIPELLTHGGGFAGFPSPTSEE 130
lbd37-4          VVA------------------LRSLWKNCQSS*--------------------------------
lbd37-5          VVA------------------LRSLWKNCQSS*--------------------------------
                  :.                      *: **::
```

B

```
LBD38        1   MSCNGCRVLRKGCSENCILRPCIQWIESPEAQGHATVFVAKFFGRAGLMSFISAVPESQCPALFQ 65
lbd38-3          MSCNGCRVLRKGCSENCILRPCIQWIESPEAQGHATVFVAKFF----------------------
lbd38-4          MSCNGCRVLRKGCSENCILRPCIQWIESPEAQGHATVFVAKFFGRAGLMSFISAVPESQCPGT*-
                 *******************************************

LBD38        66  SLLYEACGRTVNPVNGAVGLLWTGNWNVCQAAVETVLRGGSLKPIPELLNGGGFAGFPSPTSDEA 130
lbd38-3          -----------------------------------------------------------------
lbd38-4          -----------------------------------------------------------------

LBD38        131 SEICTEMLNLRKADDSGDRNIYHHCRFSSSRSRSRSTASPPKRKRLSSEQQPSSELDLSLIPIYP 195
lbd38-3          --------------------------VSSSRSRSRSTASPPKRKRLSSEQQPSSELDLSLIPIYP
lbd38-4          -----------------------------------------------------------------

LBD38        196 IKTLPFKEDTPSMYSEESVTTVSFQNNNAGDRYVRCGGGGGGATTKLLNLFA* 247
lbd38-3          IKTLPFKEDTPSMYSEESVTTVSFQNNNAGDRYVRCGGGGGGATTKLLNLFA*
lbd38-4          ----------------------------------------------------
```

C

```
LBD39        1   MSCNGCRVLRKGCSETCILRPCLQWIESAESQGHATVFVAKFFGRAGLMSFISSVPELQRPALFQ 65
lbd39-3          MSCNGCRVLRKRLQ*--------------------------------------------------
lbd39-4          MSCNGCRVLRKGCSETCILRPCLQWIESAESQGHATVFVAKFFGRAGLMSFISSVPELQRPALFQ
                 ***********  .

LBD39        66  SLLFEACGRTVNPVNGAVGMLWTRNWHVCQAAVETVLRGGTLRPISDLLESPSLMISCDESSEIW 130
lbd39-3          -----------------------------------------------------------------
lbd39-4          SLLFEACGRTVNPVNGSGWYVVDQELARMPSGG*-------------------------------
```

**Figure S3. Deduced amino acid sequences of LBD37, LBD38, and LBD39 and their CRISPR-Cas9-generated mutant proteins**

Deduced amino acid sequences of LBD37 (A), LBD38 (B), and LBD39 (C) and the corresponding mutant proteins were aligned using Clustal Omega software (https://www.ebi.ac.uk/Tools/msa/clustalo/). The DNA-binding LBD domain is highlighted in green.

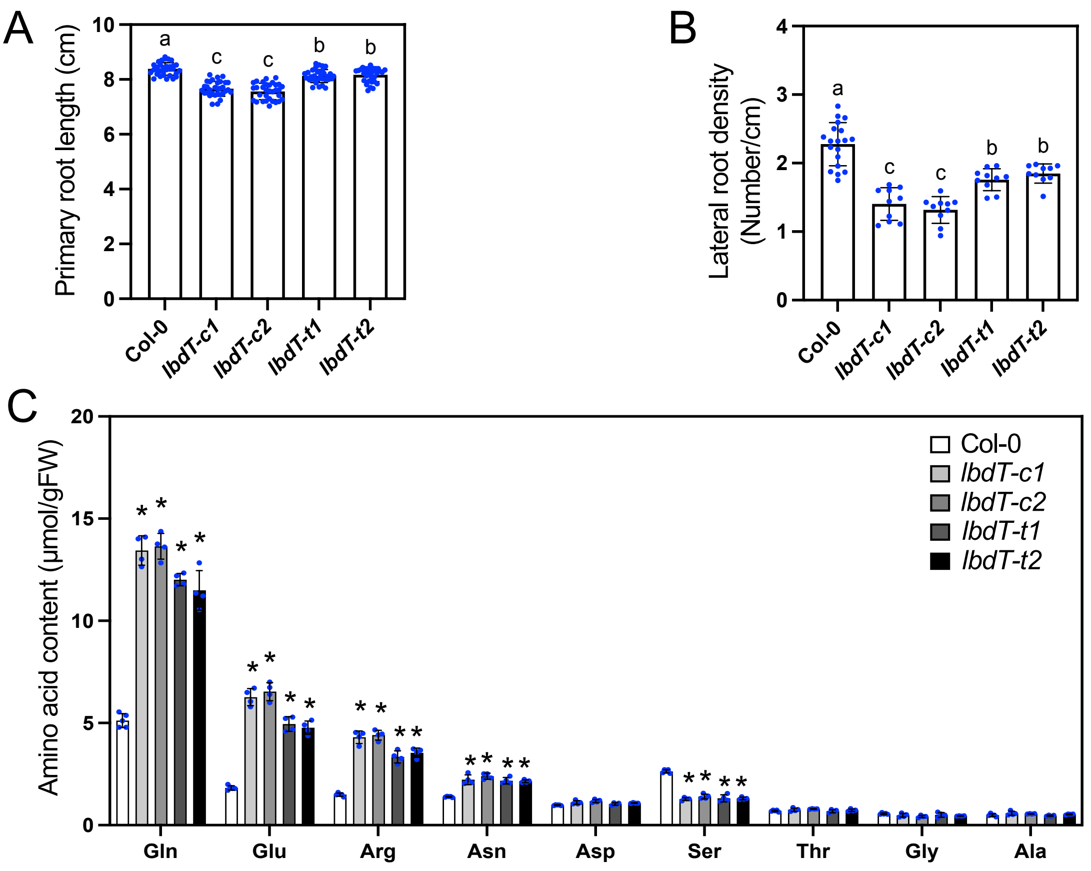


**Figure S4. Root phenotypes and levels of individual amino acids of *lbd37 lbd38 lbd39* triple mutant seedlings grown under high nitrogen availability**

Primary root length (A), lateral root density (B), and levels of individual free amino acids (C) of Col-0 (wild type) and *lbd37 lbd38 lbd39* triple mutants (*lbdT*) grown for 11 days on +N agar plates. Error bars represent SD (A, $n$ = 32; B, $n$ = 10–19; C, $n$ = 4 independent pools of the indicated tissue from 10–20 seedlings). Different lowercase letters indicate significant differences, as determined by Tukey's multiple comparisons test at $P < 0.05$. Asterisks indicate significant differences compared with Col-0, as evaluated by Student's $t$ test ($P < 0.05$).

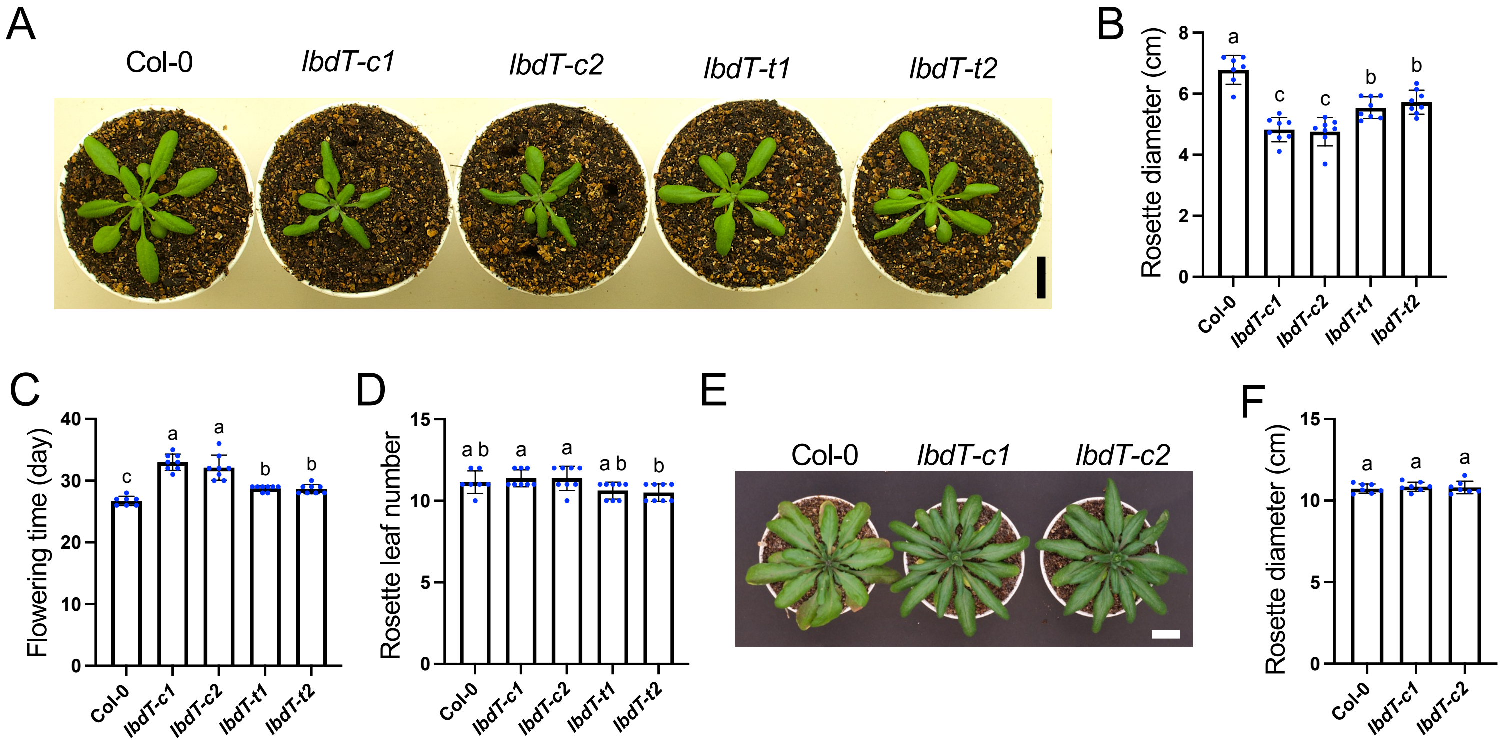


**Figure S5. Phenotypes of *lbd37 lbd38 lbd39* triple mutant plants grown in nutrient-rich soil**

Phenotypes of *lbd37 lbd38 lbd39* triple mutants grown in nutrient-rich soil. (A) Representative image and (B) rosette diameter at the vegetative stage (23-day-old). (C) Flowering time and (D) rosette leaf number at flowering. (E) Representative image and (F) rosette diameter at the reproductive stage (36-day-old). Col-0 and *lbd37 lbd38 lbd39* triple mutants (*lbdT*) were germinated and grown in nutrient-rich soil. Error bars represent SD (B to D, $n$ = 8; F, $n$ = 7). Different lowercase letters indicate significant differences, as determined by Tukey's multiple comparisons test at $P$ < 0.05. Scale bars, 2 cm.

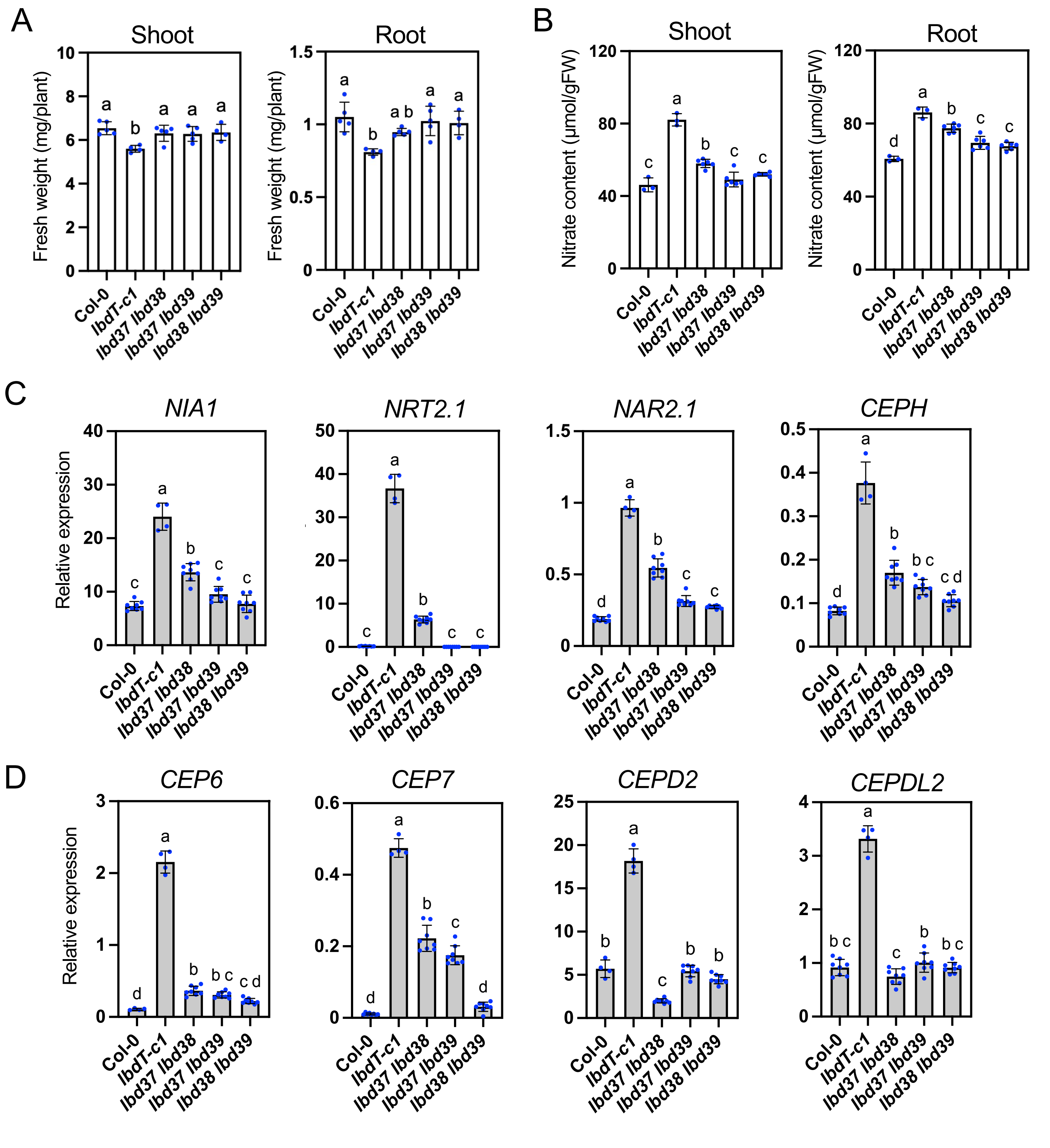


**Figure S6. Phenotypes of *lbd37 lbd38*, *lbd37 lbd39*, and *lbd38 lbd39* double mutant seedlings grown under high nitrogen availability**

Fresh weight (A), nitrate content (B), and gene expression (C, D) in Col-0, *lbdT-c1*, *lbd37-4 lbd38-3* (*lbd37 lbd38*), *lbd37-4 lbd39-3* (*lbd37 lbd39*), and *lbd38-3 lbd39-3* (*lbd38 lbd39*) seedlings grown for 11 days on +N agar plates. Genes shown in (C) are nitrate-inducible genes, either with or without N-starvation inducibility. Genes shown in (D) are induced only by N starvation. Error bars represent SD (A, $n$ = 4–5 independent pools of the indicated tissue from 13–15 seedlings; B, $n$ = 3–6 independent pools of the indicated tissue from 13–15 seedlings; C, D, $n$ = 4–8 independent pools of the indicated tissue from 10–20 seedlings). Different lowercase letters indicate significant differences, as evaluated by Tukey's multiple comparisons test at $P < 0.05$. Gene expression levels were analyzed by RT-qPCR and normalized to *TIP41* as an internal control.

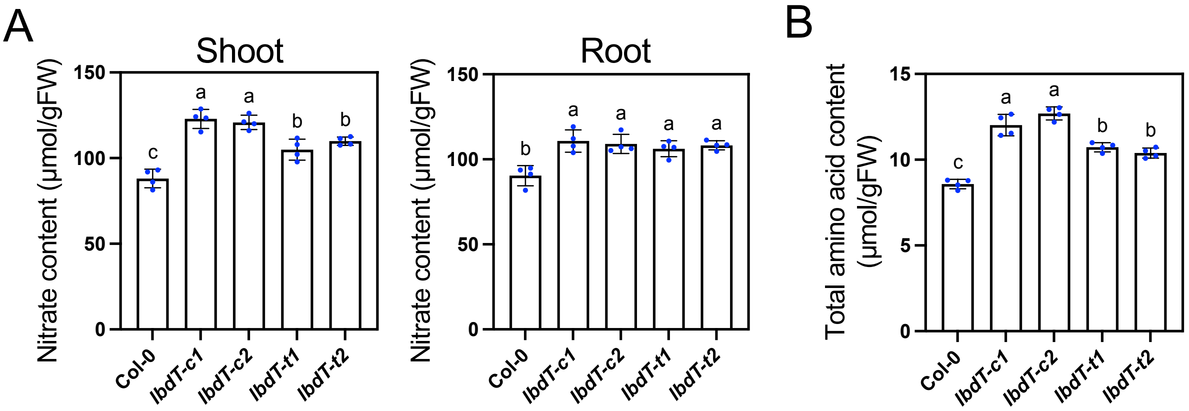


**Figure S7. Nitrate and amino acid contents of *lbd37 lbd38 lbd39* triple mutant seedlings grown with nitrate as the sole nitrogen source**

Nitrate content (A) and total free amino acid content (B) in Col-0 (wild type) and *lbd37 lbd38 lbd39* triple mutants (*lbdT*) grown for 11 days on 0.5× MS agar plates containing 15 mM nitrate as the sole N source. Error bars represent SD ($n$ = 4 independent pools of the indicated tissue from 10–20 seedlings). Different lowercase letters indicate significant differences, as determined by Tukey's multiple comparisons test at $P < 0.05$.

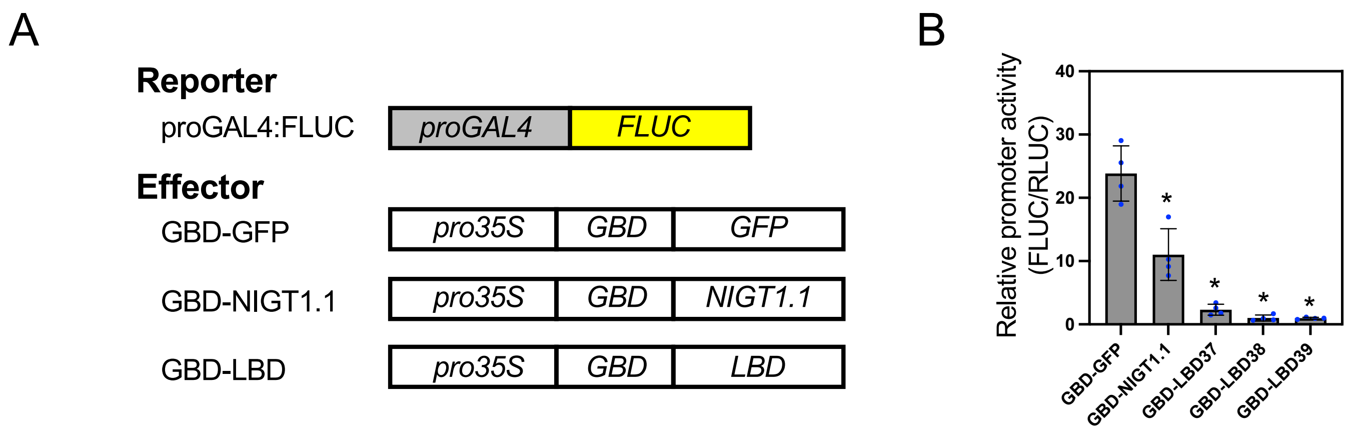


**Figure S8. Transcriptional repression activity assay of LBD37, LBD38, and LBD39 in Arabidopsis mesophyll protoplast co-transfection assays**

**(A)** Reporter, effector, and reference plasmids used in the assay. The reference plasmid harbors a *RLUC* gene driven by the CaMV 35S promoter (pro35S).

**(B)** GAL4 promoter (proGAL4) activity calculated as the ratio of FLUC activity to RLUC activity.

GBD, GAL4 DNA-binding domain. Error bars represent SD ($n$ = 4). Asterisks indicate statistically significant differences compared with the control (GBD-GFP) ($P < 0.05$, Student's $t$ test).

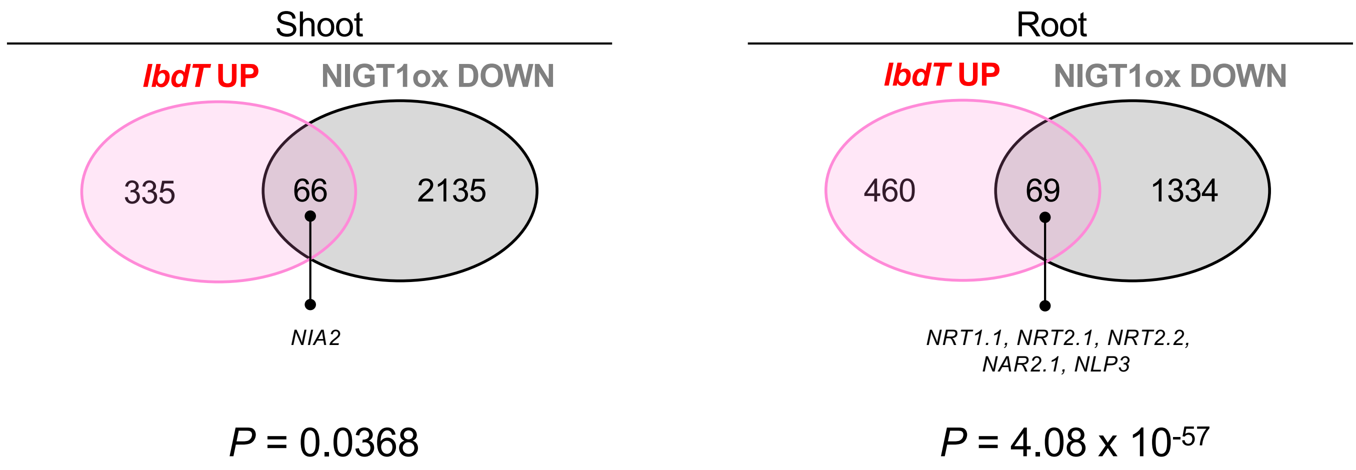


**Figure S9. Venn diagrams showing overlap between genes upregulated in *lbdT-c1* and downregulated in NIGT1.2ox**

Genes upregulated in *lbdT-c1* (*lbdT* UP; FC > 2, P < 0.05, FDR < 0.1) mutants and downregulated in NIGT1.2-overexpressing plants (NIGT1.2ox DOWN; Kiba et al. 2018) were compared. Representative N-responsive genes are presented. The significance of the overlap was assessed by a one-sided Fisher's exact test using genes testable in both analyses as the background. *P* values are shown below the diagrams (Table S11).

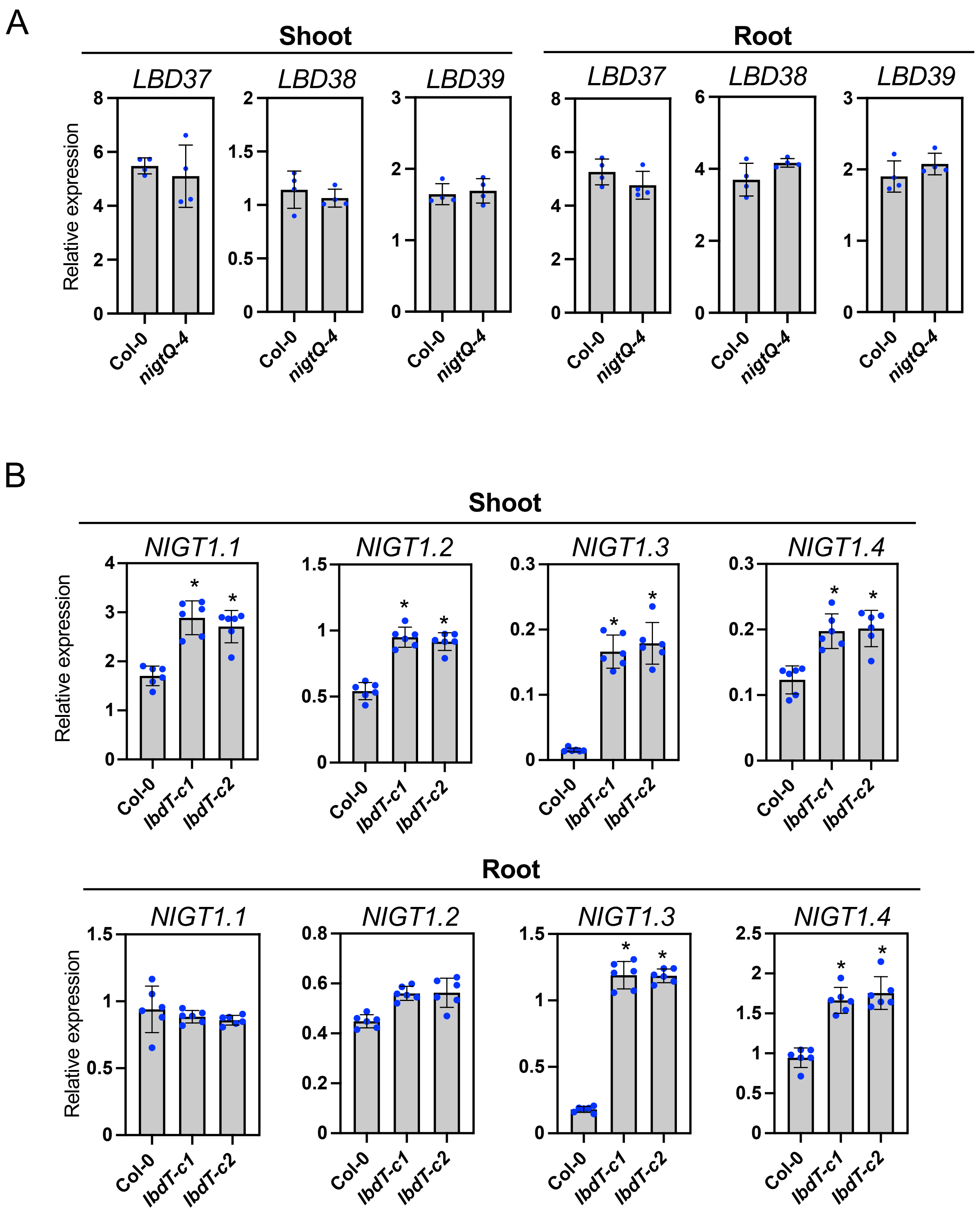


**Figure S10. Expression levels of *LBD37*, *LBD38*, *and LBD39* in *nigtQ-4* and *NIGT1* family genes in *lbdT* mutants under high nitrogen availability**

Expression levels of *LBD37*, *LBD38*, and *LBD39* in Col-0 and *nigtQ-4* (A) and *NIGT1* family genes in Col-0, *lbdT-c1*, and *lbdT-c2* (B) were analyzed in seedlings grown for 11 days on +N agar plates. Error bars represent SD (A, $n$ = 4 independent pools of the indicated tissue from 10–20 seedlings; B, $n$ = 6 independent pools of the indicated tissue from 10–20 seedlings). Asterisks indicate significant differences compared with Col-0, as evaluated by Student's $t$ test ($P < 0.05$). Gene expression levels were analyzed by RT-qPCR and normalized to *TIP41* as an internal control.

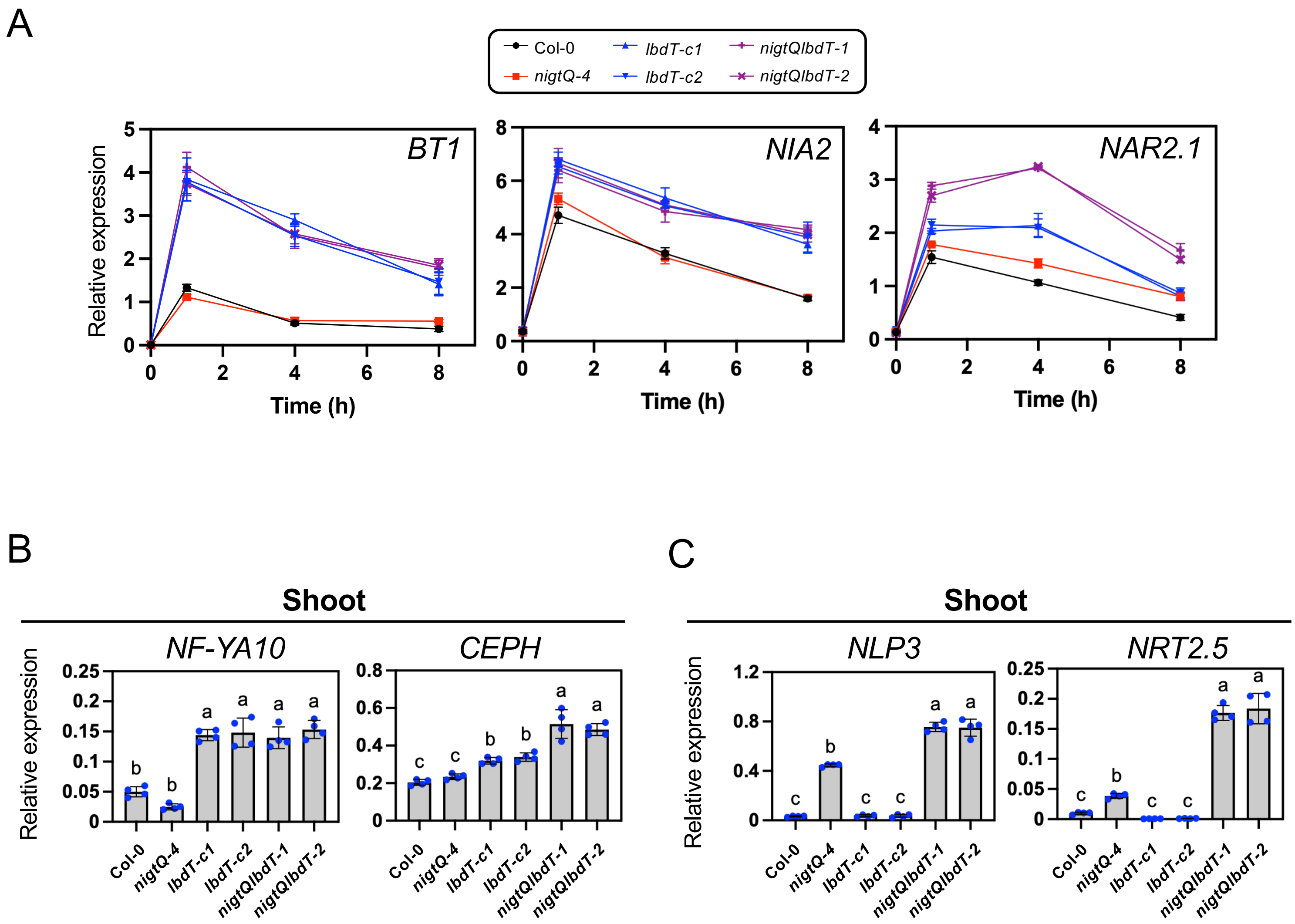


**Figure S11. RT-qPCR analysis of additional representative LBD-repressed genes in Col-0, *nigtQ-4*, *lbdT* mutants, and *nigtQlbdT* mutants**

**(A)** Time-course expression patterns of representative nitrate-inducible genes classified as non-overlapping LBD-repressed genes in Figure 4A. Seedlings of Col-0 (wild type), *nigtQ-4*, *lbdT* mutants (*lbdT-c1* and *lbdT-c2*), and *nigtQlbdT* mutants (*nigtQlbdT-1* and *nigtQlbdT-2*) were grown for 8 days in 0.5× MS liquid medium containing 0.5 mM ammonium succinate and then supplemented with 3 mM $KNO_3$.

**(B)** and **(C)** Expression levels in shoots of representative N-starvation-inducible genes classified in Figure 4A as non-overlapping LBD-repressed genes (B) or overlapping LBD/NIGT1-repressed genes (C). Col-0 (wild type), *nigtQ-4*, *lbdT*, and *nigtQlbdT* seedlings were grown for 11 days on +N agar plates.

Error bars represent SD ($n$ = 3-4 independent pools of the indicated tissue from 10–20 seedlings). Different lowercase letters indicate significant differences, as determined by Tukey's multiple comparisons test at $P$ < 0.05. Gene expression levels were normalized to *TIP41* as an internal control.

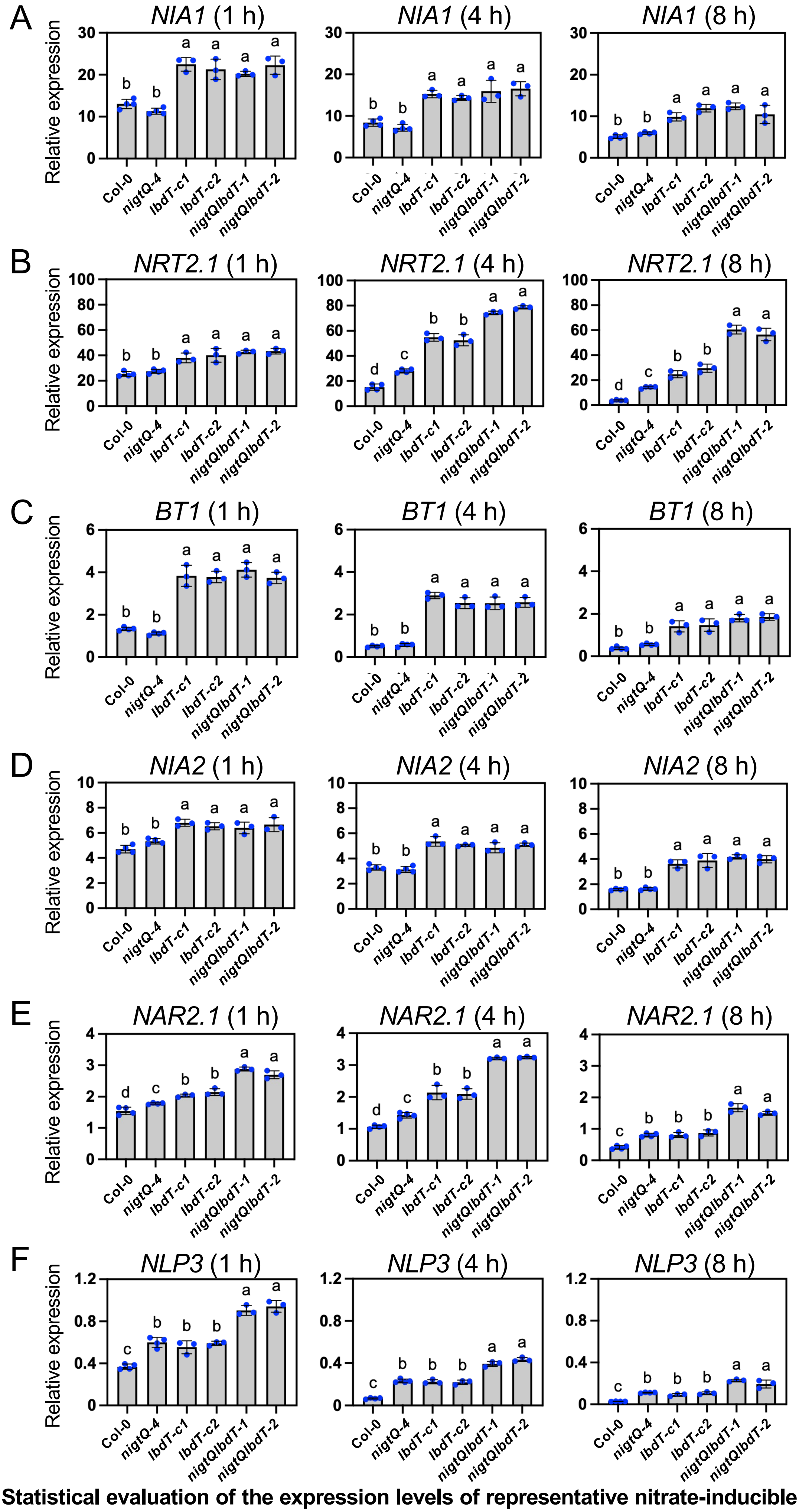


**Figure S12. Statistical evaluation of the expression levels of representative nitrate-inducible genes at individual time points in Figures 4B, 4C, and S11A**

Expression levels of *NIA1* (A), *NRT2.1* (B), *BT1* (C), *NIA2* (D), *NAR2.1* (E), and *NLP3* (F) shown as time-course plots in Figures 4B, 4C, and S11A are replotted as bar graphs at 1 hour (h), 4 h, and 8 h after nitrate addition to allow statistical comparisons among genotypes at each time. Different lowercase letters indicate significant differences among genotypes at each time point, as determined by Tukey's multiple comparisons test at $P < 0.05$.

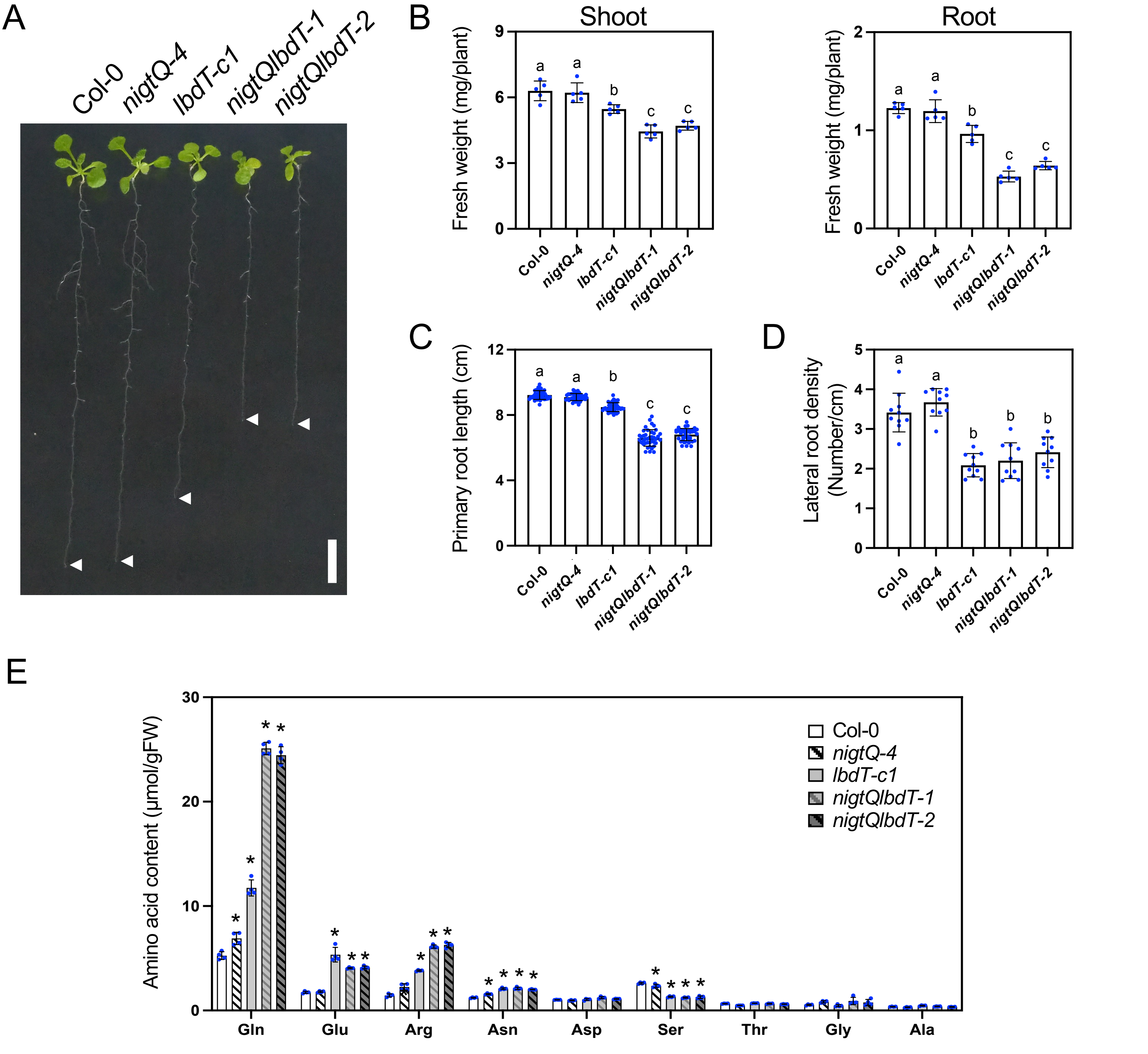


**Figure S13. Additional phenotypes of *nigtQlbdT* mutants grown under high nitrogen availability**

**(A)** to **(D)** Shoot and root growth of Col-0 (wild type), *nigtQ-4*, *lbdT-c1*, *nigtQlbdT-1*, and *nigtQlbdT-2* seedlings. Representative image (A), shoot and root fresh weights (B), primary root length (C), and lateral root density (D). Scale bar, 1 cm. Different lowercase letters indicate significant differences, as determined by Tukey's multiple comparisons test at $P < 0.05$.

**(E)** Levels of individual free amino acids in Col-0, *nigtQ-4*, *lbdT-c1*, *nigtQlbdT-1*, and *nigtQlbdT-2* seedlings.

Seedlings were grown for 11 days on +N agar plates. Error bars represent SD (B, $n$ = 5 independent pools of the indicated tissue from 10–20 seedlings; C, $n$ = 40; D, $n$ = 10; E, $n$ = 4 independent pools of the indicated tissue from 10–20 seedlings). Asterisks indicate significant differences compared with Col-0, as evaluated by Student's $t$ test ($P$ < 0.05).

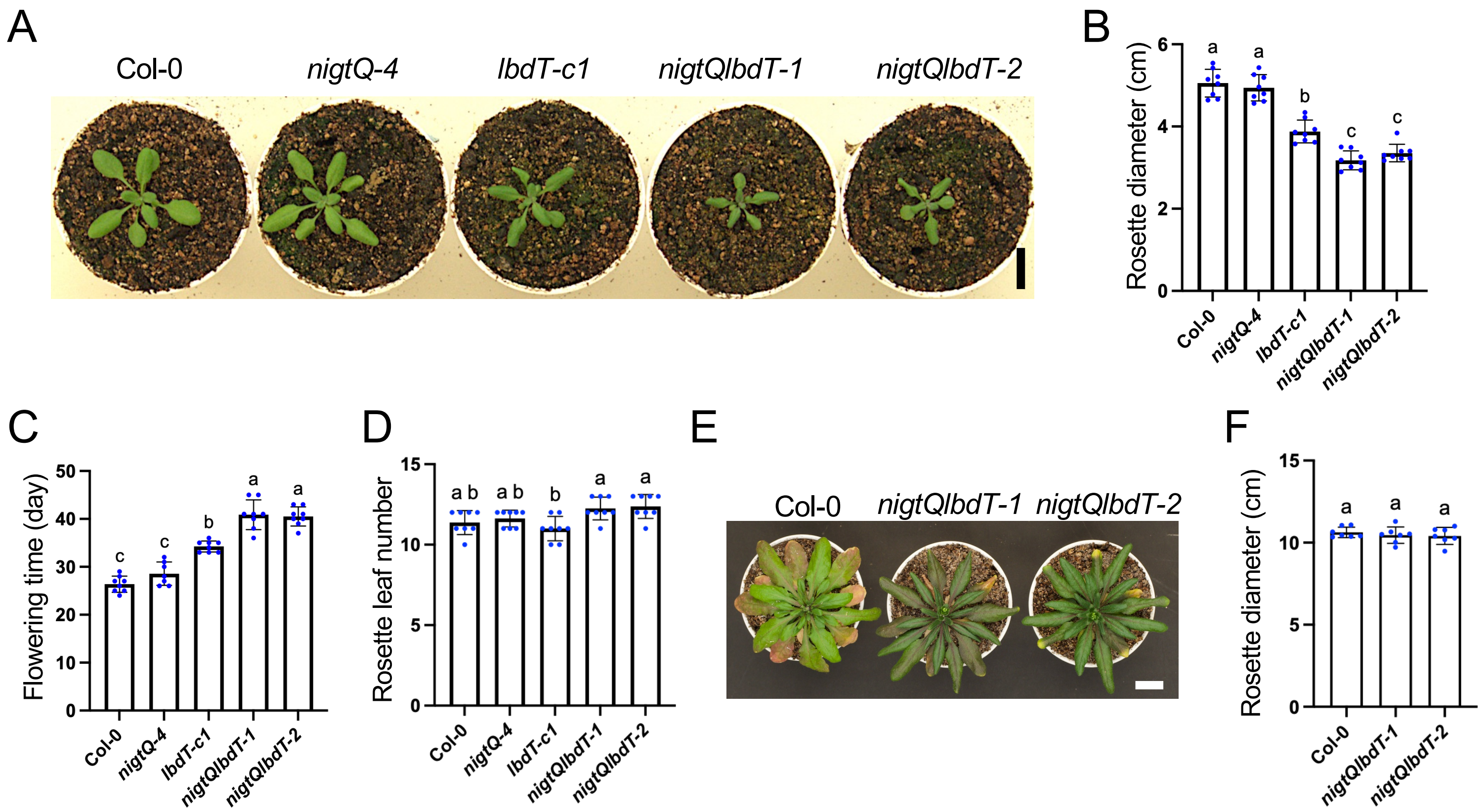


**Figure S14. Phenotypes of *nigt1.1 nigt1.2 nigt1.3 nigt1.4 lbd37 lbd38 lbd39* septuple mutant plants grown in nutrient-rich soil**

Phenotypes of *nigt1.1 nigt1.2 nigt1.3 nigt1.4 lbd37 lbd38 lbd39* septuple mutants grown in nutrient-rich soil. (A) Representative image and (B) rosette diameter at the vegetative stage (18-day-old). (C) Flowering time and (D) rosette leaf number at flowering. (E) Representative image and (F) rosette diameter at the reproductive stage (41-day-old). Col-0, *nigt1.1 nigt1.2 nigt1.3 nigt1.4* quadruple mutant (*nigtQ-4*), *lbd37 lbd38 lbd39* triple mutants (*lbdT-c1*), and *nigt1.1 nigt1.2 nigt1.3 nigt1.4 lbd37 lbd38 lbd39* septuple mutants (*nigtQlbdT-1* and *nigtQlbdT-2*) were germinated and grown in nutrient-rich soil. Error bars represent SD (B to D, $n$ = 8; F, $n$ = 7). Different lowercase letters indicate significant differences, as determined by Tukey's multiple comparisons test at $P < 0.05$. Scale bars, 2 cm.

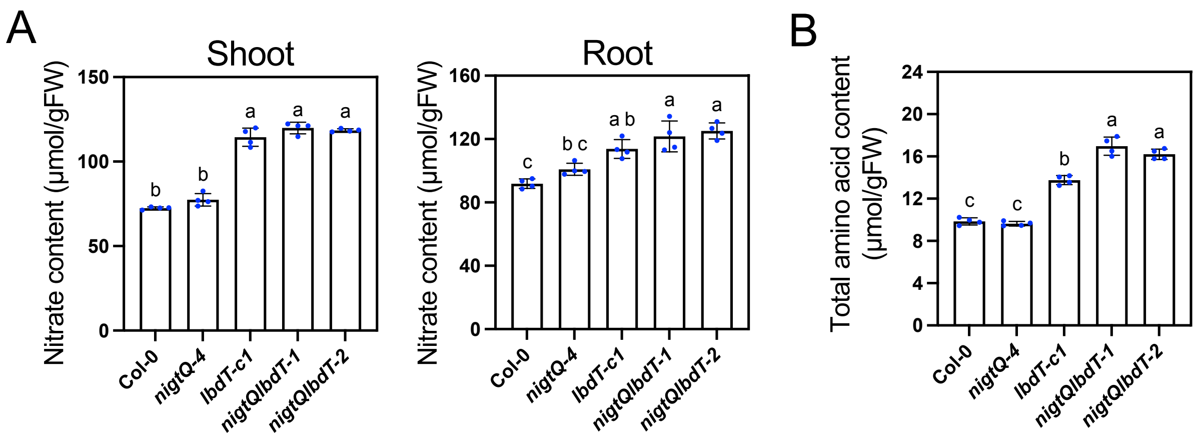


**Figure S15. Nitrate and amino acid contents of *nigt1.1 nigt1.2 nigt1.3 nigt1.4 lbd37 lbd38 lbd39* septuple mutant seedlings grown with nitrate as the sole nitrogen source**

Nitrate content (A) and total free amino acid content (B) in Col-0 (wild type), *nigtQ-4*, *lbd37 lbd38 lbd39* triple mutants (*lbdT-c1*), and *nigt1.1 nigt1.2 nigt1.3 nigt1.4 lbd37 lbd38 lbd39* septuple mutants (*nigtQlbdT*) grown for 11 days on 0.5× MS agar plates containing 15 mM nitrate as the sole N source. Error bars represent SD (*n* = 4 independent pools of the indicated tissue from 10–20 seedlings). Different lowercase letters indicate significant differences, as determined by Tukey's multiple comparisons test at $P < 0.05$.

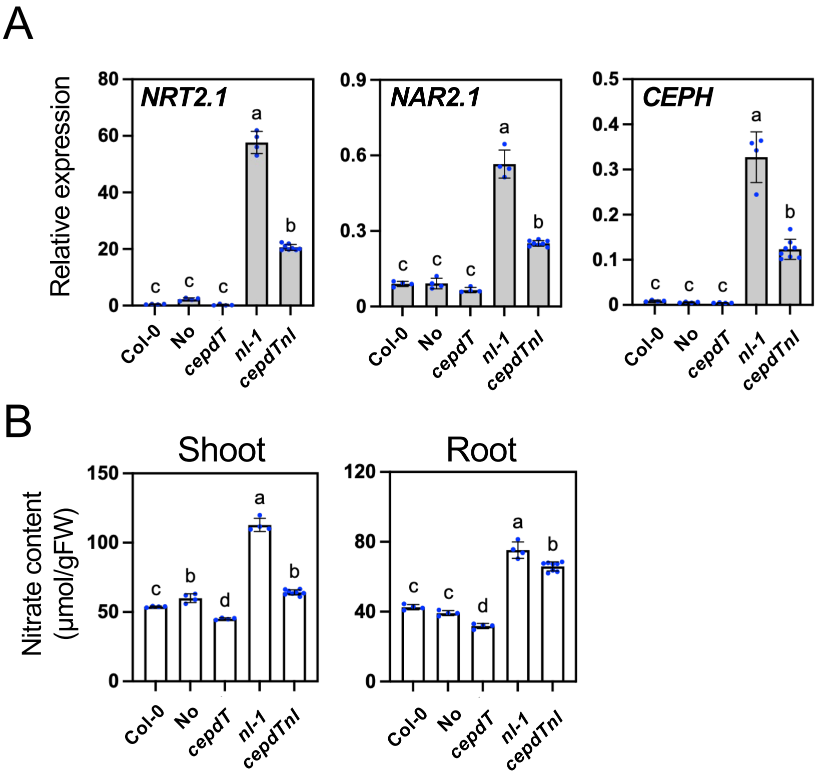


**Figure S16. Expression of *NRT2.1*, *NAR2.1*, and *CEPH* and nitrate content in the *cepd1 cepd2 cepdl2 nigtQ lbdT* decuple mutant grown under high nitrogen availability**

Expression levels of *NRT2.1*, *NAR2.1*, and *CEPH* in roots (A) and nitrate content in shoots and roots (B) in Col-0, Nössen (No), *cepd1 cepd2 cepdl2* (*cepdT*), *nigtQlbdT-1* (*nl-1*), and the *cepdT nl-1* decuple mutant (*cepdTnl*). Seedlings were grown for 11 days on +N agar plates. Because *cepdT* is in the No background, No was included as an additional wild-type control. Error bars represent SD ($n$ = 4–8 independent pools of the indicated organ from 10–20 seedlings). Different lowercase letters indicate significant differences, as determined by Tukey's multiple comparisons test at $P < 0.05$. Gene expression levels were analyzed by RT-qPCR and normalized to *TIP41* as an internal control.

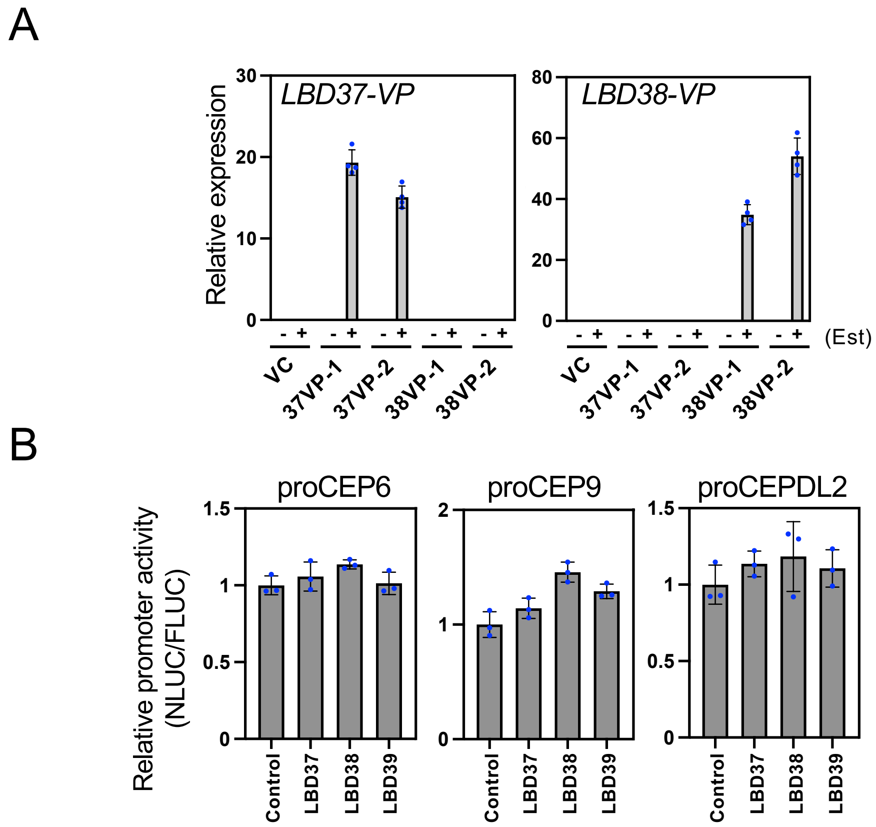


**Figure S17. Additional data for characterization of LBD-mediated repression of *CEP* and *CEPD* genes and nitrate uptake and assimilation pathway genes**

**(A)** Transcript levels of *LBD37-VP* and *LBD38-VP* in two independent transgenic lines (37VP and 38VP) and the empty-vector control (VC) used in Figures 7A, 7B, S15A, and S15B. Eleven-day-old seedlings grown on +N agar plates were incubated with (+) or without (−) 10 µM β-estradiol (Est) for 12 hours. Gene expression levels were analyzed by RT-qPCR and normalized to *TIP41* as an internal control.

**(B)** Effects of LBD37, LBD38, and LBD39 on the activities of the *CEP6, CEP9, and CEPDL2* promoters in co-transfection assays. Plasmids carrying the indicated promoters fused to *NLUC* were used as reporters. Plasmids expressing GUS (control), LBD37, LBD38, or LBD39 were used as effectors. A reference plasmid containing *FLUC* driven by the Cauliflower mosaic virus 35S promoter was used. Promoter activities are presented as NLUC/FLUC activity ratios.

Error bars represent SD (A, *n* = 4 independent pools of 10–20 seedlings; B, *n* = 3).

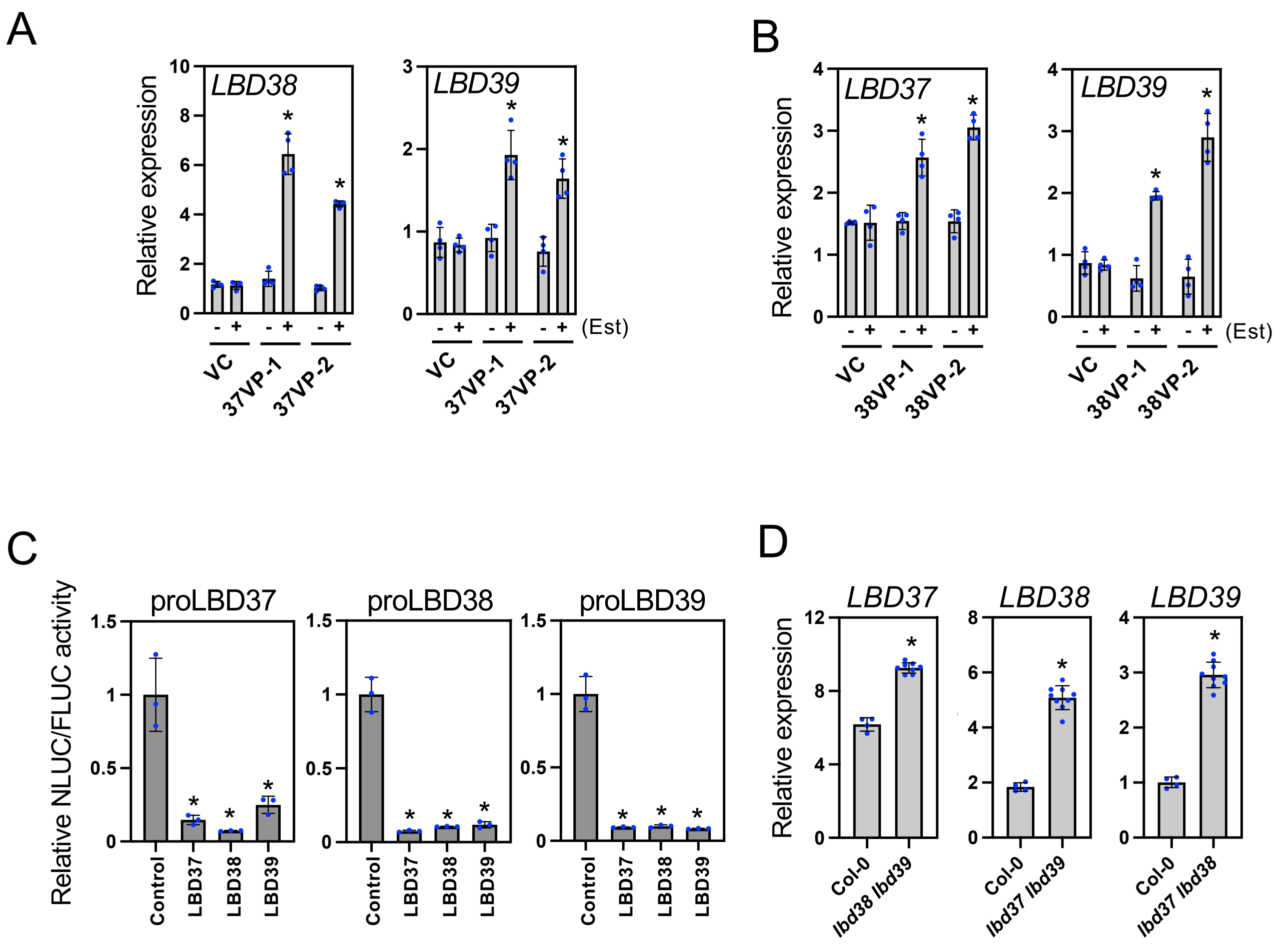


**Figure S18. Analysis of transcriptional autoregulation among *LBD37*, *LBD38*, and *LBD39***

**(A)** and **(B)** Expression of *LBD38* and *LBD39* in 37VP lines (A) and *LBD37* and *LBD39* in 38VP lines (B) grown on +N agar plates. VC indicates the empty-vector control. Eleven-day-old seedlings were incubated with (+) or without (−) 10 μM β-estradiol (Est) for 12 hours. Asterisks indicate significant differences between the −Est and +Est treatments ($P < 0.05$, Student's *t* test).

**(C)** Effects of LBD37, LBD38, and LBD39 on *LBD37*, *LBD38*, and *LBD39* promoter activities in co-transfection assays. Plasmids carrying the indicated promoters fused to *NLUC* were used as reporters. Plasmids expressing GUS (control), LBD37, LBD38, or LBD39 were used as effectors. A reference plasmid containing *FLUC* driven by the Cauliflower mosaic virus 35S promoter was used. Promoter activities are presented as NLUC/FLUC activity ratios. Asterisks indicate significant differences compared with the GUS control ($P < 0.05$, Student's *t* test).

**(D)** Expression of *LBD37*, *LBD38*, and *LBD39* in Col-0, *lbd37-4 lbd38-3* (*lbd37 lbd38*), *lbd37-4 lbd39-3* (*lbd37 lbd39*), and *lbd38-3 lbd39-3* (*lbd38 lbd39*) seedlings grown for 11 days on +N agar plates. Asterisks indicate significant differences compared with Col-0 ($P < 0.05$, Student's *t* test).

Expression levels were analyzed by RT-qPCR and normalized to *TIP41* as an internal control. Error bars represent SD (A and B, $n$ = 4 independent pools of 10–20 seedlings; C, $n$ = 3; D, $n$ = 4–8 independent pools of 10–20 seedlings).

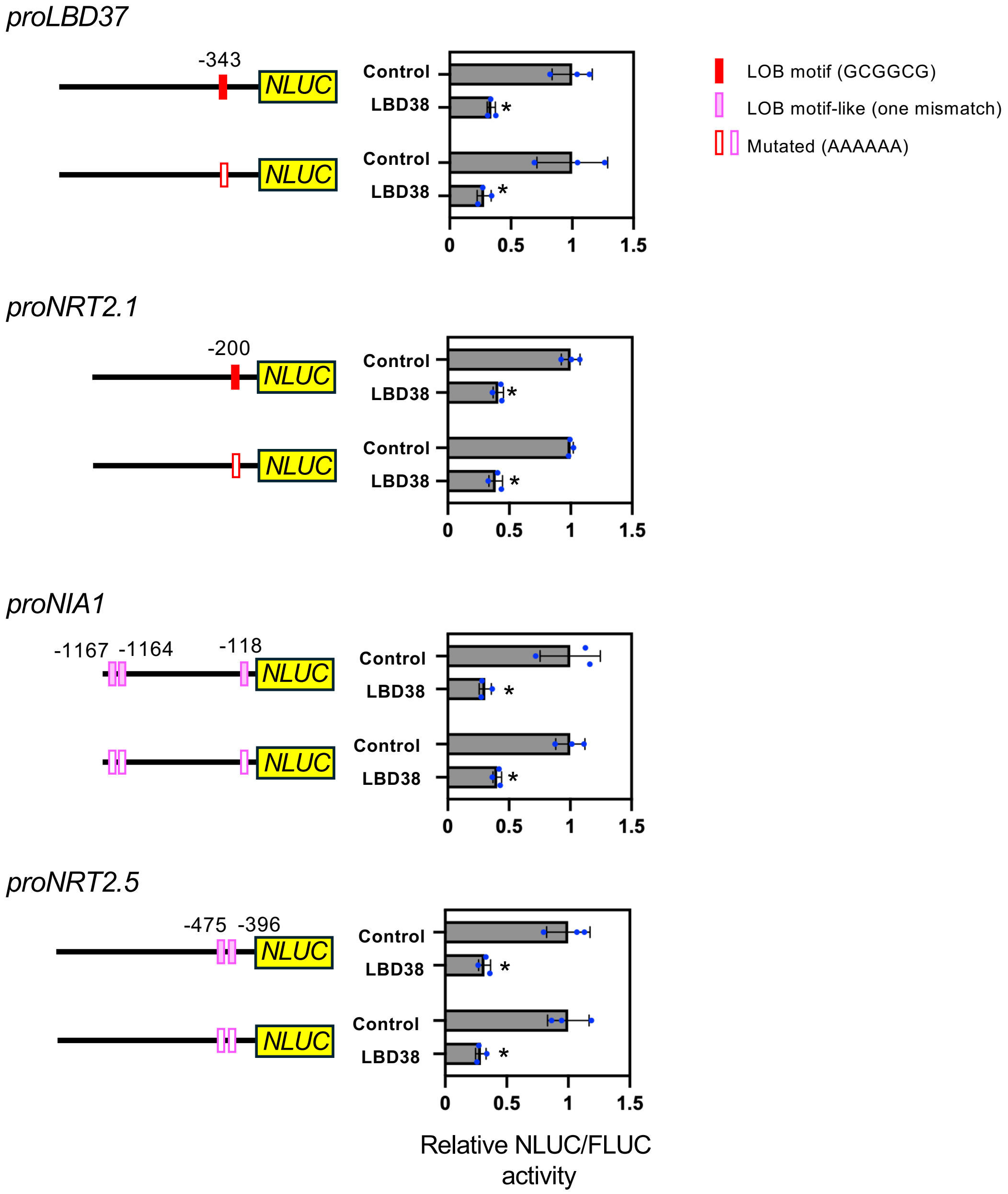


**Figure S19. Effects of mutations in the LOB motif and LOB motif-like sequences on LBD38-mediated repression**

Effects of LBD38 on the activities of native and mutated promoters of *LBD37*, *NRT2.1*, *NIA1*, and *NRT2.5* were examined in co-transfection assays. Reporter plasmids carried the indicated native promoters or their derivatives containing mutations in the previously reported LOB motif (GCGGCG) or in LOB motif-like sequences with a single-nucleotide mismatch, fused to *NLUC*. Plasmids expressing GUS (control) or LBD38 were used as effectors. A reference plasmid containing *FLUC* driven by the Cauliflower mosaic virus 35S promoter was used. Numbers denote the positions of the 5′ ends of the LOB motif or LOB motif-like sequences relative to the corresponding start codon. Promoter activities are presented as NLUC/FLUC activity ratios. Error bars represent SD ($n$ = 3). Asterisks indicate significant differences compared with the corresponding GUS control ($P$ < 0.05, Student's $t$ test).

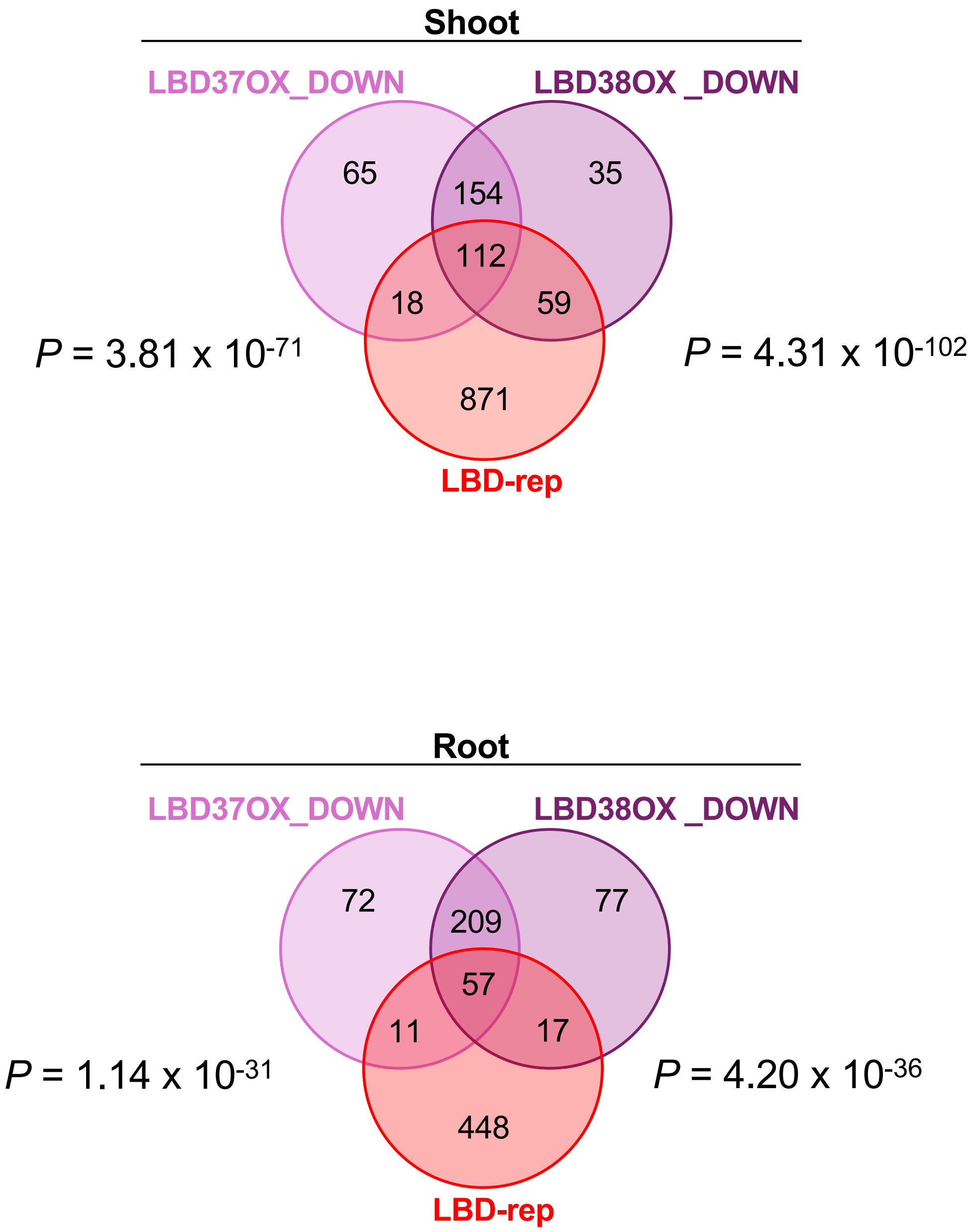


**Figure S20. Venn diagrams showing overlaps among genes downregulated in LBD37OX in LBD38OX, and LBD-repressed genes**

Overlaps among genes downregulated in LBD37OX and LBD38OX, and LBD-repressed genes in shoots and roots are shown. Genes downregulated in LBD37OX and LBD38OX were obtained from Rubin et al. (2009). The significance of the overlap was assessed by a one-sided Fisher's exact test using genes common to the compared datasets as the background. P values are also shown (Table S20).

```
BrNIA1      -----------------------------------------------GTTCGGTTGCTGG    13
CsNIA1      ------------------------------------------------------------    0
EsNIA1      -----------------------------------------ATCATAGTACCCTCCCACG    19
AlNIA1      -------------------CACACTACACAGACACATTGGCAACATAGT-----------    30
AtNIA1      ---------AAAGTTAAGTGTTGAGTGAGAGAGGTATCATAATCATAGTACCCTCCCACG    51
AhNIA1      ACTTTTCTTTGTTTTAAAGTTGTGTTGAGTGAGGTATCAATATCATAGTACCCTCCCACG    60
CrNIA1      ----------------------------------------TATCATAGTACCGTCCGAGG    20
BsNIA1      -------------------------------TTACTGAGGTATCATAGTACCCTCCTACA    29

BrNIA1      AT----------GCTCAATTATTTTTATAAACAAGTTATAATACTACAATCTTACTCTTT    63
CsNIA1      -TACTACCCACGGATCGAATCCTAAGATCCGCATTTTGTTTTA--CACA---T---AGTT    51
EsNIA1      GTACTGACCAGGGTTCGATTCCCTGGATGCGCATTTTTTTTAT--ATAT---G---TTTT    71
AlNIA1      -AACCTCCCACGGTTCGATTCCTTGATACGCATTTTTTAGTAT--TTTT---A---TCAT    81
AtNIA1      ATATTGCCCACGGATCGAATCCTTAGACCCGCATTTTATTTTT--AACT---T---CATA    103
AhNIA1      ATACTGCCCACGGATCGAATCCTTGAATCCGCATTTGGTTTTA--CCTT---A---ATTC    112
CrNIA1      ATACTGCCCACGGATCGAATCCTAGAATCCGAATTGTGGTAAT--TCGC---A---ATTC    72
BsNIA1      ATACTGCCCACGGATCGAATCCACAGATCCGCATTTTGTATTA--CCTT---A---TAAA    81
                        * ** * *

BrNIA1      TCTCCAAAAAAAACTACAATCTTACGAATATAATGATGTAGAAAAATCATAAATATTCCC    123
CsNIA1      TACATTTATTTAAGAATAAAAAAACATCATACAATACAACTGCGAAATAAAAAATCTCCC    111
EsNIA1      CTTCTTAAC--------------AAAAATAATTATATAATCTTACGTAATAAATTTTCCC    117
AlNIA1      ATTTTTGAC------ATCCGACAAAAACATATGTATATATATTACGAAATAAATGTTCCC    135
AtNIA1      TCGCATTAG------TATTTAAAAGCAAATATATATATCCTA-TGAAATAAAAATCTCCC    156
AhNIA1      GCA--TTAT------TTAAAAGAAA-AAAAAAACATATCTTACGAAATAAAAATCTTCCC    163
CrNIA1      ATTTAAAAA------ATAAA---AAATGTAAAATCGTGACCACAGAAATAAAAATCTCCC    123
BsNIA1      TTCGC----------ATTAT---TTAAAAGAAAATATATCTTACGAAATAAAAATCTCCC    128
                                                              ***   ****

BrNIA1      GCCTTAAAGGCCGTAGCTTAGGATCTAGGCA--GGTGGGTCTACCGACTCAAAGACCACA    181
CsNIA1      GCCTTAAAGGCCAAAGCTTGGGATCTAGGTTG-GTAGGTCCCTCGGACTCAGCGACCACA    170
EsNIA1      GCGTTAAAGGCCAAAGCTTAGGATCTAGGGGCCAGAGGGTCCCCCAACTTAGCGACCACA    177
AlNIA1      GCCATAAAGGCCAAAGCTTAGGATCTGGGCA--GGTGGGTCCCCCGACTCAGCGACCACA    193
AtNIA1      GCCTTAAAGGCCAAACCTTAGGATCTAGGTA--GGTGGGTCCCTCTACTCAGCGACCACA    214
AhNIA1      GCCTTAAAGGCCAAACCTTAGGATCTAGGTA--GGTGGGTCGCTCTACTCAGCGACCACA    221
CrNIA1      GCCTCAAAGGCCAAAGCTCAGGATCTAGCTA--GGTAGGTCCGTCGACCCGGCGACCACA    181
BsNIA1      GCCTTAAAGGCCAAAGCTTAGGATCTAGGTA--GGTACGTTCCTCGACTCAGTGACCACA    186
            **   *******  * **  ****** *                  **     *******

BrNIA1      CAAACTCACATTTGTCTATTTCTCCTCACATTTAATTTTTCATTATTTTATTTATTCATC    241
CsNIA1      GAAACTCACATTTGTCAATTCCTCCTTCCTCTCTTCTTTTTTCAATTTTATTTAT-----    225
EsNIA1      GAAAATCACATTTGTCAATTTCTTTTCCCTCTCATTTATACGTTTATTTTAATTGTTA--    235
AlNIA1      GAAACTCACAATTGTCAATTTCTCCTCCCTCTTGTTTTCAATTAATATATTATTAT----    249
AtNIA1      CAAACTCACATTTGTCAATTTCTCCTCTCTCTTAATTTATTAATTATTTTATTTA---CT    271
AhNIA1      CAAACTCACATTTGTCAATTTCTCCTCTCTCTTATTTTATTTATTTTTATTTCATTTATT    281
CrNIA1      TAAACTCACATTTGTCAATTTCTCCTCTCTCTTCTTTTTTTTGTTGCCTATTTTATTATT    241
BsNIA1      CAAACTCACATTTGTCAATTTCTCCTCTCTCTCTCTTATTTTTTTATTTTTAAA------    240
             *** ***** ***** *** **  *  *  *    *

BrNIA1      TT--------------------------GTAAATAACCCTATAAATACAACCCTATAACG    275
CsNIA1      ----AAACATTTATTTATTTACTAATCTCCGTAACGCCCTATAAATACAACCCTATAACT    281
EsNIA1      -------------TTTTATTATTCATCACTGTAACGGCCTATAAATACAACCCTATAACG    282
AlNIA1      -------------TTTATTTAATCTTCTCTGCAACGCCCTATAAATACAAGCCTATAACG    296
AtNIA1      CATCTCTGTAACGCCCCCTATAAATACAA-------------------------------    300
AhNIA1      CATCTCTGTAACGCTTATA-----------------------------------------    300
CrNIA1      TGTTTTGATTAAACTCATATCATCATATCTGTAACGCCCTATAAATACAACCCTATAAC-    300
BsNIA1      ----TTAAATTAATTTATTAACTCATCTCTGTAACGCCCTATAAATACAACCCTATAACG    296

BrNIA1      CCATATTATACACAAACCAAAGAAA 300
CsNIA1      CCTCATTATTCAGAAACCA------ 300
EsNIA1      CCTGCTTATTGACAAATC------- 300
AlNIA1      CCTG--------------------- 300
AtNIA1      ------------------------- 300
AhNIA1      ------------------------- 300
CrNIA1      ------------------------- 300
BsNIA1      CTTC--------------------- 300
```

**Figure S21. Conservation of the 32-bp region in the *NIA1* promoter among *Brassicaceae* species**

Promoter sequences spanning −400 to −100 bp relative to the *NIA1* start codon were obtained from Phytozome (https://phytozome-next.jgi.doe.gov/) and aligned using Clustal Omega software (https://www.ebi.ac.uk/Tools/msa/clustalo/). The analyzed *NIA1* genes were *AtNIA1* (*Arabidopsis thaliana*, AT1G77760), *AhNIA1* (*Arabidopsis halleri*, Ah2G29960), *CrNIA1* (*Capsella rubella*, Carub.0002s2466), *BsNIA1* (*Boechera stricta*, Bostr.20129s0213), *AlNIA1* (*Alyssum linifolium*, Alyli.0101s0038), *EsNIA1* (*Euclidium syriacum*, Eusyr.0119s0839), *BrNIA1* (*Brassica rapa* ssp. chinensis var. communis PCGlu, BrPCGlu.07G210900), and *CsNIA1* (*Camelina sativa* Acsn-226, CsAcsn226.07G248500). The 32-bp region mediating transcriptional repression in the *A. thaliana NIA1* promoter is highlighted, and the GGCC sequence whose substitution abolished LBD-mediated repression is shown in bold.

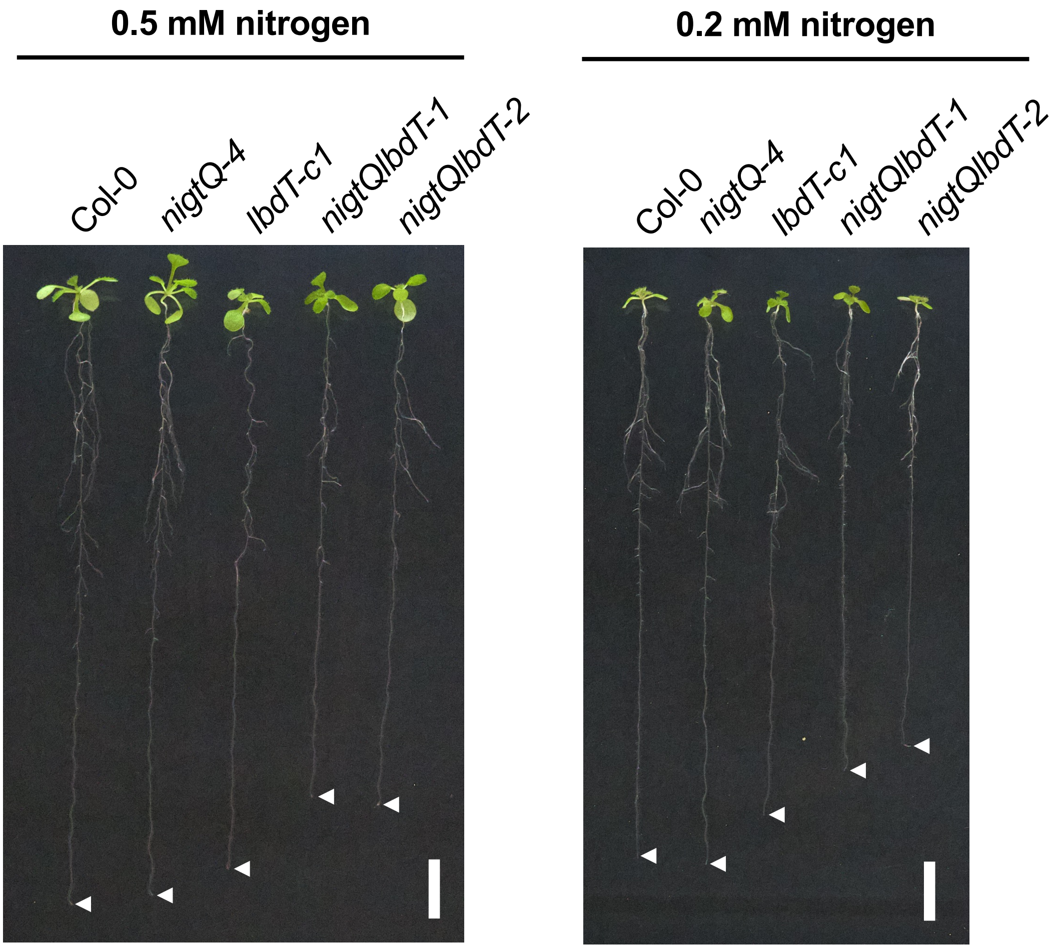


**Figure S22. Growth of Col-0, *nigtQ-4*, *lbdT-c1*, and *nigtQlbdT* mutants under low nitrogen availability**
A representative image of Col-0, *nigtQ-4*, *lbdT-c1*, *nigtQlbdT-1*, and *nigtQlbdT-2* seedlings grown for 11 days on 0.5 × MS agar plates containing either 0.17 mM $NH_4NO_3$ and 0.16 mM $KNO_3$ (0.5 mM nitrogen) or 0.07 mM $NH_4NO_3$ and 0.06 mM $KNO_3$ (0.2 mM nitrogen). Scale bar, 1 cm. White arrowhead indicates the position of primary root tip.
